# History of coadaptation to the *cag* pathogenicity island in *Helicobacter pylori*

**DOI:** 10.64898/2026.11.18.767480

**Authors:** Xiaoqi Zang, Sarah L. Svensson, Daniel Falush

## Abstract

Most *Helicobacter pylori* strains use genes encoded on the *cag* pathogenicity island (cagPAI) to inject the CagA oncoprotein into human cells, where it alters cell signalling and morphology to promote bacterial colonization. However, *cag^−^ H. pylori* strains are found worldwide. Here, we use a global sample of >9,000 *Helicobacter* genomes to investigate how strains adapt to *cag* status. We identified variants genome-wide associated with cagPAI presence, including many in *vacA* and *hopQ,* as well as in genes not previously linked to *cag.* We use these variants to construct a *cag* polygenic coadaptation score, allowing us to systematically explore the dynamics and evolutionary history of the island. Score distribution is bimodal in many populations, showing there is a stable adaptive strategy associated with *cag^−^* status. However, these *cag^−^* coadapted variants have historically been unavailable to strains in East Asia, providing a potential explanation for the low frequency of *cag^−^* strains in this region. We also infer that the cagPAI arrived in *H. pylori* before the human populations which colonized Australia and Papua New Guinea split from other groups 50,000–70,000 years ago. Adaptation to the island in other populations was achieved in part by using existing variants from the Hardy ecospecies. Our study illustrates how gene coadaptation shapes pathogen evolution after import of a major virulence factor and highlights genetic configurations that may predict increased or decreased disease risk, informing eradication strategies.

## Introduction

*Helicobacter pylori* achieves numerical dominance in the human stomach by persisting near epithelial cells and avoiding the highly acidic interior. Most strains carry the 40 kb *cag* pathogenicity island (cagPAI) which encodes a Type IV secretion system (T4SS) that injects the CagA effector protein and other molecules into gastric epithelial cells. CagA modulates cell signalling to alter cell polarity, cytoskeletal organization, proliferation, and inflammatory responses^1^. These changes promote persistent bacterial colonization and survival but can also induce gastric disease and cancer.Expression of the island is also energetically costly and the action of the cagPAI leads to strong IL-8 secretion and neutrophil recruitment that can be detrimental to bacterial survival. Loss of the island is a frequent outcome during infection and *cag^−^* strains persist at high frequency in some human populations^1^.

*H. pylori* recombines frequently during mixed infection, which has erased most signals of clonal descent between strains. Instead, it is subdivided into populations, which arose during the dispersal of humans around the world. Fifteen populations and subpopulations have been identified within the species^2^. Furthermore, the species is divided into Hardy and Ubiquitous ecospecies^3^, which likely arose in our ancestors before we became modern humans and have coexisted in our stomachs throughout our history as a species. Both ecospecies spread from Africa to Siberia, Oceania and the Americas and differ at around 100 genes, on an otherwise freely mixed genetic background. However, Hardy strains have gone extinct in most locations on their migration path and the ecospecies has only been found in a handful of Indigenous human groups and, following host jumps, in big cats and primates^3^.

The cagPAI is not found in other *Helicobacter* species that colonize the stomachs of other mammals^4^. However, according to current data, the hpAfrica2 population is the only fully cagPAI-negative (*cag^−^*) population^5^. hpAfrica2 is substantially diverged from all other *H. pylori* populations and is associated with Khoisan people in Africa, who represent the deepest splitting human population^6^. This pattern is consistent with the cagPAI being acquired in the ancestor of non-hpAfrica2 populations, before these populations spread with humans throughout Africa and globally, but it could also have spread from population to population at a later time.

Here, to investigate how strains adapt to the presence and absence of the island, we perform genome-wide association studies for *cag* status in the hpEurope and hspEAsia populations. This analysis generalizes previous inferences of genetic differences between cagPAI-positive (*cag^+^*) and *cag*^−^ bacteria such as in outer membrane proteins that facilitate cell binding and CagA injection (e.g., HopQ)^7^, as well as the vacuolating cytotoxin A (VacA)^8^, with 30 genome loci robustly associated with *cag* status in the HpEurope population.

We use the results of the GWAS to construct a polygenic coadaptation score and to investigate the dynamics of coadaptation in both ancient and recent times. We find that many variants associated with *cag^+^* status were originally imported from the Hardy ecospecies, and that the import of the island into *H. pylori* is most likely to have taken place shortly before the split of Indigenous Australians and Papua New Guinea (PNG) from other human populations, before the Sahul split, 50,000–70,000 years ago^9^. Our results shed light on how pathogens adapt to the presence of a new pathogenicity island and provide a framework to understand the distinct colonization strategies used by *H. pylori* to survive in the unique and challenging environment of the human stomach.

## Results

### GWAS identifies *cag* status associated variants

The complex genetic structure within our global collection of 9,164 genomes is visible in a phylogenetic tree **(Supplementary Fig. S1**, **Supplementary Table 1)**. To identify variants associated with carriage of the cagPAI within this highly structured dataset, we focused on two well sampled populations. HpEurope isolates have been isolated globally but form the majority in Europe, the Middle East, and the Americas. hspEAsia isolates make up a large majority of isolates in China, Japan, and Korea and are also a majority in other East Asian countries such as Vietnam, Thailand, Malaysia, and Singapore **(Supplementary Table 1)**. We assembled genomes with typical hpEurope and hspEAsia ancestry profiles, consisting of 1,425 and 1,179 isolates, respectively (**Fig. 1b, c**). These two subsets of the data have less structure than the global dataset, so that their phylogenies are more star-like. However, there are still visible subdivisions, with most Japanese isolates branching together in the hspEAsia population **(Fig. 1c)** and three previously described sub-populations clustering within the hpEurope tree **(Fig. 1b)**.

**Fig. 1.**
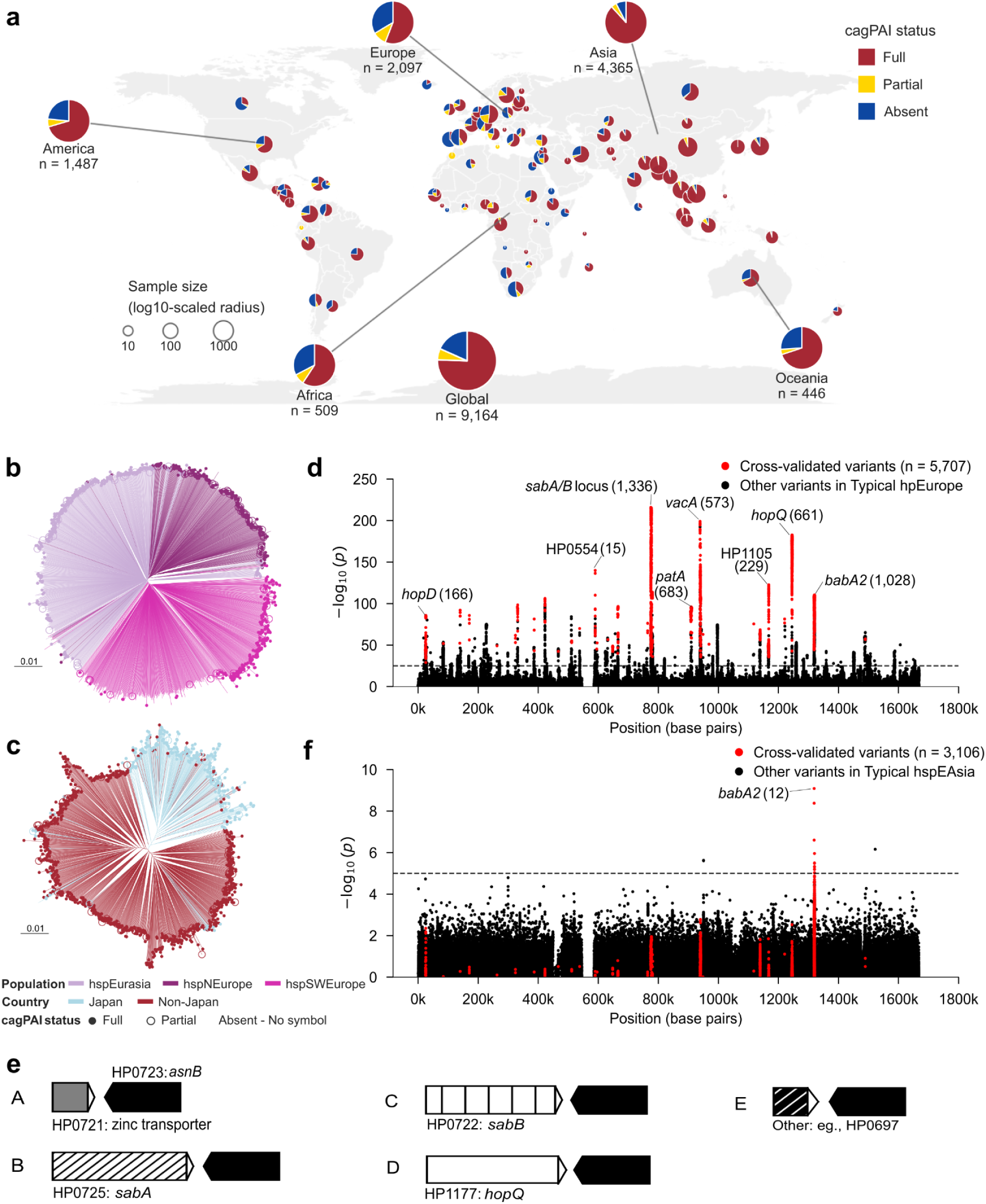
Population structure of *H. pylori* and *cag* status associated variants identified by GWAS. **a**, Global distribution of *H. pylori* cagPAI status. Pie charts indicate the proportion of strains with full (≥26 genes), partial (1–25 genes) or absent (0 genes) cagPAI in each country. Pie radii are scaled to the total number of isolates per country (log10-transformed). Geographic coordinates were obtained from publicly available datasets. **b**, Phylogenetic tree for 1,425 typical hpEurope genomes. Core genome SNP phylogenies were constructed using a neighbour-joining algorithm. Branches are coloured by population and subpopulation assignment. Tips indicate relevant countries of isolation. **c**, Manhattan plot from the GWAS analysis of *cagF*^+^ versus *cagF^−^* strains within the typical hpEurope dataset. A total of 15,976 variants exhibited –log_10_(*P*) > 25 (indicated by the black line). **d,** Phylogenetic tree for 1,179 typical hspEAsia genomes. **f,** Manhattan plot from the GWAS analysis of *cagF*^+^ versus *cagF*^−^ strains with the typical hspEAsia dataset. For Manhattan plots, variants in red overlapped with significant matches in the *cagF*^+/−^ GWAS performed in three European subpopulations. **e,** OMP configuration groups indicated by the SNP776,433 or SNP776,479 marker, downstream of HP0723 (black arrow). Arrows indicate gene orientation and length. Other genes are labeled above arrows, with hatching denoting OMP type (SabA, SabB, HopQ).

Within the typical hpEurope dataset, we performed a GWAS for *cagF* presence or absence as a marker for the cagPAI. Among the 407,886 tested SNPs, 15,976 exceeded a stringent *P*-value cutoff of –log_10_(*P*) > 25 **(Fig. 1d)**. To minimize potential false positive associations driven by population stratification within Europe, the GWAS was further refined by partitioning the collection into three distinct sub-populations: hspSWEurope, hspNEurope and hspEurasia. Sub-population-specific significance cutoffs were set at –log_10_(*P*) values of 20, 5, and 10, respectively **(Supplementary Fig. S2)**, reflecting differences in cohort size. The least well powered sub-population cohort is hspNEurope (**Supplementary Fig. S2b**), where the empirical cutoff of –log_10_(*P*) = 5 (unadjusted *P* =10^-5^) lies on the borderline of statistical significance under a standard Bonferroni correction (α = 0.05/380, 775 ≈ 1.3 × 10^-7^). Given that there is significant P value inflation (genomic inflation factors λ = 7.22, 2.15, and 3.59 for hspSWEurope, hspNEurope, and hspEurasia, respectively; **Supplementary Fig. S2d**), likely due to clonal and population-structure within the dataset, this GWAS is likely to produce some false-positives, motivating our conservative multi-population replication strategy. However, of the 7,661 SNPs identified in the hspNEurope GWAS at the empirical threshold of –log10(*P*) ≥ 5, 5,707 are replicated in both of the other two sub-population specific datasets, as well as the three-population combined GWAS (red variants, **Fig. 1d, Supplementary Table 2**), compared to an expectation of less than 20 SNPs if the three sub-population GWAS were independent of each other (**Methods**). Thus replication across three differentiated sub-populations ensures that our final list of 5,707 SNPs is unlikely to contain a substantial number of false positives due to clonal structure, population substructure or chance.

Unexpectedly, the strongest association signals amongst the triple hit SNPs were not in previously identified cag associated genes, such as *vacA* and *hopQ*, but instead mapped predominantly to the HP0721–HP0726 region of the 26695 reference strain, which harbored a total of 1,336 variants **(Supplementary Fig. S3, Supplementary Table 2)**. Specifically, the intergenic regions of HP0721|HP0722 contained the top hit in block HP0721–HP0726 region, B1H1, SNP 774,190 **(Fig. 1d, Supplementary Table 4),** where B denotes the block and H denotes the rank within that block, with both ordered by P value. Further inspection suggested that this variant is associated with the configuration of outer membrane proteins encoded at this position (**Fig. 1e**), and specifically the particular OMP encoded downstream of HP0723 (SabA, SabB, or HopQ). Specifically alignment-gap (-) allele is strongly associated with Configuration A in which the zinc transporter gene HP0721 is next to HP0723 (an asparaginase). Strains with the alternative A allele instead mostly carry Configurations B, C, and D, which have one of three outer membrane protein (OMP) genes, *sabA*, *sabB* and *hopQ*, encoded downstream of HP0723 **(Fig. 1e)**. Configuration E is rare, except in Hardy strains and corresponds to a gene other than HP0721 or an OMP at the same position and, like Configuration A strains, is mostly accompanied by the alignment gap (-) allele.

Loci previously associated with cag status identified by our GWAS included HP0887 (VacA; vacuolating cytotoxin), which included 573 variants **(Supplementary Fig. S3, Supplementary Table 3)** and other virulence-associated outer membrane proteins (e.g., HP1177, encoding HopQ, which facilitates CagA translocation; and HP1243, encoding blood group antigen binding adhesin BabA2). We also identified differentiated loci without prior links to *cag*. These included HP0554 (15 variants), annotated as Sialidase A; HP1105 (229 variants) encoding a glycosyltransferase involved in LPS Lewis antigen biosynthesis^10^, HP0855 (*patA*, peptidoglycan O-acetylation and lysozyme resistance^11^; 683 variants), and *hopD* (166 variants). Together, these findings support coordinated diversification between the cagPAI and toxin/outer membrane virulence functions in Europe, and also identify genes not previously associated with *cag* status.

We also performed a GWAS for presence or absence of *cagF* in the hspEAsia population, which identified few associations. The top hit was in HP1243 (BabA2) with –log_10_(*P*) = 9.1, which is weaker than the association of the same SNP in hpEurope of –log_10_(*P*) = 110.1 **(Fig. 1f**, **Supplementary Table 5)**. These results imply that there has been minimal coadaptation to *cag* status in East Asia.

### Polygenic Coadaptation Scores identify persistent *cag*^−^ lineages

Within the human genomics literature, polygenic risk scores (PRS) have become a popular way of summarizing the predicted effect of genomic variation on a trait^12^. For example, most tall people will have high polygenic scores for height, while people with high polygenic risk scores for a particular disease are more likely to be affected by it at some point in their lifetime^13^. We used the same methodology^14,15^, accounting for the effects of linkage by only including the top SNP based on P value within each gene, intergenic region or longer linkage block. The selected SNPs were thus represented by the 30 triple replicated loci (B1–B30), each represented by one top-ranked SNP (H1; **Supplementary Table 4**). For each strain, the cagPAI polygenic risk score (PRS) was calculated as the sum of the corresponding association-effect estimates (β) for loci at which its allele state differed from the pooled Typical European major state; gaps were retained as valid allele states in defining these differences (**Fig. 2a**).

**Fig. 2.**
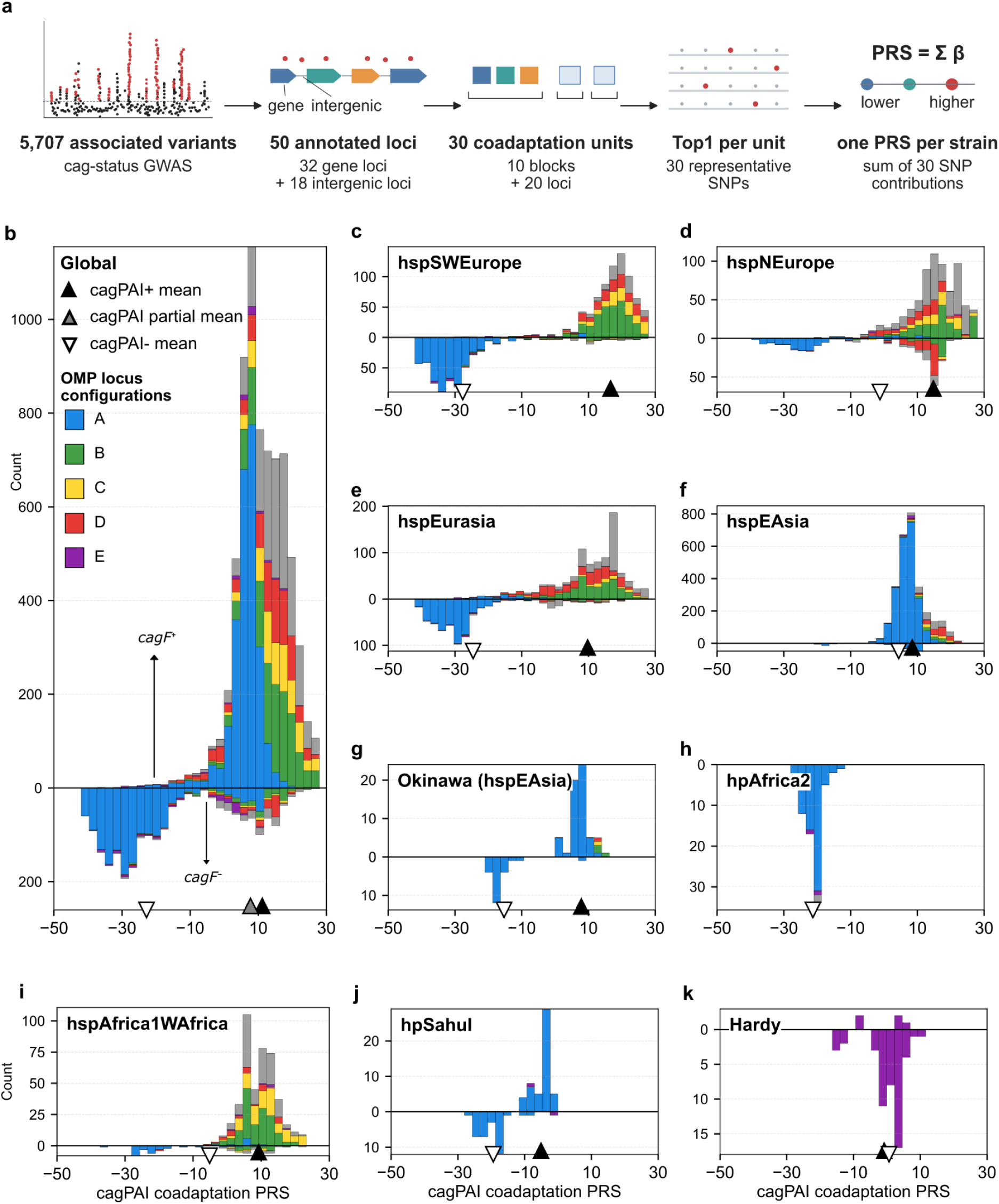
cagPAI coadaptation polygenic risk score, and its relationship with OMP configurations across global *Helicobacter pylori* populations. a, Workflow used to construct the cagPAI coadaptation PRS. A total of 5,707 cross-validated variants (red dots) associated with *cag* status were mapped to 50 annotated loci, comprising 32 genes and 18 intergenic regions. On the basis of physical in linkage (in the 26695 reference genome), these loci were consolidated into 30 coadaptation units, including 10 blocks and 20 genes. One top-ranked SNP per unit (B1H1–B30H1; 30 SNPs in total) was retained for PRS construction. The PRS for each strain was calculated as the sum of the allele-specific, effect-size-weighted contributions (β) of these 30 SNPs. **b-k**, Stacked bar plots illustrating the number of strains (n) exhibiting each HP0723 downstream gene configuration, stratified by *cag* status and the total PRS across global populations. **b** Global dataset; **c-e,** European subpopulations (hspSWEurope, hspNEurope, hspEurasia); **f & g,** Asian populations (all hspEAsia strains, hspEAsia strains from Okinawa only); **h & i**, African populations/subpopulations (hpAfrica2, hspAfrica1WAfrica); **j,** hpSahul and **k,** Hardy ecospecies. y-axis: strain count (*cagF*^+^ upper, *cagF*^−^ lower). x-axis: PRS. Upward-and downward-pointing triangles indicate the mean PRS for *cagF*+ and *cagF*− strains, respectively.

If we follow the logic used by human geneticists, our PRS should predict the chance (“risk”) that a strain carries the cagPAI, based on the variants in the rest of the genome. However, we know the actual *cag* status of the strain, so a more useful way to interpret the score is as reflecting how likely the strain is to continue to propagate successfully, while retaining its current *cag* status. Specifically, a strain with a high polygenic score has many variants that are statistically associated with *cag*^+^ strains in all three hpEurope sub-populations. Thus, assuming a polygenic model, in which each variant contributes independently to the ability of a *cag*^+^ strain to persist, a high score predicts that the strain is coadapted to the presence of the island and hence has a good chance of persisting in the long term, without losing the island. Conversely, *cag*^−^ strains are most likely to persist if they have low cagPAI PRS scores.

Across the global dataset, cagPAI PRS distribution is bimodal with most *cag^+^* strains having a high score, and most *cag*^−^ strains having a low score **(Fig. 2b)**. Although our polygenic score was defined using a GWAS for the hpEurope population, we found that *cag^+^* strains had higher average scores than *cag*^−^ strains in all populations, except in hpAfrica2 where there are no *cag*^+^ strains **(Fig. 2c-j)**. The concordance across multiple ancestry backgrounds validates our interpretation that the scores reflect coadaptation to *cag* status. However, the largest difference were observed in two of the three hpEurope subpopulations, namely hspSWEurope **(Fig. 2c)** and hspEurasia **(Fig. 2e)**, suggesting that we may systematically underestimate coadaptation in non-European populations. The bimodal distribution implies that *cag^−^* strains persist stably in the population and that many are well adapted to their status.

Notwithstanding this global trend, a minority (14%, 295 of 2,089) of *cag^−^* strains had a relatively high cagPAI PRS (> 10) **(Fig. 2b)**. Most of these (71%, 208/295) harbored partially intact islands. This suggests that most of these strains lost *cag* function recently and have not had time to undergo substantial adaptation to their new status. The cagPAI mediates CagA translocation into human cells, which relies on the structural and functional integrity of the encoded Type IV Secretion System (T4SS). Some accessory genes may be dispensable under laboratory conditions^16^, but in general the *cag* island only achieves its function of injecting CagA into human cells if the island is intact.

Outer membrane configuration in the HP0721–HP0726 region shows substantial variation between geographic populations. For example, within hspEAsia, 80% have Configuration A **(Fig. 2g, Supplementary Fig. 4-5, Extended Data Fig. 1)**, compared with only 30% in hpEurope. However, for hpEurope strains, Configuration A is strongly associated with being *cag^−^* **(Fig. 2d-f),** and these *cag^−^* strains are well adapted to the absence of the island, with a mean PRS of-30. A substantial minority (17%, 249/1,476) of hpEurope *cag^−^* strains carry Configuration B, C, or D but their mean PRS are higher (8, 10, 7), and a larger fraction have partial cagPAI islands (53%, 132/ 249), implying that on average, they have lost the island more recently and are substantially less adapted to absence of the island.

The distribution of scores differs markedly between subpopulations, but most follow one of two patterns. The first pattern is bimodality associated with stable *cag*^+^ and *cag*^−^ populations. hspSWEurope (**Fig. 2c**) is a good example, with few poorly coadapted strains circulating. Other bimodal populations include hspNEurope **(Fig. 2d),** hspEurasia **(Fig. 2e)**, hspNorthAsia **(Supplementary Fig. 6b)**, and hpNEAfrica **(Supplementary Fig. 6d)**. However, a substantial fraction of the *cag^−^* strains in the hspNEurope population are poorly coadapted to their status, with many of these strains having Configuration B or D.

The hspEAsia population is an exemplar of the second pattern, with little evidence of coadaptation to cagPAI loss. Only 1.8% (55/3,081) of all hspEAsia strains completely lack the island, while 5.1% (157/3,081) have a partial island **(Supplementary Table 1)**. This suggests that in most cases loss of function has occurred recently and that most *cag^−^* strains are transient in this population. Consistent with this, over 99% of all hspEAsia strains with partially intact islands had a cagPAI coadaption PRS greater than **-**10, with a median score almost identical to that of strains carrying a complete island (8.45 versus 8.50). Other populations with similar patterns to hspEAsia include hpAsia2 and hspSAsia **(Supplementary Fig. 6a,c)**, as well as hspCNEAfrica and hspAfrica1SAfrica **(Supplementary Fig. 6e,f)**.

### Dynamics of coadaptation to *cag* status

To study the dynamic process of coadaptation we use pairwise comparisons between strains to measure how differences at PRS loci accumulate as a function of genome-wide divergence (**Fig. 3**). We calculated the average divergence levels for pairs of *cag^−^*, *cag*^+^, or *cag^+/−^* strains based on the top representative variant (H1) for each of the 30 differentiated blocks (B1H1–B30H1, **Supplementary Table 4**), as well as their genome-wide SNP distance. There are two versions of these plots for each population, the first containing all H1 loci (all variant types, including gaps) (**Fig. 3a**), and the second containing only H1 SNP loci (**Fig. 3b**). For the second set of plots, strains were also compared with the expectation based on SNPs with matched gene frequency elsewhere in the genome (control, black line).

**Fig. 3.**
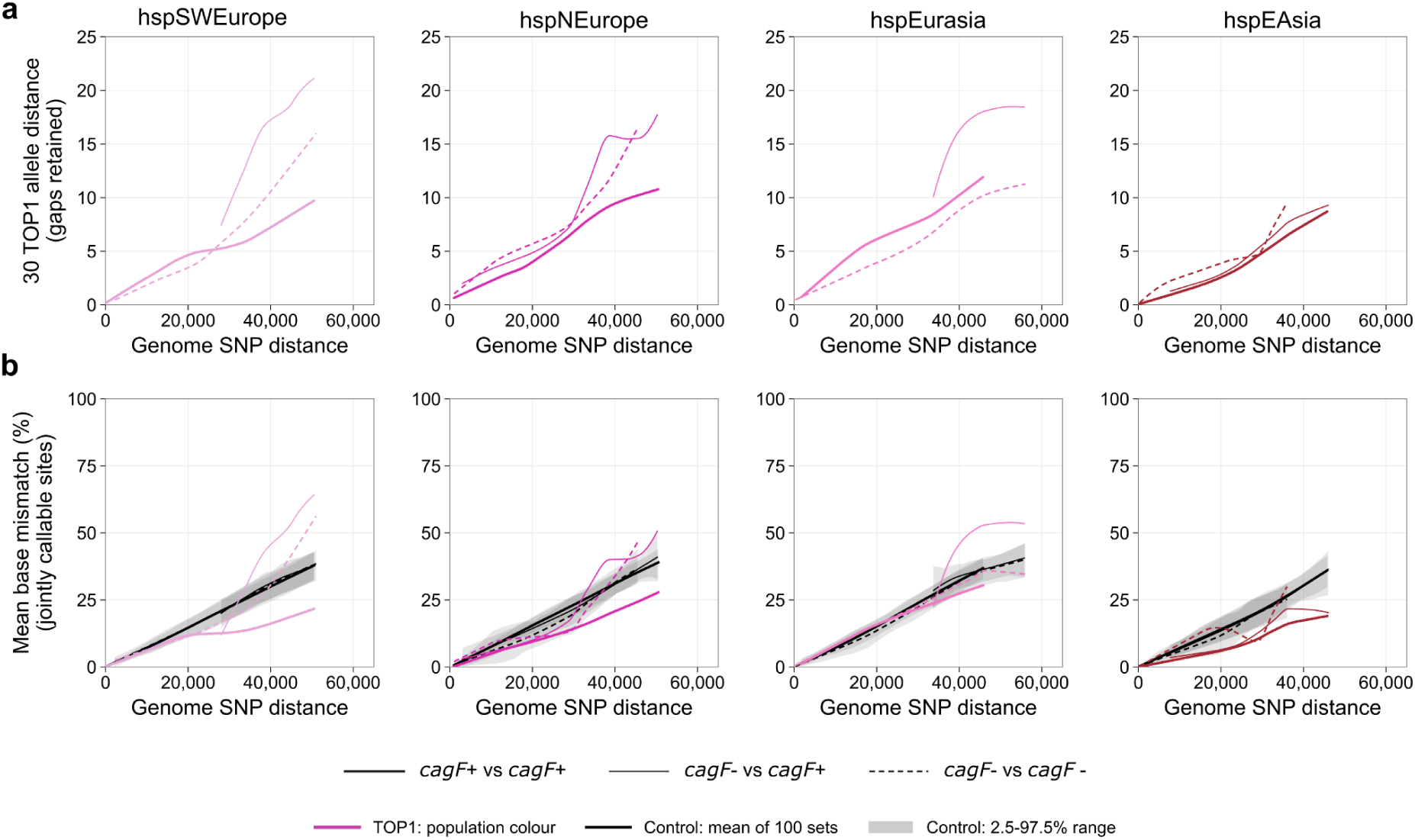
Dynamic, population-specific relationship between genome-wide SNP distance and cagPAI-associated allele distance. Each panel shows one population or defined subgroup. The x axis denotes pairwise whole-genome SNP distance between pairs of strains, grouped into 5,000-SNP windows. In **a**, the y axis shows the mean total allele distance across 30 block-representative H1 loci, with A/C/G/T/gap treated as distinct states; thus, gap–base differences are retained. In **b**, the y axis shows the mean per-pair nucleotide mismatch proportion across the same loci, calculated only at positions at which both strains carry callable A/C/G/T bases; comparisons that include a gap are excluded. Line width and style denote *cagF* pairing class: thick solid line, *cagF*^+^/*cagF*^+^; thin line, mixed *cagF*^+^/*cagF^−^*; and dashed line, *cagF^−^*/*cagF^−^*. Coloured curves show the 30 H1 loci, whereas black curves and grey bands show the mean and empirical 2.5th–97.5th percentile range across 100 region-excluded, globally frequency-matched control sets. Curves summarize 5,000-SNP windows containing at least 10 pairs per class.

In hspSWEurope and hspEurasia, pairs of strains that differ in *cag* status show the largest genome-wide SNP distances of any comparison, and their mean H1 distance exceeds that of every other pairing at all genetic distances (**Fig. 3a**). This is consistent with both *cag*^+^ and *cag^−^* populations being ancient within these groups: enough time has elapsed for differences between them to accumulate and to be maintained by selection. In hspNEurope and hspSWEurope, by contrast, H1 distances among *cag*^+^ pairs fall well below the control-SNP expectation (**Fig. 3b**), as expected for a stably coadapted *cag*^+^ population. In both populations the curves are consistent with loss of the island by *cag^−^* strains, but the timing differs. In hspNEurope, some pairs combine low SNP distance with low H1 distance – the signature of a recent island loss, with little time for coadaptation to erode. In hspSWEurope, H1 distances between *cag^−^* strains rise sharply once pairwise SNP distance exceeds 30,000, suggesting instead that the cagPAI was lost several times independently in the more distant past, and that these independent losses have not converged on a common coadapted state.

A more surprising pattern appears in hspEurasia: H1 distances between *cag*^+^ strains exceed those between *cag^−^* strains at the same genome-wide divergence (**Fig. 3a**). One explanation is that the *cag*^+^ population is the derived state, arising several times independently from *cag^−^* ancestors. Alternatively, different *cag*^+^ strains may have adopted different coadapted strategies, so that elevated H1 diversity within *cag*^+^ reflects divergent selection rather than independent origins. The two hypotheses are not mutually exclusive. It is also noteworthy that *cag^−^* and *cag*^+^ strains differ consistently in their ancestry profiles in hspSWEurope but not in hspEurasia (**Extended Data Fig. 1**). This is consistent with separate geographic origins of *cag^−^* and *cag*^+^ strains, before they came together and exchanged genes in most of the genome to form the modern hspSWEurope population. Within hspEurasia it is more likely that differentiation of *cag* status and subsequent coadaptation happened in one geographic region.

Within hspEAsia, H1 genes evolve slowly in *cag*^+^ strains (**Fig. 3**), as expected based on stabilizing selection and pairs with differing status have a similar curve to that for *cag*^+^ strains, consistent with repeated loss of the island, and minimal coadaptation. In most other populations, the *cag*^+^ pairs are also lower than the control, consistent with stabilizing selection. There are also several populations with high distances between *cag^−^* pairs at large genetic distances, including hspNorthAsia, hspWAfrica1WAfrica and hspIndigenousAmerica **(Supplementary Fig. 7)**.

Together, these dynamic trajectories reveal that maintaining an intact cagPAI imposes pervasive stabilizing selection across global lineages, constraining divergence at coadapted loci. Conversely, recurrent island loss precipitates independent divergence among persistent *cag^−^* lineages across evolutionary time, leading to different adaptive outcomes that remain stable over long time intervals.

### Example of recent changes of cagPAI status

While the great majority of *cag*^−^ hspEAsia strains have positive coadaptation scores, indicating minimal coadaptation to *cag* loss, there are 25 strains with coadaptation scores less than-10 (**Fig. 2g,h**, **Fig. 4a, Supplementary Table 1**). Strikingly, these strains have distinctive ancestry profiles from other *cag*^−^ hspEAsia isolates, with minority ancestry from hspIndigenousAmerica and hpNorthAsia **(Fig. 4c)**. These strains are all from Japan, with 22 of them coming from Okinawa (**Fig. 2h**, **Fig. 4c**). We ran clonalframeML^17^ and bactdating^18^ to estimate the relationship between the strains. They come from a “deep clone”^2^, with an estimated most recent common ancestor dating to circa 1800 (95% highest posterior density interval: 1755–1862; **Methods**), but with two subclones with more recent estimated common ancestors **(Fig. 4a)**.

**Fig. 4.**
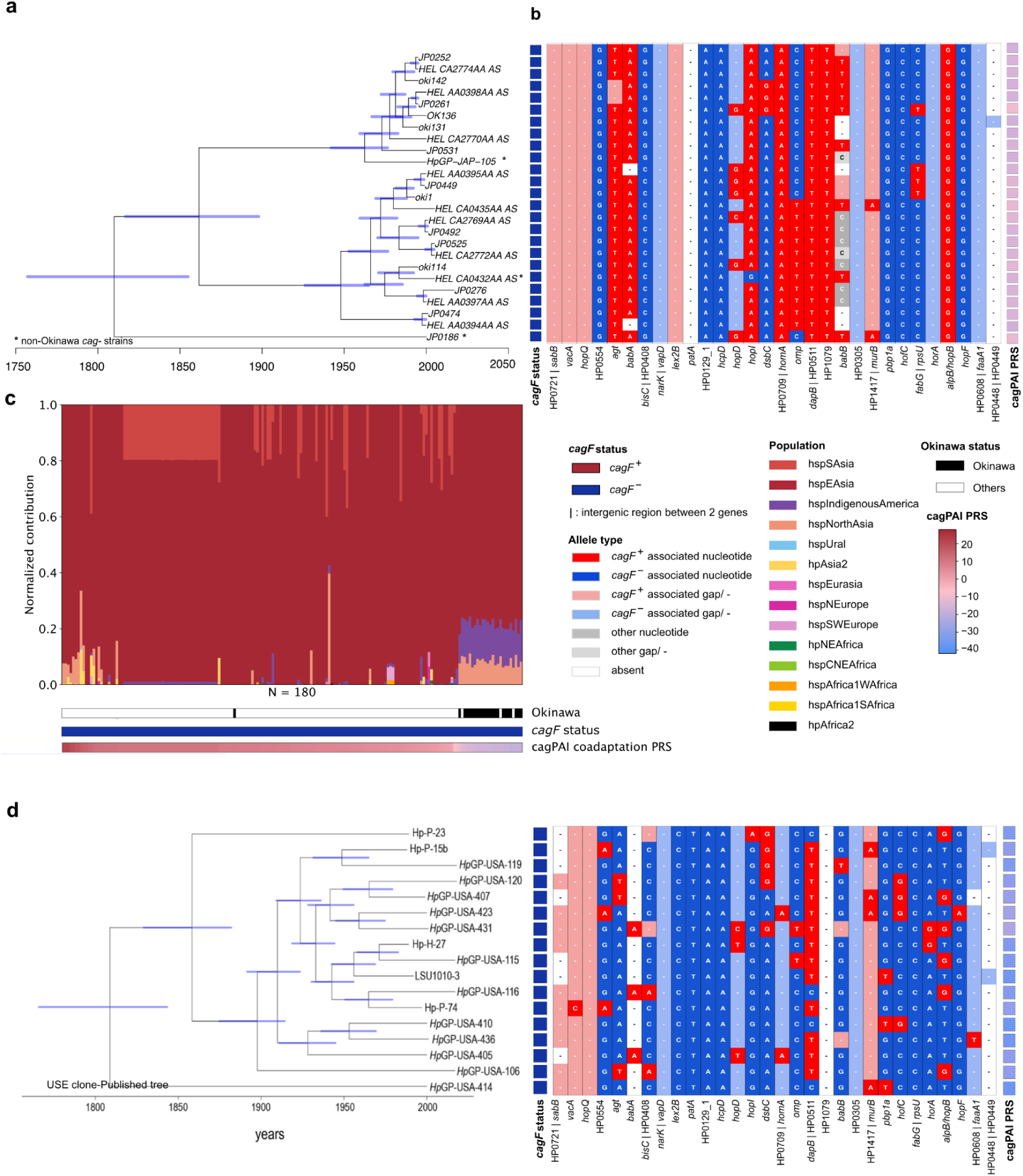
Genomic profiles, phylogenetic relationships, and coadaptation scores of Japanese and North American *cag*^−^ clonal lineages. a, Dated ClonalFrameML tree of 25 adapted *cag*^−^ hspEAsia isolates. Asterisks (*) denote non-Okinawa strains within the predominantly Okinawa lineage. **b**, cagPAI coadaptation associated Haplotype structure across 25 adapted *cag*^−^ hspEAsia isolates. **c**, Ancestral component analysis of 180 *cag*^−^ hspEAsia isolates. **d**, Dated ClonalFrameML tree analysis and cagPAI coadapted Haplotype structure for 17 *cag*^−^ USA clonal isolates. The ClonalFrameML tree has been published previously^2^.

It is not clear where strains in this deep clone acquired their minority ancestry, or the variants adapting them to *cag* loss, but there is relatively little variation between them at *cag* coadaptation loci, likely reflecting the lack of *cag*^−^ coadapted donors that are available within the Japanese *H. pylori* population where they are currently circulating. This example, however, shows that hspEAsia strains can coadapt to *cag* loss and spread within East Asia, provided that they have access to an appropriate gene pool. A *cag*^−^ clone of a similar age has previously been identified in North America^2^. This clone has lower polygenic scores than the Japanese clone (PRS ranges:-38.5 to-26.9 and-19.2 to-10.7, respectively, **Fig. 4d**), consistent with greater coadaptation to cagPAI absence. The better coadaptation is unsurprising, since within the North American gene pool there are many other *cag*^−^ lineages that the strains can acquire coadapted variants from.

We also attempted to identify examples of recent gain of the cagPAI. There are 40 *cag^+^* strains with a PRS less than-15.85, all from the hpEurope population **(Supplementary Fig. 8a)**. It is surprising for *cag^+^*strains to have such low scores and one simple explanation is that they acquired the island recently, so that they have only recently begun to adapt to their new status. Interestingly, the four *cag^+^* strains with the lowest scores were all found in hspEurasia (three in Iran and one in Israel), possibly suggesting recent selective pressure for acquisition of the island in those locations. However, the low coadaptation scores of these strains provides only circumstantial evidence for recent gain. Only one of the 40 strains was clonally related to a *cag*^−^ strain, and within that pair of strains the *cag*^+^ strain had a slightly higher score than the *cag*^−^ strain (-18.4 vs-19.8). The two strains differ at five of the SNPs used in the polygenic score, with three going in the expected direction, which does not represent convincing evidence of coadaptation **(Supplementary Fig. 8a)**.

An alternative approach to identifying instances of acquisition of the cagPAI is to find pairs of strains whose *cag* islands are closely related, although the rest of their genomes are not. Analysis of the global dataset identified two closely related pairs, which involved three isolates from Russia **(Supplementary Fig. 8b)**. This example might provide evidence that the cagPAI can be transferred in its entirety; however, all three strains have positive polygenic scores **(Supplementary Fig. 8b)**, so it is also not obviously an example of *cag* gain in a strain previously coadapted to the absence of the island and we have found no examples where we have strong evidence for recent acquisition of the island in previously *cag*^−^ strains. Therefore, we have no strong smoking guns for recent cagPAI gain within the current dataset.

### Gene-by-gene patterns of coadaptation

Although the overall distribution of PRS is bimodal, the continuous nature of the score and the presence of strains with intermediate scores allowed us to investigate whether there are stages of adaptation to cagPAI status. There may, for example, be variants which only become beneficial on either a *cag*^+^ or a *cag*^−^ background once a degree of coadaptation has already occurred. To investigate this possibility, we divided Typical Europe strains into four categories based on *cag* status and cagPAI PRS (adapted or intermediate) (**Fig. 5a**). For example, *cag*^−^ strains (**Fig. 5a**, left) were separated into those with low cagPAI scores (PRS <-30, highly adapted to being *cag*^−^) and intermediate scores (-30 > PRS >-17, less adapted to the cagPAI). We then performed three separate GWAS: between *cag^−^* strains with low vs. intermediate PRS scores (set 1 vs. 2), between strains with intermediate scores, *cag^−^* vs. *cag*^+^ (set 2 vs. 3), and between *cag*^+^ strains with intermediate vs. high PRS (**Fig. 5a, b**).

**Fig. 5.**
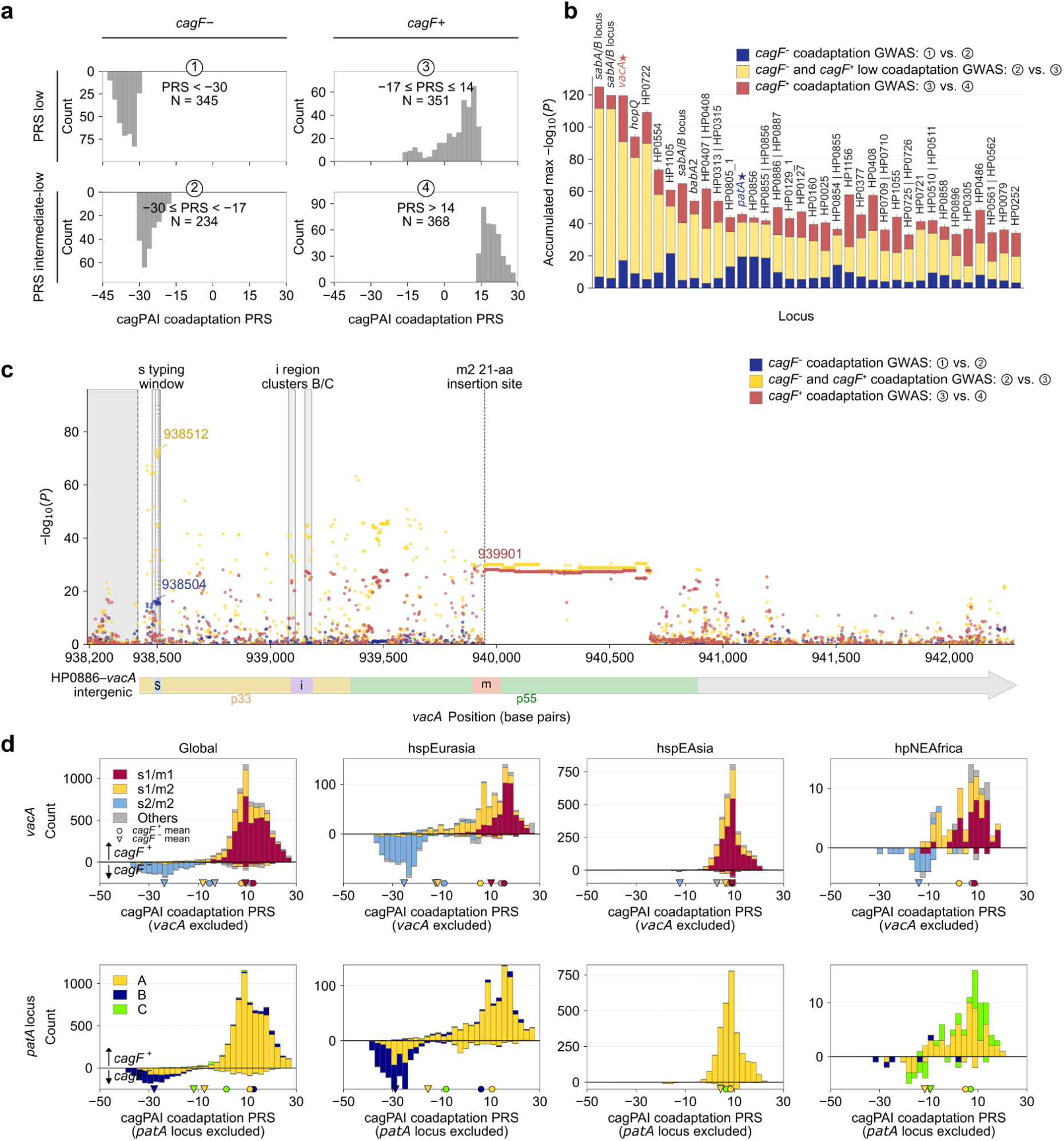
Variants associated with intermediate cag adaptation scores map to the *vacA* and *patA* regions. a, Distribution of cagPAI PRS among four groups (for details see **Methods**) of typical hpEurope strains used for GWAS, defined by *cagF* status and cagPAI PRS value. **b,** Gene-level signals from three GWAS comparing the four groups of typical hpEurope strains classified by *cag* status and cagPAI PRS. The 50 regions with the highest accumulated signals were selected and ordered by their maximum –log_10_(*P*) in the original *cagF*^+^ versus *cagF*^−^ GWAS. **c,** Association signals for intermediate strain set comparison GWAS at the *vacA* (HP0886-HP0887) region in strain 26695. Signals are shown for strain sets 1 versus 2 (blue, *cagF*^−^ PRS-low vs. PRS-high), 2 versus 3 (yellow, *cagF*^−^ PRS-high vs. *cagF*^+^ PRS-low) and 3 versus 4 (red, *cagF*^+^ PRS-low vs. PRS-high). Shading and vertical lines mark the intergenic region and the s-, i- and m-typing features of *vacA*. The three strongest variants in each comparison are labelled by genomic and VacA amino-acid position. **d,** Distribution of *vacA* and *patA* types according to cagPAI coadaptation PRS. Mirrored histograms show *cag*^+^ strains above and *cag*^−^ strains below the axis for the global dataset and geographical groups. Circles and triangles denote the mean PRS of *cag*^+^ and *cag*^−^ strains, respectively, and are shown for groups with at least ten strains. Colours indicate *vacA* and *patA* types.

The most interesting differences between these three GWAS (**Supplementary Fig. 9**) are found in the *vacA* gene. The SNPs most strongly associated with cagPAI status in our original GWAS presented above are in the *vacA* signalling (s) region (**Fig. 5c**), and this is recapitulated in the GWAS based only on *cag*^+^ and *cag*^−^ strains with intermediate PRS scores (**Fig. 5b,c**, *yellow***; Supplementary Fig. 10**) but less so in the other two GWAS (blue, red), suggesting that variation in the s region is key to adapting to presence or absence of the cagPAI.

Highly coadapted *cag^+^* strains differed from less adapted *cag^+^* strains more strongly in the m region **(Fig. 5b,c**, *red*), suggesting that subsequent fine-tuning across the intermediate-to-high coadaptation gradient occurs in this part of the *vacA* gene. Previous work^19^ has defined *vacA* haplotypes to be based on short insertions within the s region and m region. These haplotype definitions give concordant results. Specifically, the s region haplotype is strongly associated with *cag* status globally, while the mean polygenic score of *cag*^+^ s1m2 strains is lower than that of *cag*^+^ s1m1 strains in all populations with more than 10 isolates in both categories (**Fig. 5d**), suggesting that this region is associated with the extent of coadaptation to *cag*^+^ status.

A handful of genes showed the most significant P values in the GWAS between *cag*^−^ strains with low and intermediate PRS scores **(Fig. 5b**, *blue bars*). For example, the HP0854–HP0855 *patA* region has two highly distinct haplotypes (**Supplementary Fig. 3**), which suggests a long-standing polymorphism (**Supplementary Fig. 11a**). Based on this we grouped the global strains into three types by haplotype and score (A, B, C; **Fig. 5d**, *bottom;* **Supplementary Fig. 11a**). In two of the three hpEurope subpopulations, the rarer type B is associated with strongly adapted *cag*^−^ strains. However, this association is not found in other populations (**Fig. 5d**; **Supplementary Fig. 11b**) and in hspSAsia (**Supplementary Fig. 11c**), the same haplotype is surprisingly instead associated with the most strongly adapted *cag*^+^ strains. Thus, the relationship to this ancient polymorphism to cagPAI coadaptation appears to be complex.

### Origin of cagPAI status-associated variants

To investigate the coadaptation that took place when the cagPAI first arrived in the *H. pylori* genome, we selected a small panel of coadapted *cag^+^* and *cag*^−^ strains that encompass the ancient diversity of the species **(Supplementary Table 1)**, including strains from hpSahul (**Fig. 2k**) and Hardy strains (**Fig. 2j**), whose *cag^+^*strains have substantially lower coadaptation scores than *cag^+^*strains from other populations. One Hardy lineage, *H. acinonychis*, is related to hpAfrica2 strains, indicating an African origin, but made an ancient host jump to big cats^3^. A strain isolated from a primate in the Davis Primate Centre was also Hardy and was most closely related to the hpSahul population, but distinct from all modern isolates, suggesting another ancient host jump^3^.

We clustered strains based on *cag* status-associated variants **(Supplementary Table 4)** and compared the resulting tree **(Fig. 6a)** with that obtained when clustering with ecospecies differentiated genes **(Fig. 6b)** and ecospecies non-differentiated genes **(Fig. 6c, Supplementary Table 6)**. As previously described^3^, the ecospecies differentiated and non-differentiated genes gave strikingly different trees, with the former reflecting the differentiation between Hardy and Ubiquitous strains, which predates the origin of modern humans, and the latter reflecting the geographic populations shaped by the subsequent migration history of the humans who carried them.

**Fig. 6.**
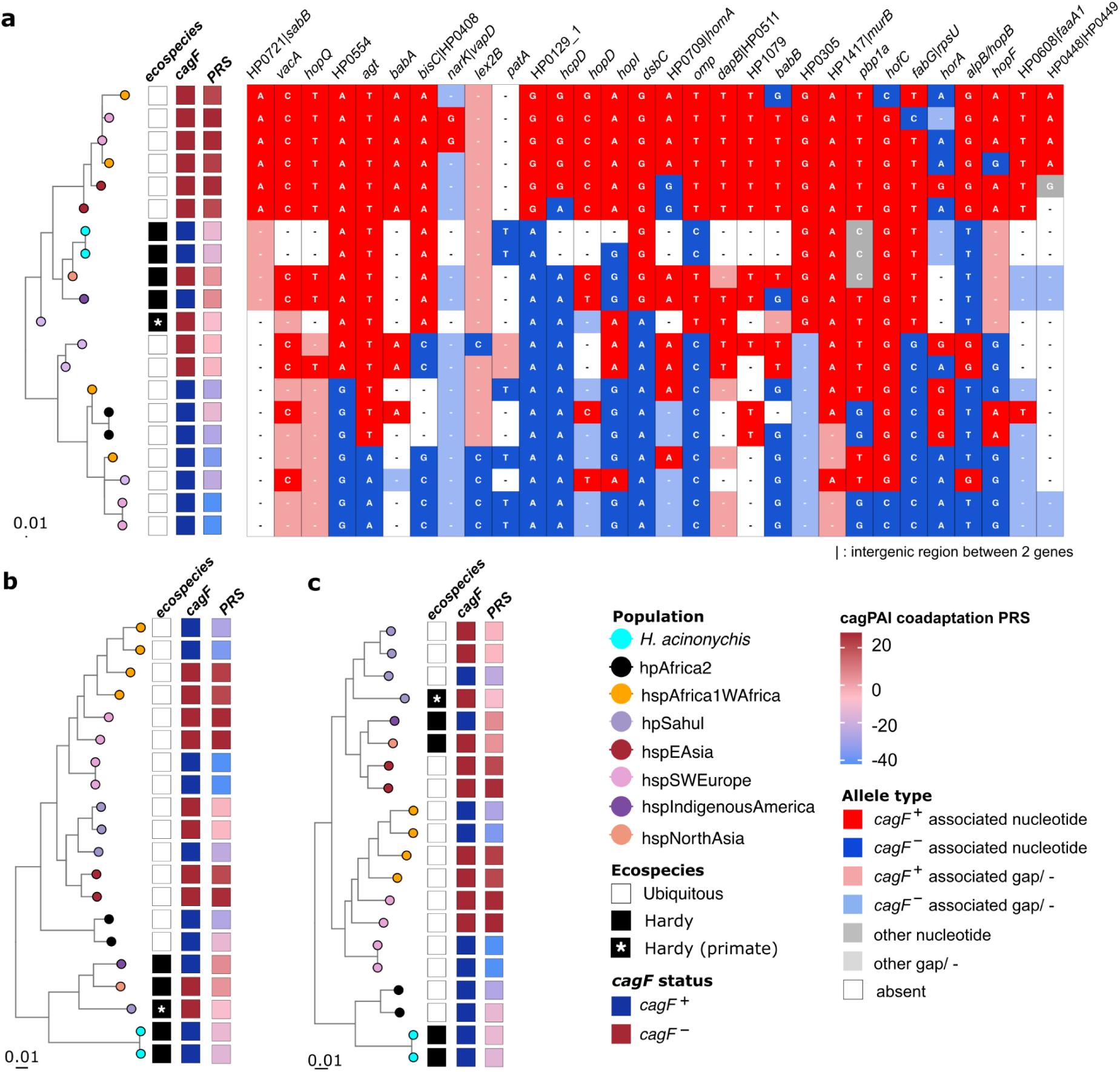
Phylogenetic clustering of *H. pylori* coadapted *cag^+^* and *cag*^−^ strains and haplotype structure. a, Tree of representative *H. pylori* strains based on cagPAI-associated variants aligned with a haplotype map for *cag*-associated SNPs. Left: the evolutionary relationship based on cagPAI-associated variants. Tip colours: population. The coupled heatmap (right) is based on 30 selected genomic loci (**Supplementary Table 4**). **b, c,** Phylogenetic reconstructions based on Hardy ecospecies highly differentiated or non-differentiated genes. Scale bars represent 0.01 substitutions per site.

The tree based on *cag* status associated variants **(Fig. 6a)** was distinct from the other two trees, and had clusters associated with *cag^+^* strains from hpEurope, hpAfrica1, and hspEAsia and a second cluster with *cag*^−^ coadapted strains from hpEurope, hpAfrica1 and hpSahul. Two hpSahul *cag^+^* strains clustered in an intermediate position. Crucially, the two outgroup hpAfrica2 strains clustered within the *cag*^−^ clade, confirming that this profile represents the ancestral species state.

Most interestingly, four of the five Hardy strains clustered within the *cag^+^* strains in this tree **(Fig. 6a)**, indicating shared derived alleles acquired through directional introgression rather than ancestral retention. Across these 30 loci, this polarity defines three distinct classes: ancestral alleles retained by hpAfrica2 and European *cag*^−^ lineages (yielding the basal score of ∼-20), derived coadapted alleles shared between Hardy and modern *cag^+^* strains (e.g., in HP0305), and unadapted intermediate configurations in hpSahul. The Hardy primate isolate clustered more deeply but was also clearly more closely related to cagPAI-positive strains than cagPAI-negative strains at these genes.

We propose that the clustering of Hardy and *cag^+^* strains together based on *cag*-associated variants occurs because many of these variants were initially associated with Hardy but were imported by Ubiquitous *cag^+^* strains as part of the initial adaptation to presence of the island. For example, gene HP0305 shows a particularly simple pattern, with three variants, including the most strongly associated variant shown in **Fig. 6a**, differentiating the top half of the tree from the bottom. The other two variants in HP0305 show the same pattern (**Extended Data Fig. 2**). This pattern can be explained if the variants were initially specific to Hardy strains but were imported by Ubiquitous *cag^+^* strains. The only *cag^+^* strains which lack these variants are the two *cag^+^*strains from hpSahul.

The hpSahul population is associated with Indigenous Australian and Papua New Guinea human populations, reflecting their split from other human groups 50,000-70,000 years ago, followed by long-term regional isolation and the *H. pylori*, hpSahul population has had limited subsequent contact with other *H. pylori* populations until modern times^20^. Concordant with its history of separate evolution, the hpSahul population is highly bottlenecked and shows low variation at *cag*-associated variants **(Fig. 2j**, **Extended Data Fig. 1**), but with small but consistent differences in polygenic score between *cag^+^* and *cag*^−^ strains, indicating stable coadaptation. We propose that the cagPAI was imported by *H. pylori* prior to the original human population split but at that point had only undergone limited coadaptation, explaining why many of the *cag*^+^ coadapted variants identified in hpEurope are not shared with hpSahul, despite being found in all other populations with *cag*^+^ strains.

While many genes (**Fig. 6a, Supplementary Table 4**) show allele distributions similar to HP0305 at *cag* status associated variants (*e.g*., HP0377, encoding a thiol–disulfide interchange protein), other genes showed distinct patterns. For example, at the *patA* gene block (683 variants in HP0855, **Supplementary Table 3, Extended Data Fig. 2**), *H. acinonychis* strains share the type associated with very low coadaptation scores in hpEurope, consistent with a longstanding polymorphism in this genome region.

## Discussion

Many pathogens have arisen through import of pathogenicity islands; thus, it is of considerable interest to find out how their genomes adapt to their presence^21,22^. Our analysis shows that after the cagPAI arrived in the *H. pylori* genome, substantial coadaptation occurred via homologous recombination. However, the cagPAI can be disadvantageous in some circumstances and our analysis shows that strains that have lost the island have also re-acquired variants from the original *cag*^−^ gene pool. Thus adaptation to this major virulence island extends beyond the island itself and involves coordinated changes across the wider genomic background.

We used variation associated with the cagPAI within hpEurope strains to define polygenic coadaptation scores, whose purpose is to estimate how well adapted each strain is to presence or absence of the island. High scores indicate that the strain is adapted to carrying the island, while low scores indicate adaptation to its absence. The validity of our polygenic score firstly depends on the validity of the GWAS used to identify causal loci. Accordingly, we only included GWAS hits within our polygenic scores that are “triple replicated”, with significant associations with *cag* status found in three European subpopulations as well as the entire HpEurope dataset.

Our unusually strict replication protocol does not imply that every associated SNP is causal, because many are found within strong linkage blocks. Further, even if the loci used to build the polygenic score are correctly identified as coadapted, our approach might nevertheless underestimate the extent of coadaptation outside the hpEurope population either because different variants are involved, or because interactions with the cagPAI depend on genetic background or environmental factors^23,24^. Notwithstanding these potential pitfalls, *cag*^+^ strains have higher mean scores than *cag*^−^ strains in all *H. pylori* populations harboring the island. Indeed, in most populations, the mean coadaptation scores of *cag*^+^ strains are between +8 to +18, a narrow range that implies that many of the same variants are used worldwide.

To interpret the scores, we have to consider both ancient and recent *cag* evolution. Our presumption is that the hpAfrica2 population is the best available proxy for *H. pylori* prior to the arrival of the island. Accordingly, these strains have a relatively narrow range of scores around-20. Most of the variants that they share with hpEurope *cag*^−^ strains are thus likely to be ancestral alleles, present before the cagPAI was acquired by *H. pylori*, rather than newly evolved adaptations to *cag*^−^ status. Among populations containing *cag*^+^ strains, the lowest scores are found in the hpSahul population, where all scores were negative. Since people from Sahul have had a long period of separate evolution^9^, the most likely explanation is that the *cag* island had already been acquired, but had only undergone limited coadaptation, when this population became isolated from the rest of the *H. p*y*lori* gene pool.

*cag*^−^ strains with low scores might be like hpAfrica2 strains, that is to say they never acquired the island, or they may have had the island and coadapted to it, before losing it and undergoing secondary coadaptation to their new status. We can begin to distinguish between these hypotheses by considering the dynamics of *cag* evolution. Within hpEurope, the scores are bimodal, which is consistent with long term persistence of *cag*^−^ lineages. However, we can interrogate the dynamics more directly by looking at how divergence at coadaptation loci varies as a function of genetic distance and *cag* status.

In the hspEAsia population, most of the *cag*^−^ strains (specifically lacking the *cagF* gene) have partially intact islands and many are closely related to *cag*^+^ strains, which is consistent with recent loss of *cag* function. Furthermore, they show limited coadaptation to cagPAI absence, while *cag*^+^ strains retain similar allelic configurations at coadaptation loci over long time periods (**Fig. 3**, rightmost column). These results are consistent with *cag*^−^ strains arising repeatedly, but being transient.

It is unknown why *cag*^−^ strains are rare and generally fail to persist in East Asia. One intriguing possibility is the historical absence of a gene pool coadapted to absence of the island. The only strains with scores of-10 or less belong to a *cag*^−^ “deep clone”, consisting of 25 hspEAsia strains, 22 from Okinawa and three from the Japanese mainland, which have an estimated common ancestor around 200 years ago. The existence of this lineage shows that hspEAsia strains can evolve into persistent *cag*^−^ lineages and spread within East Asia under modern conditions, but only if they have access to an appropriate gene pool of *cag*^−^ associated variants.

The three hpEurope subpopulations show distinct patterns, which give intriguing evidence about different adaptive dynamics. For this purpose we contrasted genetic distance across the genome as a whole with coadaptation distances, which are based only on the 30 coadapted regions. In hspEurasia and hspSWEurope, the genetic distances between *cag*^−^ and *cag*^+^ strains are high, implying most lineages have persisted with their current *cag* status for hundreds or thousands of years, while in hspNEurope lower distances show that new *cag*^−^ lineages have arisen recently.

If *cag*^−^ strains have a single origin within a population, then we might expect coadaptation distances between pairs of *cag*^−^ strains to be low, while if they have independent origins, then the distinct lineages will have adapted via different paths and may have high coadaptation distances. The hpEurope subpopulations show different patterns. In hspEurasia, the coadaptation distances for *cag*^−^ strains are low, consistent with a single origin or ancestral retention of *cag*^−^ status. However, for hspNEurope and hspSWEurope, the distances bend upwards for more distantly related pairs of *cag*^−^ strains, which provides evidence for multiple origins of *cag*^−^ lineages.

Aside from different histories, variation in coadaptation scores between strains with the same *cag* status can arise because they have evolved distinct evolutionary strategies. For example, the *vacA* cytotoxin is highly polymorphic^25^, with many of those polymorphisms associated with cagPAI status. Polymorphisms in the s-region determine the efficiency with which the toxin is secreted and inserts itself into host cell membranes^26^, while polymorphisms in the m-region determine cell specificity^26–28^. The strongest associations with cagPAI presence are found in the s region. However, if we restrict analysis to *cag*^+^ strains, then the genotype associated with high polygenic scores is the m1 allele, while the m2 allele is found preferentially on strains with intermediate scores. This association of specific alleles with intermediate scores implies that the coadaptation is not simply a matter of becoming more compatible with the presence, or absence, of the island but reflects different strategies for interacting with the host.

Based on previous functional genomics work, we propose that *cag*^+^ strains, together with coadapted alleles of *vacA*, *babA*, *hopQ*, and *sabA*, drive a more intimate and aggressive interaction with host cells, exploiting inflammation through mechanisms such as induction of the SabA ligand sialyl-Lewis X and increased iron availability – at the potential cost of increased immune exposure^1^. In contrast, *cag*^−^ strains may persist within the mucus layer, adopting a more immune-evasive lifestyle, or exploit the modifications generated by co-infecting *cag*^+^ strains. HopQ binds CEACAM receptors and facilitates CagA translocation by the T4SS, while BabA has also been shown to aid CagA secretion^29^, providing a plausible molecular mechanism for these associations. The reason for CagA and VacA interaction is less clear and may be more indirect.

Many of the *cag* coadaptation associated genes that we identified are unstudied, including the fourth most strongly associated gene, HP0554. This gene is annotated as encoding a sialidase A, but *H. pylori* does not appear to express sialidase activity^30^. Moreover, a structural homology-based search using Phyre^31^ and InterProScan^32^ did not identify a sialidase domain in HP0554. Instead, we detected putative transmembrane and HTH (helix-turn-helix) nucleic acid binding (Lambda repressor-like) domains. Differentiated SNPs have been identified in HP0554 in a handful of population genomics studies^24,33^, but otherwise, to our knowledge, the gene has not been flagged as related to the cagPAI or virulence. While the molecular details of how HP0554 contributes to *Helicobacter* ecology in the stomach remains unclear, this highlights the potential of coadaptation screens to identify putative novel host interaction factors in an unbiased manner.

We identified an intriguing association of gene configurations at the HP0721–HP0726 locus with *cag* status. Most strains adapted to *cag* loss in hpEurope carried configurations without an additional OMP gene (in addition to *sabA*) downstream of *asnB* (Configurations A and E). In strains such as 26695, this position is occupied by *sabB*, while other strains encode a second copy of *sabA* or *hopQ* here. An analysis of North American clinical strains likewise found that *cag*^−^ strains also lacked an OMP downstream of HP0723^34^. We found no evidence that Configuration A and E strains encode *sabB* elsewhere in the genome, suggesting that *sabB* is either dispensable, or unproductive, in a *cag*^−^ genetic background.

The specific function and ligand of SabB is currently unclear. However, the association of the *sabB*^−^ configuration with adapted *cag*^−^ strains in Europe could also reflect how *H. pylori* regulates its OMP repertoire. The paralogous OMPs *sabA*, *sabB*, and *hopQ* undergo gene conversion, driven by recombination at homologous 5**′**/3**′** ends, which can alter adherence^34^. Because Configuration A/E strains lack an OMP gene downstream of HP0723, they may not have access to this mechanism. A possible reason for retaining this mechanism would be to allow *sabA* duplication in response to the inflammation-induced sialyl-Lewis X receptor promoted by the *cag*PAI and toxigenic VacA to take advantage of this increase in ligand ability, or to upregulate HopQ expressed to facilitate CagA translocation^35^. Losing this mechanism would avoid the increase in association with the host that changes in adhesins might confer, allowing *cag*^−^ strains to prevent unnecessary interaction with the host immune response. However, as Configuration A is predominant in HspEAsia strains, which are mostly *cag*^+^^36^, the benefit provided by this configuration requires further investigation.

Notably, we found that several cagPAI-adapted alleles appear to be shared with the Hardy ecospecies, including *vacA*, *hopQ*, and *babA*. Because Hardy was present in the population before the arrival of the cagPAI, the most likely explanation is that these alleles were originally imported from Hardy strains. Interestingly, while most Hardy strains so far are *cag*^−^, they encode three copies of VacA, as well as distinct versions of ∼ 20 OMPs, suggesting they might also be adapted to more closely interact with and modulate their host.

Strains with cagPAI deletions are frequently studied in the laboratory and can be useful in validating mechanistic hypotheses about what the island does^37,38^. However, our analysis implies that these knockouts are entirely distinct, genomically, from the persistent *cag*^−^ lineages that exist within natural populations. Therefore, if we wish to understand the ecological and evolutionary strategies used by *cag*^−^ and *cag^+^* strains, we need to compare these strains both in experimental populations and in mixed infections *in vivo*. Moreover, phenotypic assays may need to incorporate physiologically relevant gastric conditions to capture potential differences in the environmental adaptation and fitness of distinct *H. pylori* lineages, including *cag*^−^ and less-inflammatory strains. More broadly, our findings suggest that horizontally acquired pathogenicity islands should not be viewed simply as autonomous genetic modules, but as elements embedded within, and potentially driving the evolution of, genome-wide coadapted states. Quantifying these epistatic architectures provides a broader framework for investigating how bacterial genomes accommodate major evolutionary innovations across natural populations.

## Methods

### Genome collection

A total of 9,164 *Helicobacter* whole-genome sequences were used for bioinformatics analysis ^3^, which comprises 7,800 second-generation sequencing samples (incomplete genomes) and 1,364 third-generation sequencing samples (complete genomes). This dataset was from a previously published global collection of *H. pylori* genomes^3^, after excluding duplicate and low-quality strains. These strains include 9,159 *H. pylori* from different regions around the world, which were previously assigned to 15 geographic populations (designated hp) or less differentiated subpopulations (designated hsp), by chromosome painting **(Supplementary Table 1)**, as well as five *H. acinonychis* strains.

Genomes were assigned to *H. pylori* populations and classified as typical hpEurope or typical hspEAsia strains if they were assigned to the corresponding population and exhibited its characteristic genetic profile. Within these populations, redundant genomes were removed^3^, with redundancy defined as a pairwise SNP distance of <200. Low-quality assemblies were excluded if they contained >500 contigs, covered <70% of the *H. pylori* 26695 reference genome (NC_000915.1), or showed >10% non–*H. pylori* contamination as estimated by Kraken v.2.1.2^39^. After filtering, 1,425 typical hpEurope genomes and 1,179 typical hspEAsia genomes were retained for downstream analyses (**Supplementary Table 1**).

### Sequence alignment and genome variant calling

Single-nucleotide polymorphisms from the core genome (core SNPs) were called using an algorithm based on MUMmer v.3.20^40^, as previously described^3^. We first aligned each genome sequence of the entire dataset to the *H. pylori* 26695 reference strain using nucmer v.3.1^40^. Next, snp-sites v.2.5.2^41^ was used to call all variants from the whole-genome alignment obtained. Variants present in at least 99% of genomes were finally extracted using VCFtools v.0.1.17^42^. This yielded a total of 704,713 core SNPs for the entire dataset of 9,164 strains, 639,685 core SNPs among typical hpEurope genomes and 631,858 core SNPs among typical hspEAsia genomes. In parallel, the complete set of SNPs identified from the whole-genome alignments comprised 864,067 SNPs for typical hpEurope genomes and 833,634 SNPs for typical hspEAsia genomes.

### Phylogenetic analysis

To generate population phylogenetic trees, the cagPAI coadaptation tree and the cagPAI highly associated intergenic region tree, we used coding sequences that were aligned with reference strain 26695. Population phylogenetic trees are based on core SNPs, including the global, typical hpEurope (**shown in Fig. 1b**), typical hspEasia (**shown in Fig. 1d**). The *cag* coadaptation tree (**Fig. 6a**) was built to understand the coadaptation that took place when the cagPAI arrived in the *H. pylori* genome, based on 20 representative strains (**Supplementary Table 1**). These included four hpEurope strains (two *cag^+^* strains and two *cag*^−^ adapted strain), four hspAfrica1WAfrica strains (two *cag^+^* strains and two *cag*^−^ adapted strain), two hspEasia *cag^+^* strains, two hpSahul *cag^+^* strains and one *cag*^−^ strain, two hpAfrica2 *cag*^−^ strains, and five hardy strains (two *H. acinonychis* and three *H. pylori*). This tree was reconstructed using 30 representative SNPs (**Supplementary Table 4**) selected from 5,707 *cag* coadaption associated SNPs shared between typical hpEurope and the three hspEurope sub-populations **(Supplementary Table 2)**, cover 30 coadaptation blocks. The Hardy high-differentiated gene tree, and hardy low-differentiated gene tree (shown in **Fig. 6bc**) are based on the whole SNPs in the genes published previously **(Supplementary Table 6)**^3^. The *cag* coadaption associated intergenic region (**Supplementary Table 3**, *e.g*., HP0721|HP0722 etc.) tree is shown in **Extended Data Fig. 4.**

When looking at 32 specific *cag* coadaptation associated genes from the 5,707 *cag* coadaptation associated SNPs (**Supplementary Table 3**, *e.g*., *vacA*, *hopQ*, *sabA, etc*), gene sequences were first obtained from the individual strains annotation file then aligned using MAFFT (v.7.505, option --auto)^43^. The tree shown in **Extended Data Fig. 3** was built using SNPs from all coding sequences. In addition, gene presence and absence of these genes were analyzed, for particular gene CDS sequences, the sequences were separated into two types because some strains had two copies of the gene. The choice of copy type was based on the similarity between sequences^3^.

Using the various alignments of nucleotide sequences, maximum-likelihood phylogenetic trees were constructed with FastTree (v.2.1.10,-nt)^44^ to infer evolutionary relationships among the sequences. Plotting was done using the R package ggtree v.3.2.1 ^45^. The population-level trees were built via a neighbourjoining algorithm (R package ape, function nj), using a matrix of the average distance between genomes. Distances between strains were calculated using the dist.dna function (option model = ‘raw’) from the ape R package.

### Genome-wide association study and identification of differentiated loci

Genome-wide association studies (GWAS) were performed using *cagF* status as the phenotype in typical hpEurope and typical hspEAsia. Presence/absence of *cagF* (HP0543) was coded as 1 and 0, respectively. SNPs were extracted from whole-genome alignments using snp-sites (v2.5.1) ^41^ and converted into VCF format. Multiallelic variants were collapsed into pseudo-biallelic categories (major versus minor allele) and analyzed using pyseer (v1.3.6)^46^. Population structure was controlled for using multidimensional scaling (MDS) components derived from pairwise genetic distances and supplied to pyseer through the ‘--distances’ option. Variants with allele frequencies below 2% or above 98% were excluded using --min-af 0.02 and --max-af 0.98, respectively.

For typical hpEurope, 682 *cagF*-negative (*cagF*^−^) strains were compared with 743 *cagF*-positive (*cagF*^+^) strains. SNPs showing strong association with cag status were initially defined as those with –log_10_(*P*) ≥ 5. A more stringent threshold of –log_10_(*P*) > 25 was subsequently applied, and variants located within the cagPAI were excluded, resulting in 15,976 candidate SNPs from 407,886 tested variants. Because modest population structure was observed among the three subpopulations, separate GWAS analyses were additionally performed within each subpopulation. Significance thresholds were set at −log_10_(*P*) ≥ 20, 5 and 10 for hspSWEurope, hspNEurope and hspEurasia, respectively. Under the null hypothesis that the three subpopulation-specific GWAS were independent, the expected three-way overlap was calculated as *n*_hspSWEurope_ × *n*_hspNEurope_ × *n*_hspEurasia_ / *N*^2^, where *N* = 325,592 was the number of variants tested in all three analyses, and *n*_hspSWEurope_ = 18,492, *n*_hspNEurope_ = 7,610 and *n*_hspEurasia_ = 14,620 were the numbers of significant variants **among these shared tested variants** in hspSWEurope, hspNEurope and hspEurasia, respectively. This yielded an expected three-way overlap of approximately 20 variants. To reduce false-positive associations arising from population stratification, only variants that exceeded the respective significance thresholds in all three subpopulation-specific GWAS and in the combined typical hpEurope GWAS were retained. This procedure yielded 5,707 *cagF*-associated SNPs shared across the overall hpEurope population and subpopulations **(Supplementary Table 2)**.

For typical hspEAsia (n = 1,179), 50 *cagF*^−^ strains were compared with 1,129 *cagF*^+^ strains. Using a significance threshold of −log_10_(*P*) ≥ 5 and excluding variants located within the cagPAI region, 15 SNPs were identified from 338,527 tested variants as significantly associated with *cag* status (**Supplementary Table 5**). To further investigate genetic signatures associated with the OMP configuration A lineage in hspEAsia, an additional GWAS was performed by comparing the strains carrying OMP Configuration A with strains carrying all other patterns. Using significance threshold (−log_10_(*P*) ≥ 5) and excluding variants located within the cagPAI region, 800 SNPs were identified from 338,527 tested variants as significantly associated with the Configuration A lineage. All of these variants are located in a block of genes (HP0721–HP0726; 773,625–782,101 bp in *H. pylori* 26695), as shown in **Supplementary Fig. 5a**.

Among the 5,707 *cag* status-associated SNPs identified in hpEurope, 50 annotated loci were initially considered (**Supplementary Table 3**), comprising 32 single-gene loci and 18 intergenic or gene-boundary loci. Physically connected loci were merged using the strain 26695 annotation. Genes linked by an intergenic annotation, or adjacent in the reference genome and represented in the candidate SNP set, were assigned to the same unit, with connectivity extended transitively across consecutive regions. This procedure identified 10 multigene blocks and 20 unmerged gene or intergenic loci, yielding 30 coadaptation units. Loci were labelled as BxHy, where x denotes the linkage block and y denotes the rank of the locus by −log10(*P*) within that block. To reduce redundancy arising from linkage disequilibrium, the most strongly associated SNP from each unit was retained, yielding 30 representative SNPs (B1H1–B30H1) for coadaptation-score calculation, phylogenetic reconstruction and haplotype analysis (**Supplementary Table 4**).

### Coadaptation score calculation and global coadaptation landscape analysis

PRS was defined as a single-value summary of an individual strain’s genome-wide genetic contribution to the phenotype, calculated as the weighted sum of genotype values using effect size estimates (β coefficients) derived from GWAS summary statistics^14^.

The total cagPAI coadaptation score was calculated using hpEurope alleles at the 30 representative SNPs (B1H1–B30H1), with the SNP showing the highest −log₁₀(*P*) value selected from each differentiated block (**Supplementary Table 4**). PRS values were calculated by summing alleles at these loci, weighted by their direction of differentiation^14^. For each strain, the PRS was calculated as:

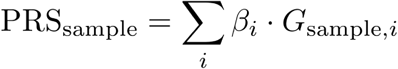

where G*_sample,i_*is the genotype of a given strain at locus i, encoded as 0 (reference allele) or 1 (alternative allele), and β*_i_* is the GWAS-derived effect size for that locus.

In addition to the total cagPAI coadaptation score, we calculated complementary locus-specific and locus-excluded scores. For the *vacA* locus, a *vacA*-only coadaptation score was calculated from the 573 variants within the 5,707-SNP set that mapped to positions 938,415–942,287 bp in *H. pylori* 26695. A complementary non-*vacA* coadaptation score was calculated using the same top-one-SNP-per-block scheme (B1H1–B30H1) as for the total score, but after excluding selected score variants mapping to the *vacA* locus. We applied the same approach to the alginate block, hereafter referred to as the *patA* locus. The *patA*-locus-only score was calculated from the 1,173 variants in the 5,707-SNP set mapping to the 4,792-bp interval from positions 907,503 to 912,294 bp, spanning the HP0854–HP0858 region and intervening intergenic sequences. The corresponding *patA*-locus-excluded score was calculated by removing the selected SNP within this interval from the total score. We further examined associations between locus-excluded coadaptation PRS and OMP configurations (**Fig. 2**), *vacA* genotypes and *patA*-locus genotypes (**Fig. 5d**).

### Stage-specific genome-wide association analyses of cagPAI coadaptation

To identify genetic variation associated with successive stages of coadaptation following gain or loss of the cagPAI, typical hpEurope strains were divided into four datasets according to *cag* status and cagPAI coadaptation polygenic risk score (PRS; **Fig. 5a**). Thresholds were selected empirically from the PRS distributions stratified by *cag* status to define four non-overlapping groups representing the principal coadaptation gradient. The thresholds of-30 and 14 were close to the medians of the *cag*^−^ and *cag*^+^ distributions, respectively, whereas-17 was obtained by rounding the Youden’s J-optimal cut-off of −16.795 between the intermediate *cag*^−^ and *cag*^+^ groups. This scheme retained sufficiently sized groups at successive positions along the PRS gradient while excluding strains with discordant *cag* status and PRS values. The four datasets comprised strongly coadapted *cag*^−^ strains with PRS <-30 (dataset 1; *n* = 345); less strongly coadapted *cag*^−^ strains with-30 ≤ PRS <-17 (dataset 2; *n* = 234); less strongly coadapted *cag*^+^ strains with-17 ≤ PRS ≤ 14 (dataset 3; *n* = 351); and strongly coadapted *cag*^+^ strains with PRS > 14 (dataset 4; *n* = 368).

To investigate stage-specific genetic associations during coadaptation to *cag* status, the four PRS- and *cag*-status-defined datasets were used to perform three adjacent GWAS comparisons. Dataset 1 versus dataset 2 compared *cag*^−^ strains with low and intermediate-low PRS values; dataset 2 versus dataset 3 compared *cag*^−^ and *cag*^+^ strains at intermediate PRS ranges; and dataset 3 versus dataset 4 compared *cag*^+^ strains with intermediate and high PRS values. These three comparisons were designed to identify variants associated with successive phases of coadaptation to *cag* status.

All variants tested in each stage-specific association analysis were assigned to genes or intergenic regions using the strain 26695 reference annotation. For each annotated region and each comparison, the strongest association signal was defined as the maximum −log₁₀(P) across all variants assigned to that region. The three region-level maximum values were summed to obtain an accumulated association score, and the 50 regions with the highest accumulated scores were selected. These regions were subsequently matched at the gene or intergenic-region level to the previously defined summary of 5,707 *cag*-associated variants. Regions represented in this summary were retained for visualization and ordered according to their maximum −log_10_(*P*) in the original *cag*^+^ versus *cag*^−^ association analysis (**Fig. 5b**). Thus, the stage-specific association analyses were not restricted to the 5,707 variants; the previously defined set was used only for region-level matching and ordering in the final visualization.

### Classification of outer membrane protein configurations

We defined the HP0721–HP0726 region as a contiguous genomic block encompassing the HP0723 downstream locus, which underlies the strongest GWAS signal. To investigate structural variation at this locus, *e.g*., position SNP 776,433 located 48 bp downstream of HP0723 (*ansB*, encoding asparaginase), we analysed this genomic interval across the global population with the gene annotation (.gff) files generated by Prokka (v1.14)^47^. The *ansB* gene is located on the reverse strand of reference strain 26695; therefore, genomic coordinates were interpreted such that decreasing positions correspond to transcriptional downstream direction. Within this block, isolates in which the region was contained within a single contig were classified according to the identity of the first annotated gene downstream of *ansB*.

Herein, five discrete outer membrane protein (OMP) configurations were identified: Configuration A (HP0721 adjacent to HP0723), Configuration B (HP0725/*sabA*), Configuration C (HP0722/*sabB*), Configuration D (HP1177/*hopQ*), and Configuration E (all remaining downstream gene arrangements, e.g., HP0697). For genomes in which this region was fragmented across multiple contigs and local gene order could not be directly resolved, isolates were assigned to Configuration F. As shown in **Supplementary Fig. S4**, haplotype analysis revealed that each OMP configuration possesses a characteristic set of SNPs within the HP0721–HP0726 block.

OMP configurations were scanned across global populations and stratified by *cag* status. These structural classes were subsequently integrated with genome-wide total gene coadaptation score, *vacA* coadaptation score and other gene coadaptation score, computed per isolate as described above, to assess variation in coadaptation across OMP-defined structural backgrounds and population strata.

### Amino-acid-based *vacA* typing and association with cagPAI coadaptation

VacA proteins were classified using amino-acid residues homologous to Ala33 and Arg496 of the strain 26695 s1/i1/m1 reference protein. VacA candidate sequences were placed into a reference-anchored amino-acid alignment containing the strain 26695 sequence and representative s1/m1, s1/m2 and s2/m2 sequences. Alignment columns corresponding to residues 33 and 496 of strain 26695 were identified from the ungapped reference sequence. At the position homologous to residue 33, alanine and serine were classified as s1 and s2, respectively. At the position homologous to residue 496, arginine and lysine were classified as m1 and m2, respectively. Other amino acids were classified as ambiguous, whereas gaps, stop codons and unknown residues were treated as missing.

For strains containing multiple *vacA* candidates, all full-length copies with an aligned coverage of at least 1,100 amino acids were considered. If no full-length copy was available, candidates retained by the previous fragment-reconstruction procedure were used. Consistent calls across copies were assigned to the corresponding s or m type; conflicting recognized residues were retained as mixed calls. The combined amino-acid-based types were summarized as s1/m1, s1/m2, s2/m1, s2/m2 or other/ambiguous.

To examine cagPAI coadaptation-associated variation at *vacA*, variant-level association signals were extracted across the HP0886–HP0887 intergenic region and the complete *vacA* coding sequence in the strain 26695 reference genome (positions 938,191–942,287). We focused on the two comparisons representing progressive coadaptation within each *cag* state: dataset 1 versus dataset 2 for *cag*^−^ strains and dataset 3 versus dataset 4 for *cag*^+^ strains. Association strength was expressed as −log_10_(*P*). Signals were displayed at their strain 26695 genomic coordinates. The HP0886–HP0887 intergenic region, the *vacA* s–region typing interval, the i-region B and C sequence clusters and the m2-associated 21-amino-acid insertion site were annotated using the strain 26695 reference sequence. For each comparison, the three variants with the largest −log_10_(*P*) were labelled with their genomic coordinates and corresponding amino-acid positions in the strain 26695 VacA protein.

To examine the relationship between *vacA* type and cagPAI coadaptation while avoiding circularity, cagPAI coadaptation PRS values were recalculated after excluding variants within the *vacA* locus. Strains were classified as *cag*^+^ or *cag*^−^ according to the presence or absence, respectively, of HP0543. The distributions of the *vacA*-excluded PRS were displayed as mirrored stacked histograms, with *cag*^+^ and *cag*^−^ strains plotted above and below the horizontal axis, respectively. Mean PRS values were calculated separately for each combination of *vacA* type and *cag* status and displayed only for groups containing at least ten strains. Analyses were performed for the global dataset and four geographical groupings: Europe (hspSWEurope, hspNEurope and hspEurasia), Asia (hspEAsia, hpAsia2, hspNorthAsia and hspSAsia), Africa (hpAfrica2, hpNEAfrica, hspCNEAfrica, hspAfrica1SAfrica and hspAfrica1WAfrica) and other lineages (hpSahul, hspUral, hspIndigenousAmerica and Hardy strains).

### Haplotype analysis

We classified nucleotide states at 5,707 cagPAI-associated SNPs using 1,425 typical hpEurope strains as the reference cohort. At each site, the frequency of each nucleotide (A, C, G or T) was calculated separately among *cagF*^+^ and *cagF*^−^ strains. A nucleotide was classified as *cagF*^+^-associated or *cagF*^−^-associated according to the group in which it was more frequent; ties were classified as other alleles. Gaps and ambiguous or missing characters were classified as absent. These definitions were then applied to 9,164 global strains.

Genomic blocks were defined from the physical arrangement of genes in the strain 26695 annotation. Genes linked by an intergenic annotation, or adjacent in the GFF and represented in the candidate SNP set, were assigned to the same block, with connectivity extended transitively across consecutive regions. Within each multigene block or unmerged gene or intergenic region, SNPs were ranked by likelihood-ratio-test P value, absolute effect size and genomic position. The most strongly associated SNP from each unit was retained, yielding 30 representative SNPs (B1H1–B30H1) for haplotype visualization.

Sequence completeness was assessed independently of SNP state. Regions with ≤10% nucleotide coverage were classified as absent, those with >10% but <90% coverage as partial or truncated, and those with ≥90% coverage as present. Absent regions, gaps within partial regions and local deletions within otherwise complete regions were distinguished from observed nucleotides. The resulting 30-SNP haplotypes were examined across the global dataset and a curated set of 20 representative isolates to investigate *cag*-associated population structure and ancestry.

For locus-specific analyses, *vacA* haplotypes were reconstructed from 573 *cag*-associated SNPs within the strain 26695 *vacA* coding sequence. Haplotype analysis was also performed across the 4,792 bp *patA*-associated region (positions 907,503–912,294), encompassing the HP0854–HP0855 intergenic region and HP0855–HP0858. Haplotypes across the HP0721–HP0726 outer-membrane-protein region were reconstructed from 1,336 *cag*-associated SNPs. Structural coverage of the 8,477-bp HP0721–HP0726 interval (positions 773,625–782,101) was additionally estimated from the reference-anchored global alignment and classified as absent, partial or present using the same 10% and 90% coverage thresholds.

### Chromosome painting analysis

Two complementary ancestry inference analyses were performed on the global dataset using ChromoPainter v.2^48,49^. To assign a population to the final *H. pylori* dataset, we first defined a 15–component model for the present study. This model was generated based on the 19 populations/subpopulations in previous reports^3^, by merging hspAfrica1LatinAmerica with hspAfrica1WAfrica, hspSWEuropeLatinAm with hspSWEurope, hspSiberia with hspNorthAsia, and hspIndigenousNAmerica together with hspIndigenousSAmerica into hspIndigenousAmerica. In addition, a South Asian component (hspSAsia) was incorporated into the model. The final 15 ancestry components comprised hpAfrica2, hspAfrica1WAfrica, hspAfrica1SAfrica, hpNEAfrica, hspCNEAfrica, hspNEurope, hspSWEurope, hspEurasia, hpAsia2, hpSahul, hspUral, hspNorthAsia, hspIndigenousAmerica, hspEAsia, and hspSAsia. To validate this reference donor panel, We confirmed the population structure by running fineSTRUCTURE^49^ based on haplotype data from core SNPs, using 200,000 iterations of burn-in and Markov chain Monte Carlo. This subset was then used as a donor panel to paint each sample across the entire dataset using ChromoPainter v.2. Genomes were labelled with a population based on the largest fraction of their genome painted by a population.

Moreover, we selected representative donor strains from five ancestral populations described previously^48^, including hspEAsia (n = 20), hpAsia2 (n = 20), hspCNEAfrica (n = 10), hspENEAfrica (n = 10), and hspAfrica1WAfrica (n = 10). Core SNPs were partitioned into two genomic subsets corresponding to loci within the cagPAI and the remainder of the genome. Chromosome painting was performed separately for each partition using ChromoPainter v.2, estimating site-by-site copying probabilities from each donor population. Ancestry profiles were then summarized and visualized for recipient strains grouped by subpopulation.

### Core gene multilocus sequence typing (cgMLST)

To investigate the existence of clonal relationships in *H. pylori*, we estimated the total number of identical loci shared among strains from the global dataset by performing a cgMLST as implemented by chewBBACA software v2.8.5^50^. The Schema was accessed from the HpGP dataset as previously reported^2^, and then applied to the analysis of the global dataset. chewBBACA uses a gene-by-gene method to compare coding sequences and assign alleles based on a BLAST Score Ratio (BSR). We first used Prodigal^51^ v2.6.3, including the option-t to create a training file from the assembled version of the 26695 *H. pylori* reference strain resequenced as part of the HpGP dataset. Then, the “CreateSchema” module of chewBBACA was applied to the 1011 HpGP genomes and the Prodigal training file to estimate a whole-genome MLST (wgMLST) scheme. The 3943 wgMLST genes were then compared with the “AlleleCall” module, using the default BSR threshold of 0.6. A total of 867 genes identified as paralogs were removed from the wgMLST using the “RemoveGenes” module, reducing the scheme to 3076 loci. We then used the “ExtractCgMLST” module to create a cgMLST with all loci present in more than 95 percent of strains (--t 0.95), obtaining a total of 981,110 alleles for 1040 loci, an average of 943 different alleles per locus.

We used the cgMLST allelic profile to calculate pairwise distances with GrapeTree^52^ v1.5.0, running it in “--wgMLST” mode with the “distance” method (-method distance) while ignoring missing data (--missing 0). Subsequently, we clustered samples based on distances, considering those that shared over 90% identical alleles (cgMLST distance = 0.90), which represent the clonal groups.

### Dating of the hspEAsia *cag^−^* adapted deep clone

A core gene alignment of the 25 Japan isolates (**Supplementary Table 1**) with cagPAI coadaptation score less than-27 was generated with panaroo^53^ using the settings described above. A phylogenetic tree was computed using PhyML^54^ v3.1and input to ClonalFrameML^17^ v1.12, executed using default parameters. Node ages were determined using the R BactDating^18^ package, using 100,000 Markov chain Monte Carlo iterations and a fixed mutation rate of 1.38 × 10^−5^ per site per year (updateMu = FALSE), as has previously been estimated^55^. MCMC convergence was formally confirmed through inspection of parameter traces and verified by effective sample sizes (*ESS* > 400 across all valid parameters; median *ESS* = 900. 94; root node *ESS* = 1000. 00), well exceeding the standard convergence threshold of 200. Under this model, the most recent common ancestor was dated to 1812.6 (95% highest posterior density interval: 1755.3–1862.0).

### Evaluating *cag*PAI coadaptation dynamics and natural selection against genomic controls

Pairwise divergence at any set of loci is expected to increase as strains diverge across the genome. Thus, elevated divergence at the 30 representative SNPs (B1H1–B30H1) used to calculate the cagPAI coadaptation score could simply reflect genome-wide phylogenetic divergence rather than natural selection acting specifically on these loci. We therefore asked whether allelic configurations at these 30 loci differed across *cagF* status categories and populations when comparing strain pairs with equivalent levels of genome-wide core-SNP divergence.

Strain pairs within each population were stratified by *cagF* status (*cagF*⁺/*cagF*⁺, *cagF*⁻*/cagF*⁺, and *cagF*⁻/*cagF*⁻) and partitioned into left-closed, right-open 5,000-SNP windows of genome-wide core divergence spanning 0 to 65,000 SNPs. Within each window, we first evaluated the mean total allele distance across the 30 representative SNPs (B1H1–B30H1), treating single-nucleotide variants and insertion/deletion events (gaps) as distinct allele states (**Fig. 3a**). To distinguish true nucleotide substitution from gap-related differences, we next calculated the per-pair mismatch proportion strictly at jointly callable A/C/G/T positions and averaged these proportions within each window (**Fig. 3b**).

These observed values were compared with 100 independently sampled control sets, each comprising 30 loci drawn from the 99% core genome. To minimize confounding by *cag*-associated variation, control candidates were sampled after excluding all 5,707 *cagF*-associated positions, the complete regions containing the 30 representative SNPs (B1H1–B30H1), and the cagPAI together with its 50-kb flanking regions. Control loci were matched to the representative SNPs based on the global five-state (A/C/G/T/gap) allele frequency spectrum across all 9,164 strains where possible, or otherwise on the frequency of the most common allele state.

Windows containing fewer than 10 strain pairs were excluded from curve fitting. Trajectories across retained windows were smoothed using robust locally weighted scatterplot smoothing (two robustifying iterations) followed by shape-preserving piecewise cubic Hermite interpolating polynomials evaluated over a uniform 261-point grid. For **Fig. 3b**, this smoothing pipeline was applied independently across all 100 control sets to derive the pointwise mean and empirical 2.5th–97.5th percentile envelope.

### Quantification of pairwise SNP distances and comparative analysis

To evaluate the genomic divergence between the cagPAI and the background genome, a globally representative dataset of *H. pylori* isolates (n = 2,381, **Supplementary Table 7**) was constructed. Isolates were selected based on the complete presence of all 28 genes within the cagPAI, defined population lineage assignment, and high sequence quality across the cagPAI region (genomic coordinates 537,328–583,481, requiring a sequence coverage cutoff >75%). Multi-FASTA alignments were generated for both the trimmed cagPAI region and whole-genome sequences. Pairwise SNP distances between the cagPAI region and whole-genome sequences (**shown in Figure S8b**) were quantified using snp-dists v0.8.2 (https://github.com/tseemann/snp-dists). For every pair of isolates, cagPAI SNP distances were plotted against full-genome SNP distances. Pairs were stratified and color-coded according to population assignment: pairs sharing the same defined population lineage were assigned their respective population colors, whereas pairs spanning different populations or possessing ambiguous lineage classifications were categorized as Mixed and visualized in grey. Data processing and visualization were performed using Python (v3.10) with the pandas (v2.0), seaborn (v0.12), and matplotlib (v3.7) packages.

## Notes

### Competing Interest Statement

The authors have declared no competing interest.

