## Supplementary Figs. 1-11 for "History of coadaptation to the *cag* pathogenicity island in *Helicobacter pylori*"

Supplementary materials:
**Supplementary Figs.** 1-11

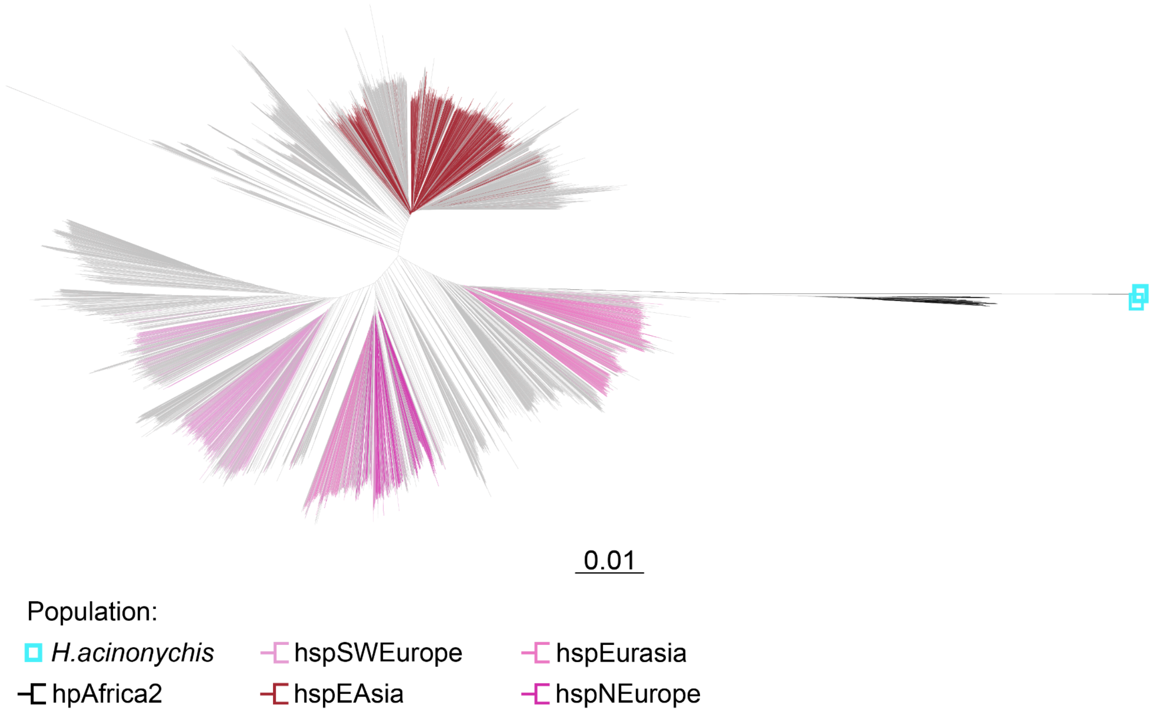


**Supplementary Fig. 1.** Phylogenetic tree for 9,159 globally sampled *H. pylori* and five *H. acinonychis* genomes.

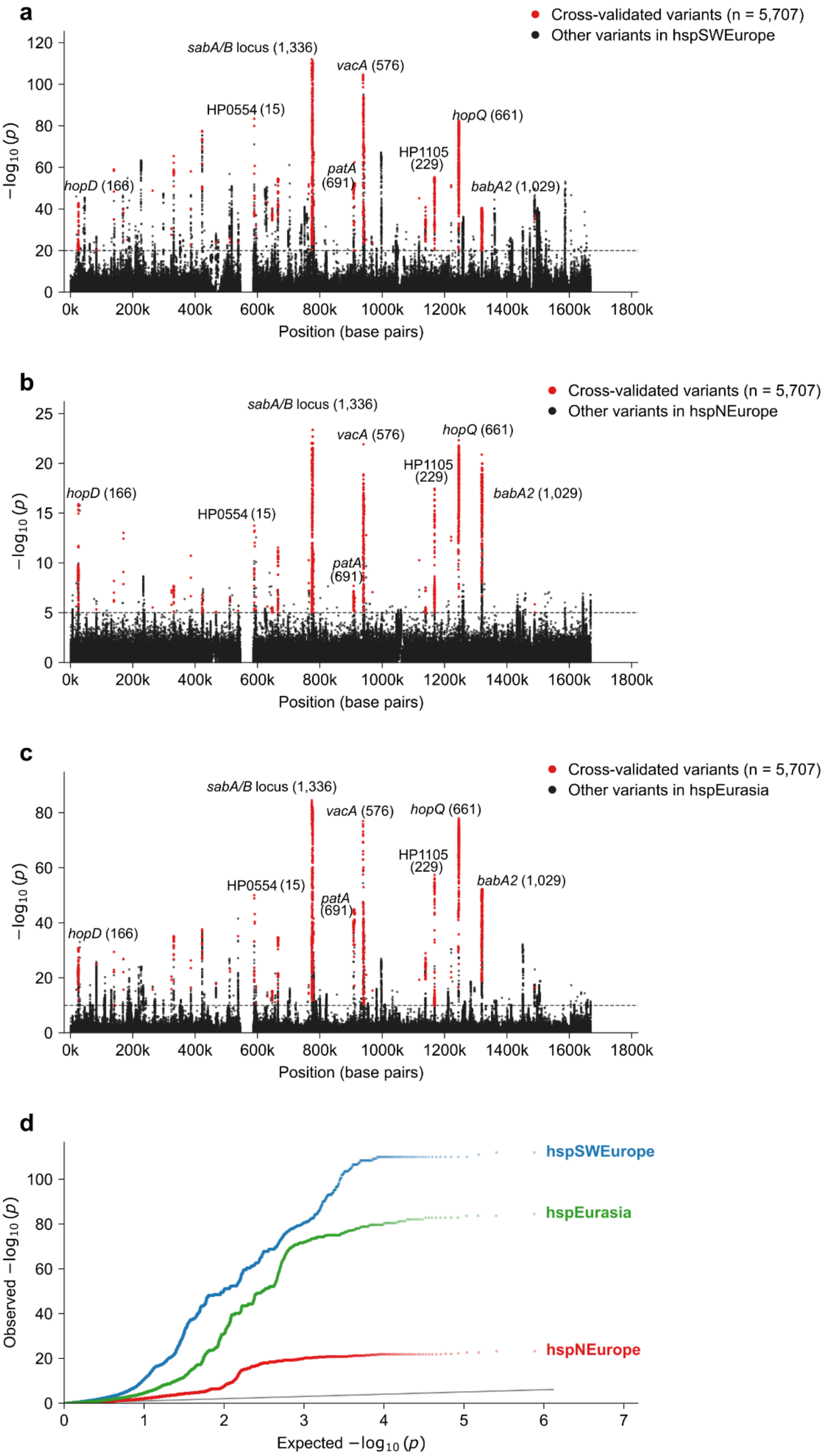


**Supplementary Fig. 2. Manhattan plot from GWAS analysis of *cagF*$^{+}$ versus *cagF*$^{-}$ strains within three hpEurope subpopulations. a, hspSWEurope; b, hspNEurope; and c, hspEurasia.** Sub-population-specific significance cutoffs were set at $-\log_{10}(P)$ values of 20, 5, and 10, respectively. **d,** Quantile–quantile plots of *cag* status GWAS results in three European subpopulations. Observed and expected $-\log_{10}(P)$ are shown for GWAS comparing *cagF*$^+$ and *cagF*$^-$ strains in hspSWEurope (blue; $\lambda_{GC} = 7.22$), hspNEurope (red; $\lambda_{GC} = 2.15$) and hspEurasia (green; $\lambda_{GC} = 3.59$). The grey diagonal indicates the null expectation. The cagPAI region and it's 3 kb flanking regions (*H. pylori* 26695 positions 541,336–588,350) were excluded, whereas the ten PRS-associated genomic blocks were retained.

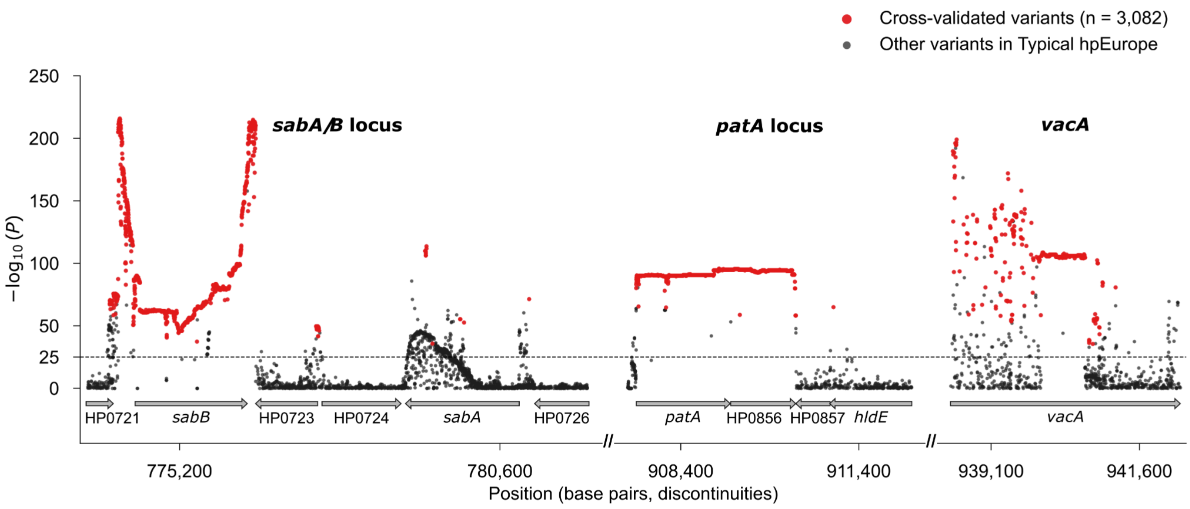


**Supplementary Fig. 3. Regional association signals at three loci associated with *cag* status in hpEurope.** Regional Manhattan plot showing variants from the typical hpEurope *cag*$^+$ versus *cag*$^-$ GWAS across three genomic regions. HP0721–HP0726, the *patA*-associated block comprising the HP0854–HP0855 intergenic region and HP0855–HP0858, and *vacA* (HP0887). Red points indicate 3,082 variants within these regions that were replicated in all three European subpopulation-specific GWAS (hspSWEurope, hspNEurope and hspEurasia), whereas black points indicate the remaining variants. The horizontal dashed line denotes the significance cutoff at $-\log_{10}(P) = 25$. Gene positions and orientations in the *H. pylori* 26695 reference genome are shown by arrows below the plot. The three regions are displayed on a compressed, discontinuous genomic axis, with double slashes (//) indicating coordinate discontinuities; distances between regions are not drawn to genomic scale.

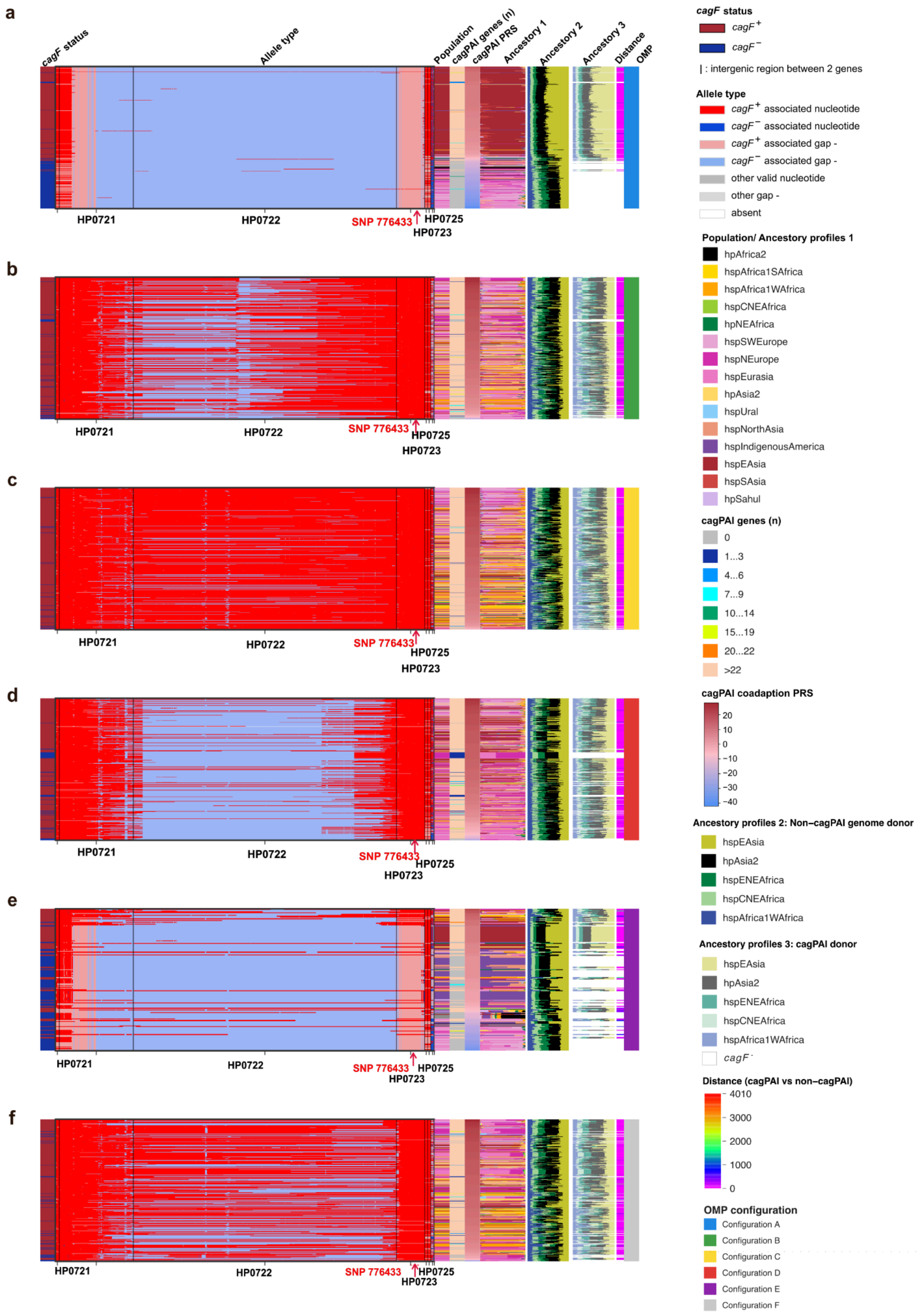


**Supplementary Fig. 4. Haplotype structure for six outer membrane protein (OMP)**

**configurations at the HP721–HP0726 locus.** Panels a-f correspond to OMP configurations A-F (see Main **Fig. 1e**). Column (C) contents are as follows: 1) *cagF* (HP0543) status; 2) allele type; 3) geographic population; 4) number of cagPAI genes: 5) cagPAI coadaption PRS; 6) ancestry profile (15-component model); 7) non-cagPAI ancestry donor contribution (5-component model), 8) cagPAI ancestry donor contribution (5-component model); 9) distance between cagPAI/non-cagPAI ancestry donor contributions in *cagF*$^+$ strains; 10) OMP Configuration. Strains are ordered by cagPAI coadaption PRS.

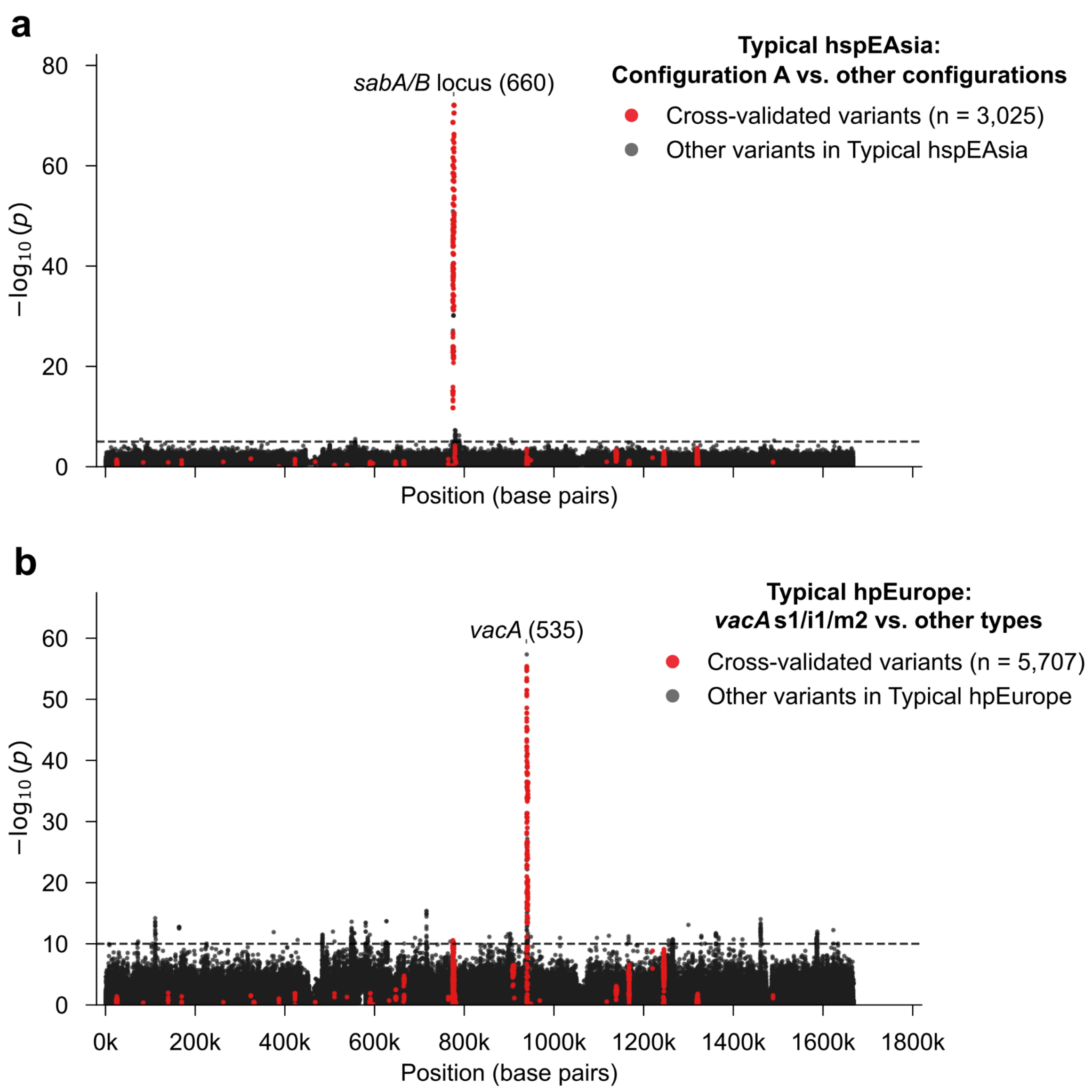


**Supplementary Fig. 5. Genome-wide association signals associated with Configuration A and the *vacA* s1/i1/m2 genotype.** **a**, Manhattan plot comparing typical hspEAsia strains carrying Configuration A with strains carrying other configurations. **b**, Manhattan plot comparing typical hpEurope strains carrying the *vacA* s1/i1/m2 genotype with strains carrying other *vacA* types. Black points denote other tested variants, whereas red points indicate variants overlapping the predefined set of 5,707 crossed variants. Horizontal dashed lines indicate the significance thresholds of $–\log_{10}(P) = 5$ in panel **a** and $–\log_{10}(P) = 10$ in panel **b**. Only the strongest significant peak in each analysis is annotated. Numbers in parentheses indicate the number of significant crossed variants assigned to the corresponding region. *sabA/B* locus: combined HP0721–HP0725 region.

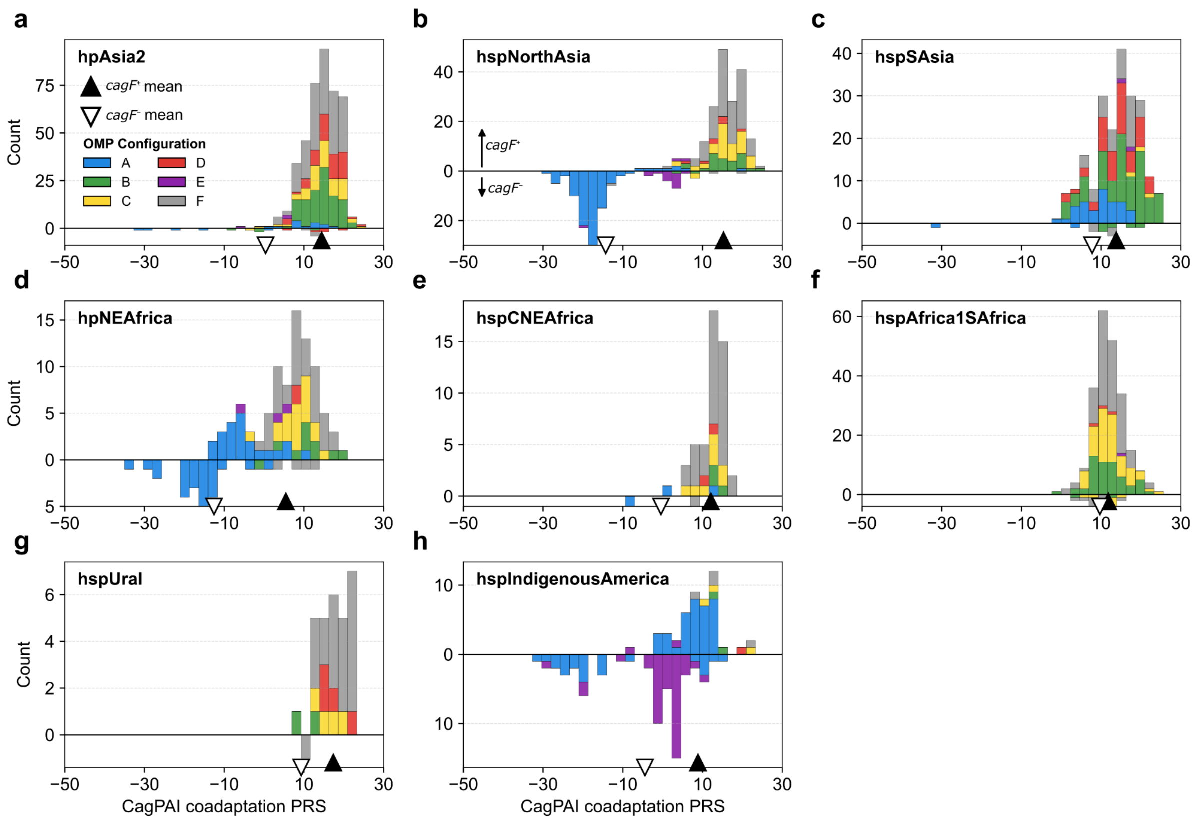


**Supplementary Fig. 6. Distribution of the cagPAI coadaptation PRS across the remaining global *Helicobacter pylori* populations.** **a**, hpAsia2; **b**, hspNorthAsia; **c**, hspSAsia; **d**, hpNEAfrica; **e**, hspCNEAfrica; **f**, hspAfrica1SAfrica; **g**, hspUral; **h**, hspIndigenousAmerica. Mirrored stacked bars show the distribution of the cagPAI coadaptation PRS, with *cagF*$^{+}$ strains above and *cagF*$^{-}$ strains below the horizontal axis. The x axis indicates the cagPAI coadaptation PRS and the y axis indicates strain count. Filled black upward triangles and open downward triangles mark the mean PRS of *cagF*$^{+}$ and *cagF*$^{-}$ strains, respectively. Colours denote OMP Configurations A–F, as defined in **Fig. 1e**; light grey indicates unassigned configurations.

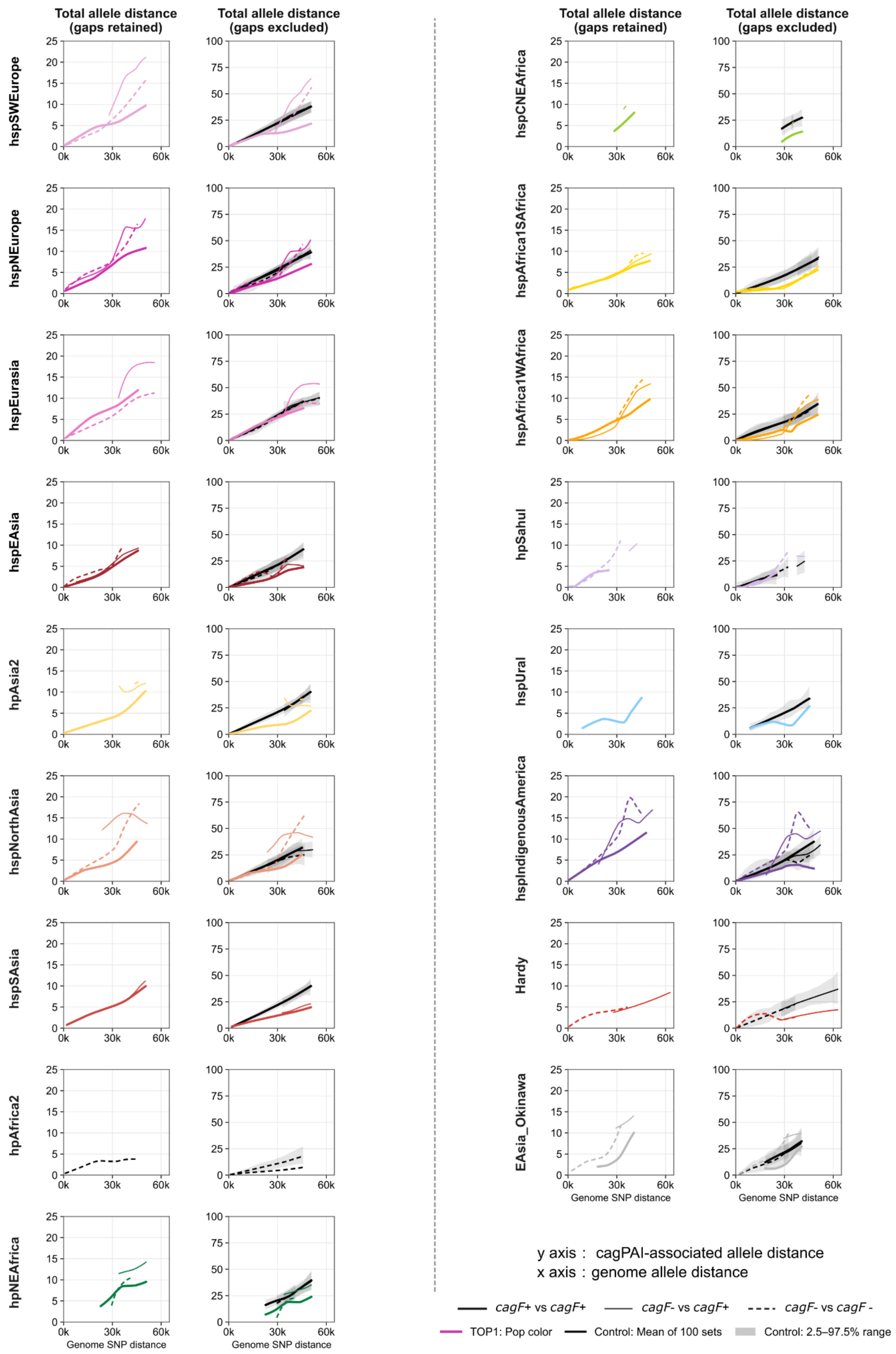
Total allele distance (gaps retained)
Total allele distance (gaps excluded)
hspSWEurope
hspNEurope
hspEurasia
hspEAsia
hpAsia2
hspNorthAsia
hspSAsia
hpAfrica2
hpNEAfrica
hspCNEAfrica
hspAfrica1SAfrica
hspAfrica1WAfrica
hspSahul
hspUral
hspIndigenousAmerica
Hardy
EAsia_Okinawa
Genome SNP distance
y axis : cagPAI-associated allele distance
x axis : genome allele distance
cagF+ vs cagF+
cagF- vs cagF+
cagF- vs cagF -
TOP1: Pop color
Control: Mean of 100 sets
Control: 2.5–97.5% range

**Supplementary Fig. 7. Dynamic, population-specific divergence at 30 *cag*PAI-associated H1 loci across global populations.** Panels display pairwise divergence trajectories across global *H. pylori* populations and defined subgroups, arranged in two paired-column blocks separated by the central dashed line (populations progress top-to-bottom in the left columns, followed by the right columns. For all panels, the x-axis denotes pairwise whole-genome core-SNP distance grouped into 5,000-SNP windows (spanning 0 to 65,000 SNPs). The y-axis denotes *cag*PAI-associated allele divergence across the 30 block-representative Top1 loci under two filtering criteria: **left columns** (columns 1 and 3) show the mean total allele distance with all variant types retained (A, C, G, T, and indel gaps treated as distinct states; range 0–25); **right columns** (columns 2 and 4) show the mean per-pair nucleotide mismatch proportion (%), calculated strictly across jointly callable A/C/G/T sites with gap-involved positions excluded separately for each strain pair (range 0–100%). Curve width and style denote *cagF* pairing status: thick solid lines, *cagF*$^{+}$/*cagF*$^{+}$; thin lines, mixed *cagF*$^{+}$/*cagF*$^{-}$; and dashed lines, *cagF*$^{-}$/*cagF*$^{-}$. Coloured trajectories represent the 30 Top1 loci using population-specific colours. Solid black curves and shaded grey envelopes depict the pointwise mean and empirical 2.5th–97.5th percentile range across 100 region-excluded, globally frequency-matched core-genome control sets. Fitted trajectories summarize 5,000-SNP windows containing at least 10 strain pairs per class.

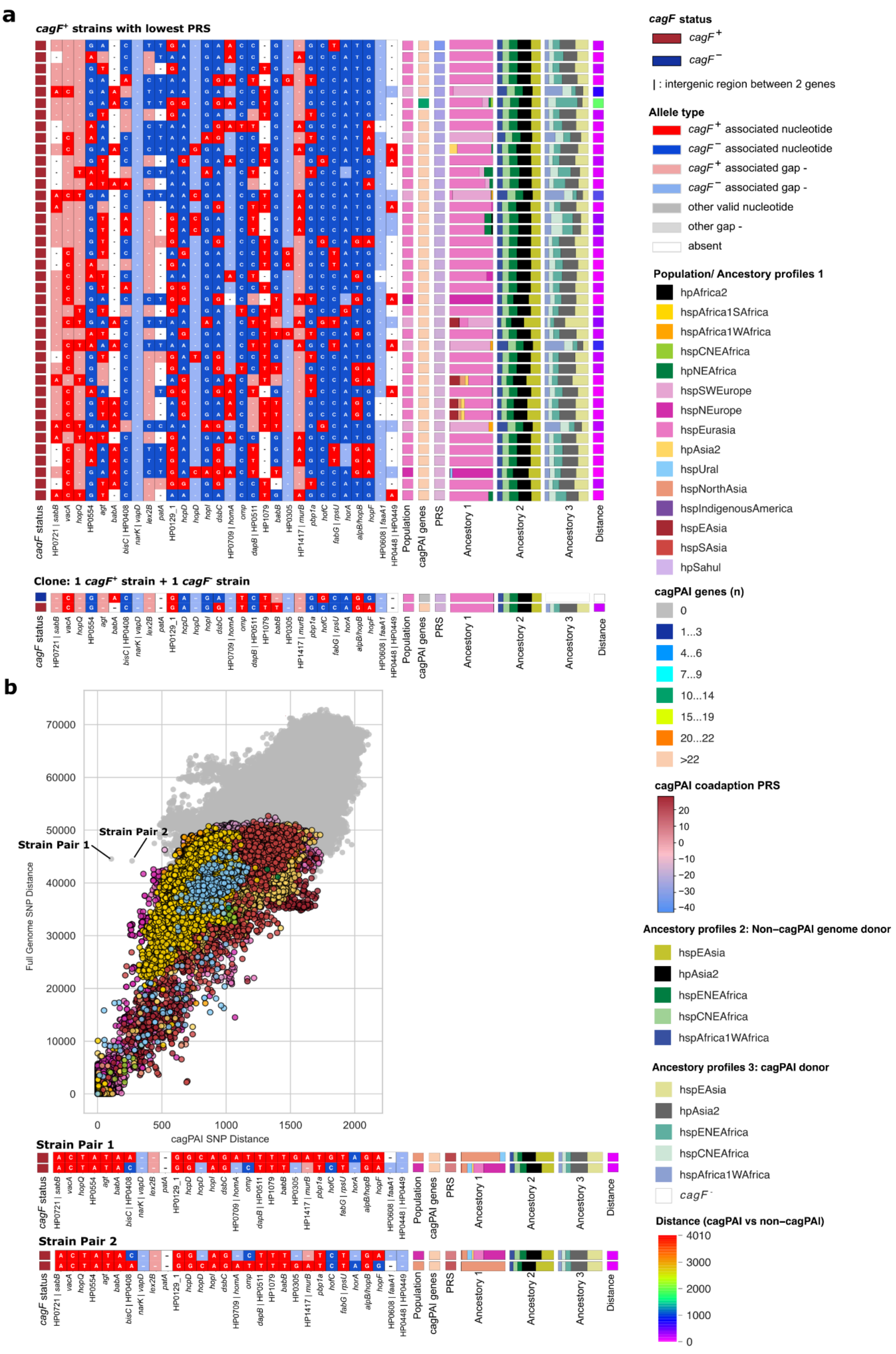


**Supplementary Fig. 8. a**, cagPAI coadaptation associated Haplotype structure across 40 *cag*+ hpEurope isolates (PRS < -15.85) and their clonal relationships with related *cag*- strains. **b**, cagPAI coadaptation associated Haplotype structure across three closely related Russian isolates (2 pairs). These 2 pairs of strains have a short Pairwise SNP distance of the cagPAI region (x-axis; positions 537,328–583,481, comprising 28 genes) against the whole-genome sequence (y-axis). For the scatter plot, each point represents a pair of isolates. Points are color-coded by population lineage when both isolates belong to the same population (according to the color legend); strain pairs belonging to different populations or unidentified groups are indicated in grey (Mixed). cagPAI coadaptation associated Haplotype structure of each pair of strains were shown below.

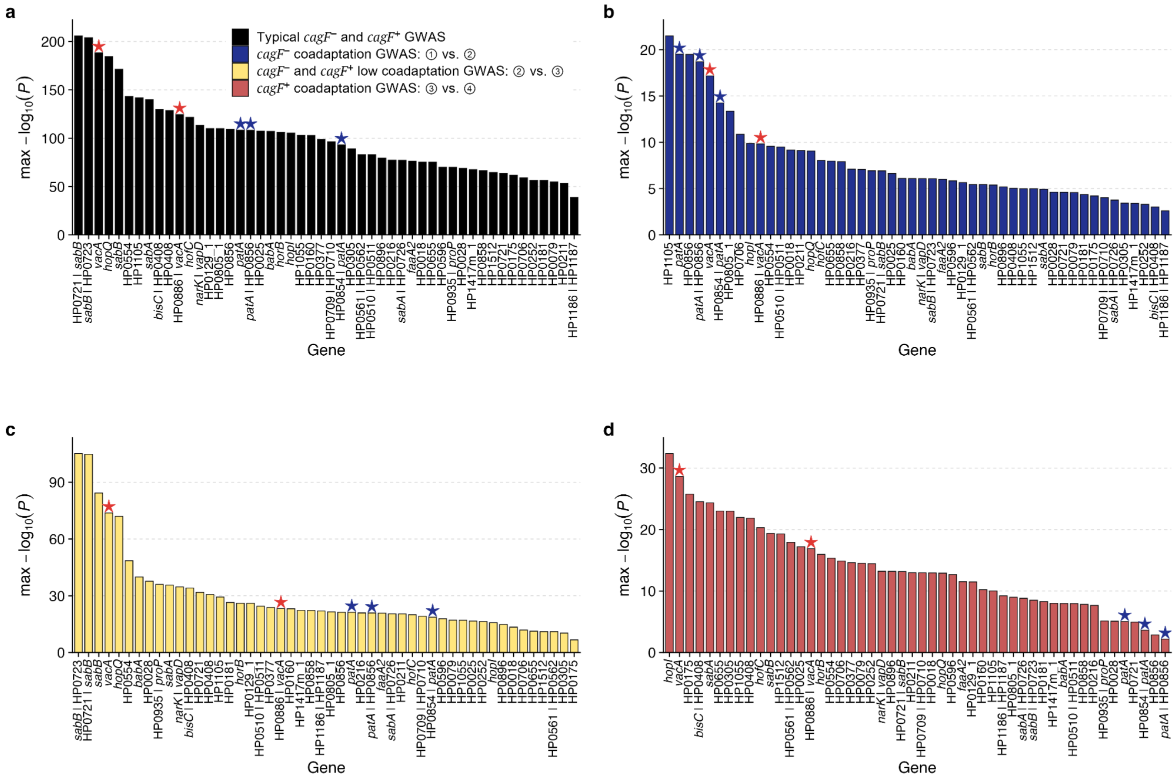


**Supplementary Fig. 9. Comparison-specific GWAS signals underlying candidate coadaptation loci.** Bar plots show the top gene-level association signals from four GWAS comparisons: typical hpEurope *cagF*$^-$ versus *cagF*$^+$ strains, *cagF*$^-$ low versus intermediate coadaptation groups (① vs. ②), *cagF*$^-$ intermediate versus *cagF*$^+$ low coadaptation groups (② vs. ③), and *cagF*$^+$ low versus high coadaptation groups (③ vs. ④). The labels ①–④ used here are defined in **Fig. 5a**.Within each panel, genes or intergenic regions are ordered by their maximum $-\log_{10}(P)$ value in that comparison. Gene names were assigned from the 26695 GFF annotation where available; HP locus tags were retained when no confident gene name was assigned. Red and blue stars mark *vacA*- and *patA*-associated signals, respectively. This comparison-specific view complements the cumulative signal plot by showing whether candidate loci are driven by a single GWAS comparison or recur across multiple stages of the cagPAI coadaptation trajectory.

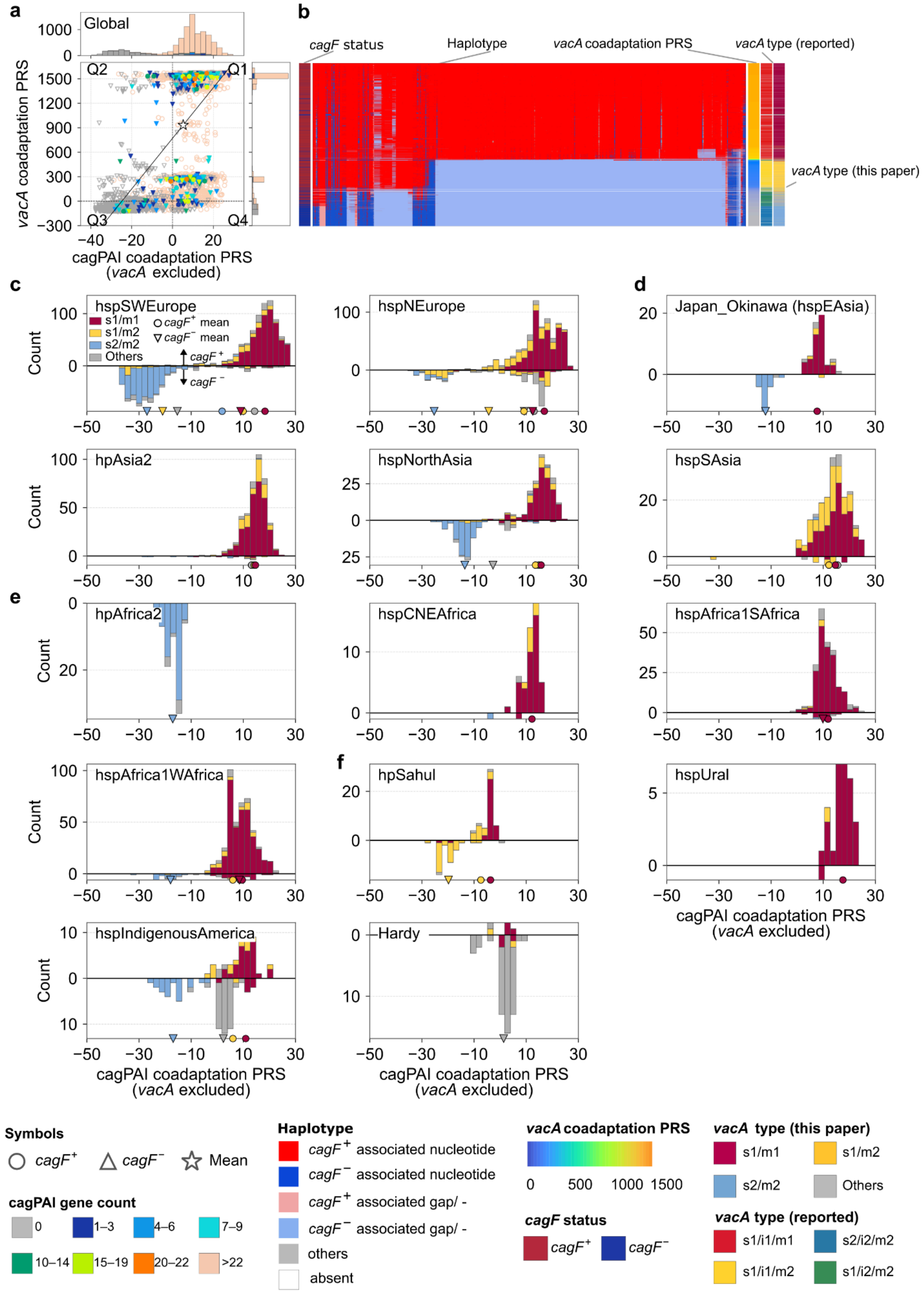


**Supplementary Fig. 10. Relationships among *vacA* variation, cagPAI integrity and coadaptation across global *H. pylori* populations. a,** Association between *vacA* and other gene coadaptation polygenic risk score (PRS) for global isolates, $cag^{-}$ isolates showed enrichment in the lower-left quadrant (*vacA* PRS < 0 and other gene PRS < 0), whereas $cag^{+}$ isolates were predominantly distributed in the upper-right quadrant (*vacA* PRS > 0 and other gene PRS > 0). Each strain was represented as a point in this space, with point shape indicating *cag* status (circles for *cagF*$^+$, triangles for *cagF*$^-$). Each point is coloured according to the number of cagPAI genes, grouped into eight categories. Point shapes defined by *cagF* status. Dashed lines: zero PRS on each axis, dividing the plots into four quadrants (Q1–Q4). Solid black line: linear regression fitted across all isolates. Star: overall mean PRS value. Marginal histograms show the distributions of PRS values along each axis, coloured by cagPAI gene count category. **b,** Heatmap of global *vacA* haplotype patterns. Strains are sorted from top to bottom by *vacA* coadaptation PRS in descending order, corresponding with the scatter plot (left). This top-to-bottom sorting reveals three clear evolutionary groups: Top: A *cag*$^+$ group with *cag*$^+$ associated allele (red blocks); Middle: An intermediate transition zone showing mixed patterns (red and blue mixed blocks); Bottom: A *cag*$^-$ adapted group with *cag*$^-$ associated allele (blue blocks). **c-f,** Mirrored stacked histograms showing the distributions of four *vacA* genotype categories—s1m1, s1m2, s2m2 and others—according to the cagPAI coadaptation PRS (exclude *vacA*), recalculated with variants in the *vacA* locus excluded, across the global dataset and individual populations, recalculated with the *vacA* locus excluded, across the global dataset and individual populations. Upward bars represent *cag*$^+$ strains (*cagF*$^+$), downward bars represent *cag*$^-$ strains (*cagF*$^-$), and bar heights indicate strain counts. Coloured circles and downward-pointing triangles on the x axis indicate the mean PRS for each *vacA* class among *cag*$^+$ and *cag*$^-$ strains, respectively; these markers are shown only for *vacA* class–cag status groups containing at least ten strains. Populations are grouped as **c,** European (hspSWEurope, hspNEurope), **d,** Asian (Okinawa_hspEasia, hpAsia2, hspNorthAsia, hspSAsia), **e,** African (hpAfrica2, hspCNEAfrica, hspAfrica1SAfrica and hspAfrica1WAfrica) and **f,** other lineages (hpSahul, hspUral, hspIndigenousAmerica and Hardy)**.**

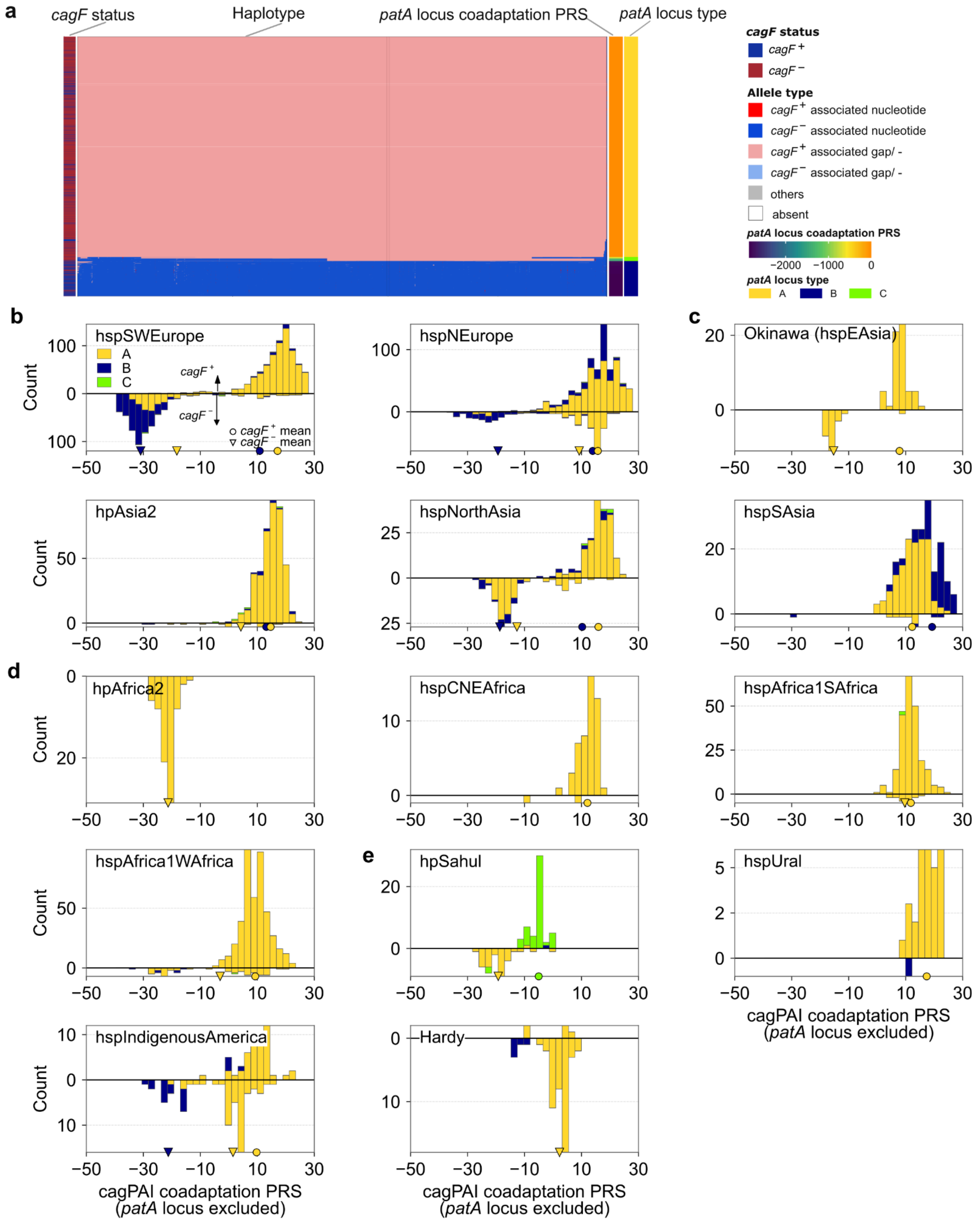

**Supplementary Fig. 11. *patA* locus haplotypes and cagPAI coadaptation across global *H. pylori* populations. a,** Haplotype structure across the HP0854-0858 (*patA*) locus. Each row represents a strain. Tracks show *cag* status based on *cagF* presence, alleles associated with *cag*$^{+}$ or *cag*$^{-}$ status and partial/truncated regions, the *patA*-locus coadaptation PRS, and the three major *patA* locus haplotypes—type A, B and C. **b-e,** Mirrored stacked histograms showing the distribution of *patA*-locus haplotypes according to the cagPAI coadaptation PRS, recalculated with variants in the patA-locus excluded. Upward and downward bars represent *cag*$^{+}$ (*cagF*$^{+}$) and *cag*$^{-}$ (*cagF*$^{-}$) strains, respectively; bar heights indicate strain counts and colours denote *patA* haplotypes. Filled upward-pointing, and open downward-pointing triangles indicate the mean PRS of strains with complete, and absent cagPAIs, respectively. Results are shown for the individual populations.
