## Extended_Data_Fig_1 for "History of coadaptation to the *cag* pathogenicity island in *Helicobacter pylori*"

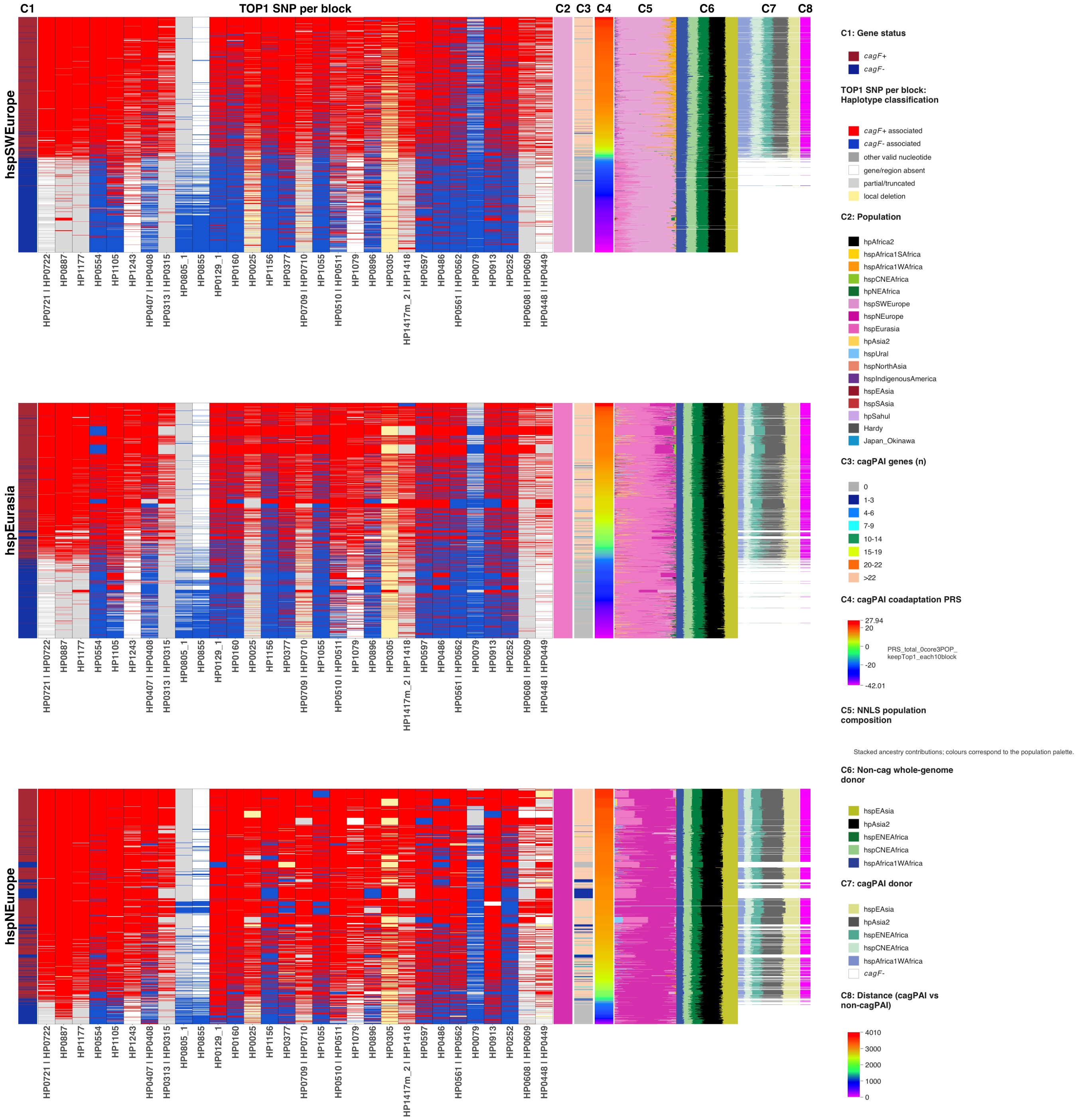
C1
TOP1 SNP per block
C2
C3
C4
C5
C6
C7
C8
hspSWEurope
hspEurasia
hspNEurope
HP0721 | HP0722
HP0887
HP1177
HP0554
HP1105
HP1243
HP0407 | HP0408
HP0313 | HP0315
HP0805_1
HP0855
HP0129_1
HP0160
HP0025
HP1156
HP0377
HP0709 | HP0710
HP1055
HP0510 | HP0511
HP1079
HP0896
HP0305
HP1417m_2 | HP1418
HP0597
HP0486
HP0561 | HP0562
HP0079
HP0913
HP0252
HP0608 | HP0609
HP0448 | HP0449
C1: Gene status
cagF+
cagF-
TOP1 SNP per block: Haplotype classification
cagF+ associated
cagF- associated
other valid nucleotide
gene/region absent
partial/truncated
local deletion
C2: Population
hpAfrica2
hspAfrica1SAfrica
hspAfrica1WAfrica
hspCNEAfrica
hpNEAfrica
hspSWEurope
hspNEurope
hspEurasia
hpAsia2
hspUral
hspNorthAsia
hspIndigenousAmerica
hspEAsia
hspSAsia
hpSahul
Hardy
Japan_Okinawa
C3: cagPAI genes (n)
0
1-3
4-6
7-9
10-14
15-19
20-22
>22
C4: cagPAI coadaptation PRS
27.94
20
0
-20
-42.01
PRS_total_0core3POP_keepTop1_each10block
C5: NNLS population composition
Stacked ancestry contributions; colours correspond to the population palette.
C6: Non-cag whole-genome donor
hspEAsia
hpAsia2
hspENEAfrica
hspCNEAfrica
hspAfrica1WAfrica
C7: cagPAI donor
hspEAsia
hpAsia2
hspENEAfrica
hspCNEAfrica
hspAfrica1WAfrica
cagF-
C8: Distance (cagPAI vs non-cagPAI)
4010
3000
2000
1000
0

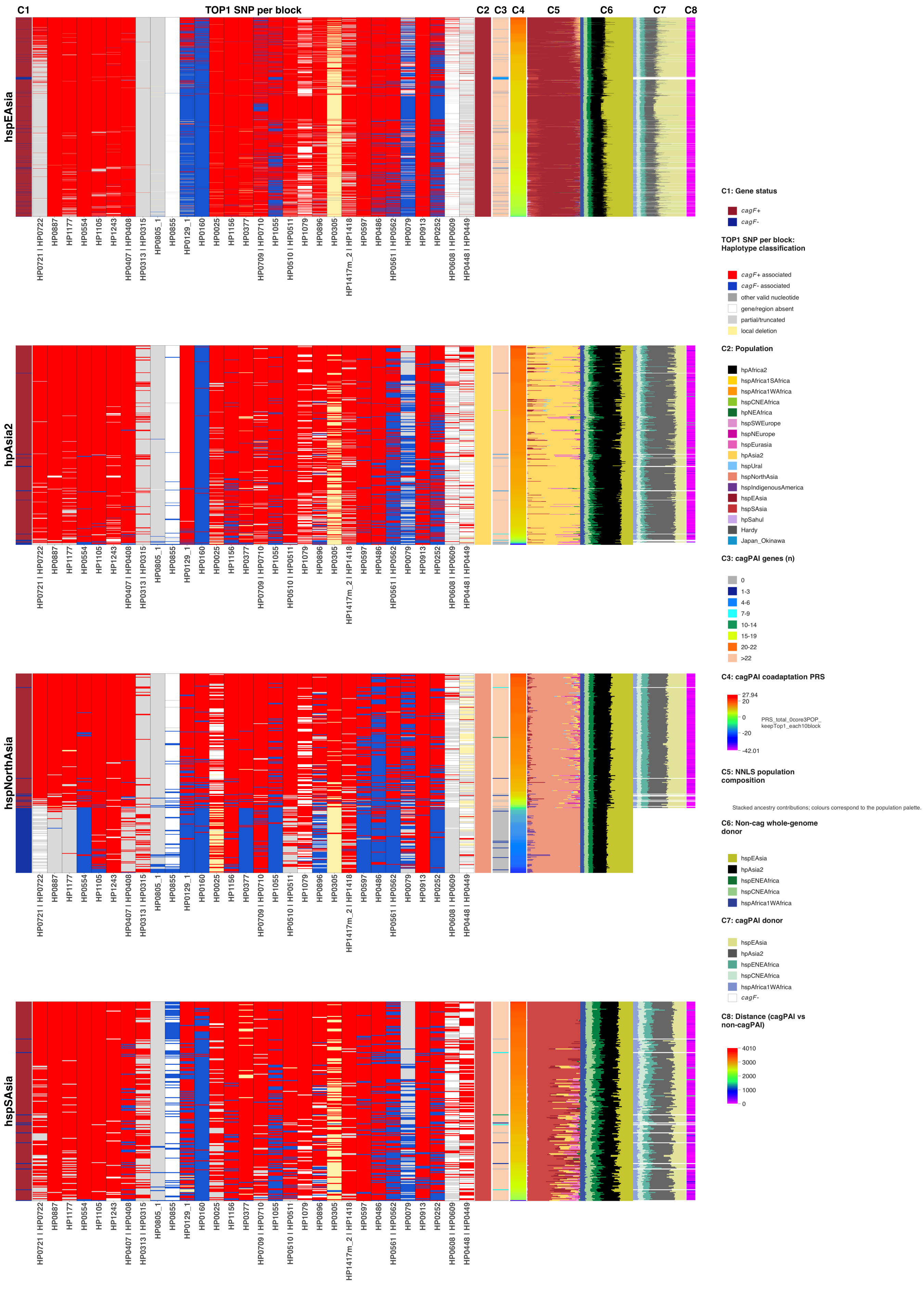

C1
TOP1 SNP per block
C2
C3
C4
C5
C6
C7
C8
hspEAsia
hpAsia2
hspNorthAsia
hspSAsia
HP0721 | HP0722
HP0887
HP1177
HP0554
HP1105
HP1243
HP0407 | HP0408
HP0313 | HP0315
HP0805_1
HP0855
HP0129_1
HP0160
HP0025
HP1156
HP0377
HP0709 | HP0710
HP1055
HP0510 | HP0511
HP1079
HP0896
HP0305
HP1417m_2 | HP1418
HP0597
HP0486
HP0561 | HP0562
HP0079
HP0913
HP0252
HP0608 | HP0609
HP0448 | HP0449
C1: Gene status
cagF+
cagF-
TOP1 SNP per block: Haplotype classification
cagF+ associated
cagF- associated
other valid nucleotide
gene/region absent
partial/truncated
local deletion
C2: Population
hpAfrica2
hspAfrica1SAfrica
hspAfrica1WAfrica
hspCNEAfrica
hpNEAfrica
hspSWEurope
hspNEurope
hspEurasia
hpAsia2
hspUral
hspNorthAsia
hspIndigenousAmerica
hspEAsia
hspSAsia
hpSahul
Hardy
Japan_Okinawa
C3: cagPAI genes (n)
0
1-3
4-6
7-9
10-14
15-19
20-22
>22
C4: cagPAI coadaptation PRS
27.94
20
0
-20
-42.01
PRS_total_0core3POP_keepTop1_each10block
C5: NNLS population composition
Stacked ancestry contributions; colours correspond to the population palette.
C6: Non-cag whole-genome donor
hspEAsia
hpAsia2
hspENEAfrica
hspCNEAfrica
hspAfrica1WAfrica
C7: cagPAI donor
hspEAsia
hpAsia2
hspENEAfrica
hspCNEAfrica
hspAfrica1WAfrica
cagF-
C8: Distance (cagPAI vs non-cagPAI)
4010
3000
2000
1000
0

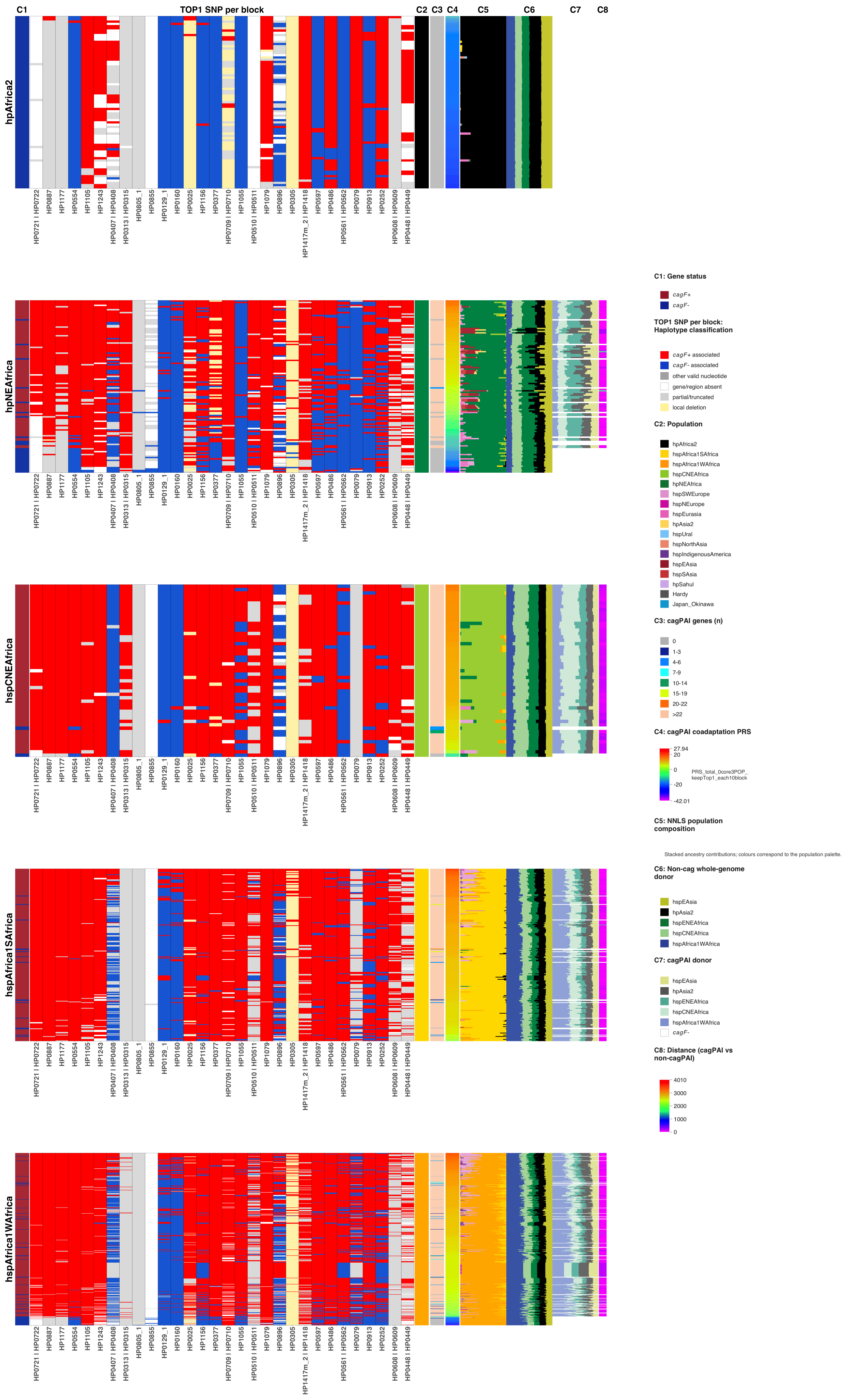
C1
TOP1 SNP per block
C2
C3
C4
C5
C6
C7
C8
hpAfrica2
hpNEAfrica
hspCNEAfrica
hspAfrica1SAfrica
hspAfrica1WAfrica
HP0721 | HP0722
HP0887
HP1177
HP0554
HP1105
HP1243
HP0407 | HP0408
HP0313 | HP0315
HP0805_1
HP0855
HP0129_1
HP0160
HP0025
HP1156
HP0377
HP0709 | HP0710
HP1055
HP0510 | HP0511
HP1079
HP0896
HP0305
HP1417m_2 | HP1418
HP0597
HP0486
HP0561 | HP0562
HP0079
HP0913
HP0252
HP0608 | HP0609
HP0448 | HP0449
C1: Gene status
cagF+
cagF-
TOP1 SNP per block: Haplotype classification
cagF+ associated
cagF- associated
other valid nucleotide
gene/region absent
partial/truncated
local deletion
C2: Population
hpAfrica2
hspAfrica1SAfrica
hspAfrica1WAfrica
hspCNEAfrica
hpNEAfrica
hspSWEurope
hspNEurope
hspEurasia
hpAsia2
hspUral
hspNorthAsia
hspIndigenousAmerica
hspEAsia
hspSAsia
hpSahul
Hardy
Japan_Okinawa
C3: cagPAI genes (n)
0
1-3
4-6
7-9
10-14
15-19
20-22
>22
C4: cagPAI coadaptation PRS
27.94
20
0
-20
-42.01
PRS_total_0core3POP_keepTop1_each10block
C5: NNLS population composition
Stacked ancestry contributions; colours correspond to the population palette.
C6: Non-cag whole-genome donor
hspEAsia
hpAsia2
hspENEAfrica
hspCNEAfrica
hspAfrica1WAfrica
C7: cagPAI donor
hspEAsia
hpAsia2
hspENEAfrica
hspCNEAfrica
hspAfrica1WAfrica
cagF-
C8: Distance (cagPAI vs non-cagPAI)
4010
3000
2000
1000
0

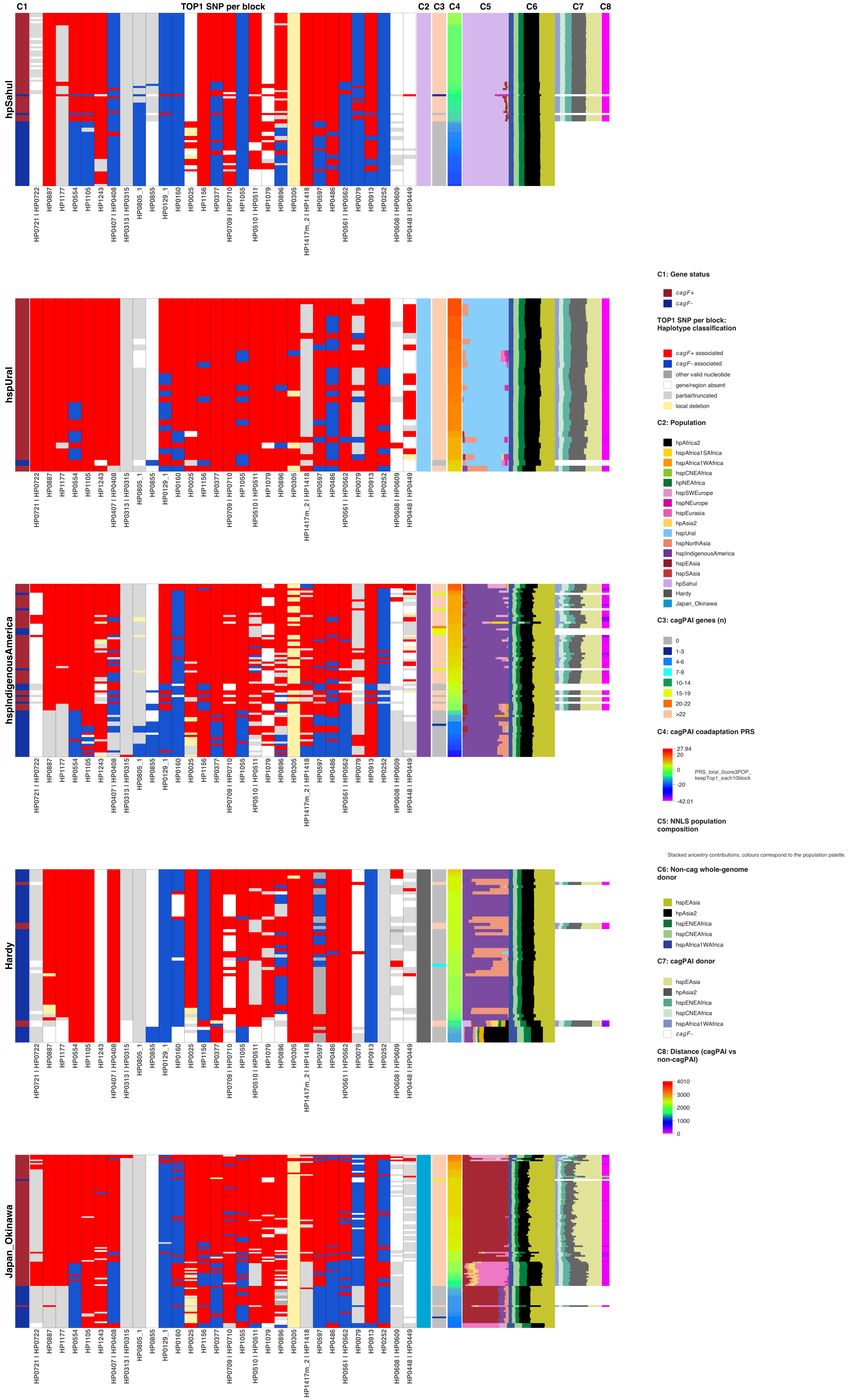
C1
TOP1 SNP per block
C2
C3
C4
C5
C6
C7
C8
hpSahul
hspUral
hspIndigenousAmerica
Hardy
Japan_Okinawa
HP0721 | HP0722
HP0887
HP1177
HP0554
HP1105
HP1243
HP0407 | HP0408
HP0313 | HP0315
HP0805_1
HP0855
HP0129_1
HP0160
HP0025
HP1156
HP0377
HP0709 | HP0710
HP1055
HP0510 | HP0511
HP1079
HP0896
HP0305
HP1417m_2 | HP1418
HP0597
HP0486
HP0561 | HP0562
HP0079
HP0913
HP0252
HP0608 | HP0609
HP0448 | HP0449
C1: Gene status
cagF+
cagF-
TOP1 SNP per block: Haplotype classification
cagF+ associated
cagF- associated
other valid nucleotide
gene/region absent
partial/truncated
local deletion
C2: Population
hpAfrica2
hspAfrica1SAfrica
hspAfrica1WAfrica
hspCNEAfrica
hpNEAfrica
hspSWEurope
hspNEurope
hspEurasia
hpAsia2
hspUral
hspNorthAsia
hspIndigenousAmerica
hspEAsia
hspSAsia
hpSahul
Hardy
Japan_Okinawa
C3: cagPAI genes (n)
0
1-3
4-6
7-9
10-14
15-19
20-22
>22
C4: cagPAI coadaptation PRS
27.94
20
0
-20
-42.01
PRS_total_0core3POP_
keepTop1_each10block
C5: NNLS population composition
Stacked ancestry contributions; colours correspond to the population palette.
C6: Non-cag whole-genome donor
hspEAsia
hpAsia2
hspENEAfrica
hspCNEAfrica
hspAfrica1WAfrica
C7: cagPAI donor
hspEAsia
hpAsia2
hspENEAfrica
hspCNEAfrica
hspAfrica1WAfrica
cagF-
C8: Distance (cagPAI vs non-cagPAI)
4010
3000
2000
1000
0

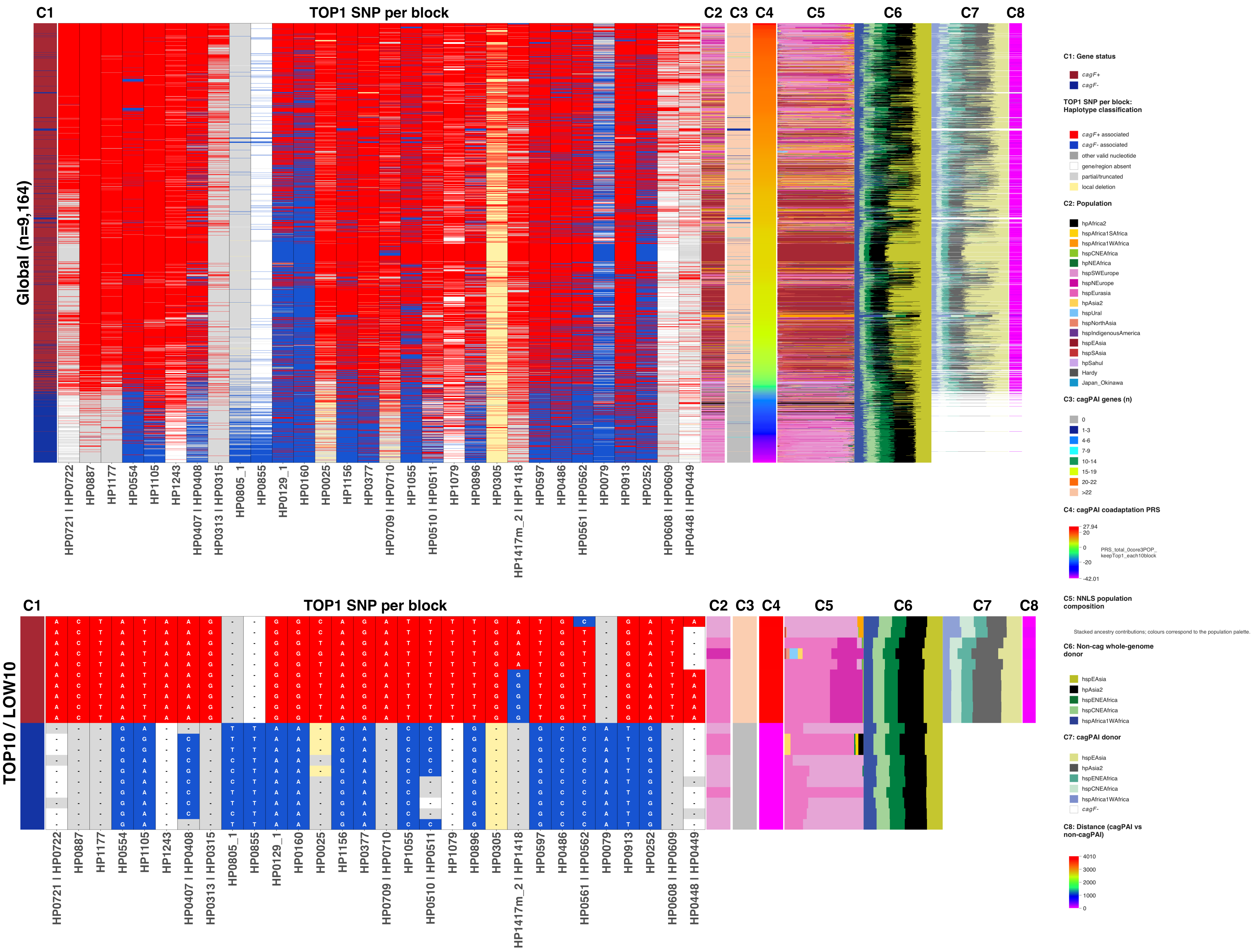
C1
TOP1 SNP per block
C2
C3
C4
C5
C6
C7
C8
Global (n=9,164)
TOP10 / LOW10
HP0721 I HP0722
HP0887
HP1177
HP0554
HP1105
HP1243
HP0407 I HP0408
HP0313 I HP0315
HP0805_1
HP0855
HP0129_1
HP0160
HP0025
HP1156
HP0377
HP0709 I HP0710
HP1055
HP0510 I HP0511
HP1079
HP0896
HP0305
HP1417m_2 I HP1418
HP0597
HP0486
HP0561 I HP0562
HP0079
HP0913
HP0252
HP0608 I HP0609
HP0448 I HP0449
C1: Gene status
cagF+
cagF-
TOP1 SNP per block: Haplotype classification
cagF+ associated
cagF- associated
other valid nucleotide
gene/region absent
partial/truncated
local deletion
C2: Population
hpAfrica2
hspAfrica1SAfrica
hspAfrica1WAfrica
hspCNEAfrica
hpNEAfrica
hspSWEurope
hspNEurope
hspEurasia
hpAsia2
hspUral
hspNorthAsia
hspIndigenousAmerica
hspEAsia
hspSAsia
hpSahul
Hardy
Japan_Okinawa
C3: cagPAI genes (n)
0
1-3
4-6
7-9
10-14
15-19
20-22
>22
C4: cagPAI coadaptation PRS
27.94
20
0
-20
-42.01
PRS_total_0core3POP_keepTop1_each10block
C5: NNLS population composition
Stacked ancestry contributions; colours correspond to the population palette.
C6: Non-cag whole-genome donor
hspEAsia
hpAsia2
hspENEAfrica
hspCNEAfrica
hspAfrica1WAfrica
C7: cagPAI donor
hspEAsia
hpAsia2
hspENEAfrica
hspCNEAfrica
hspAfrica1WAfrica
cagF-
C8: Distance (cagPAI vs non-cagPAI)
4010
3000
2000
1000
0
