## Supplementary figures and images for "History of coadaptation to the *cag* pathogenicity island in *Helicobacter pylori*"

### Extended_Data_Fig_2

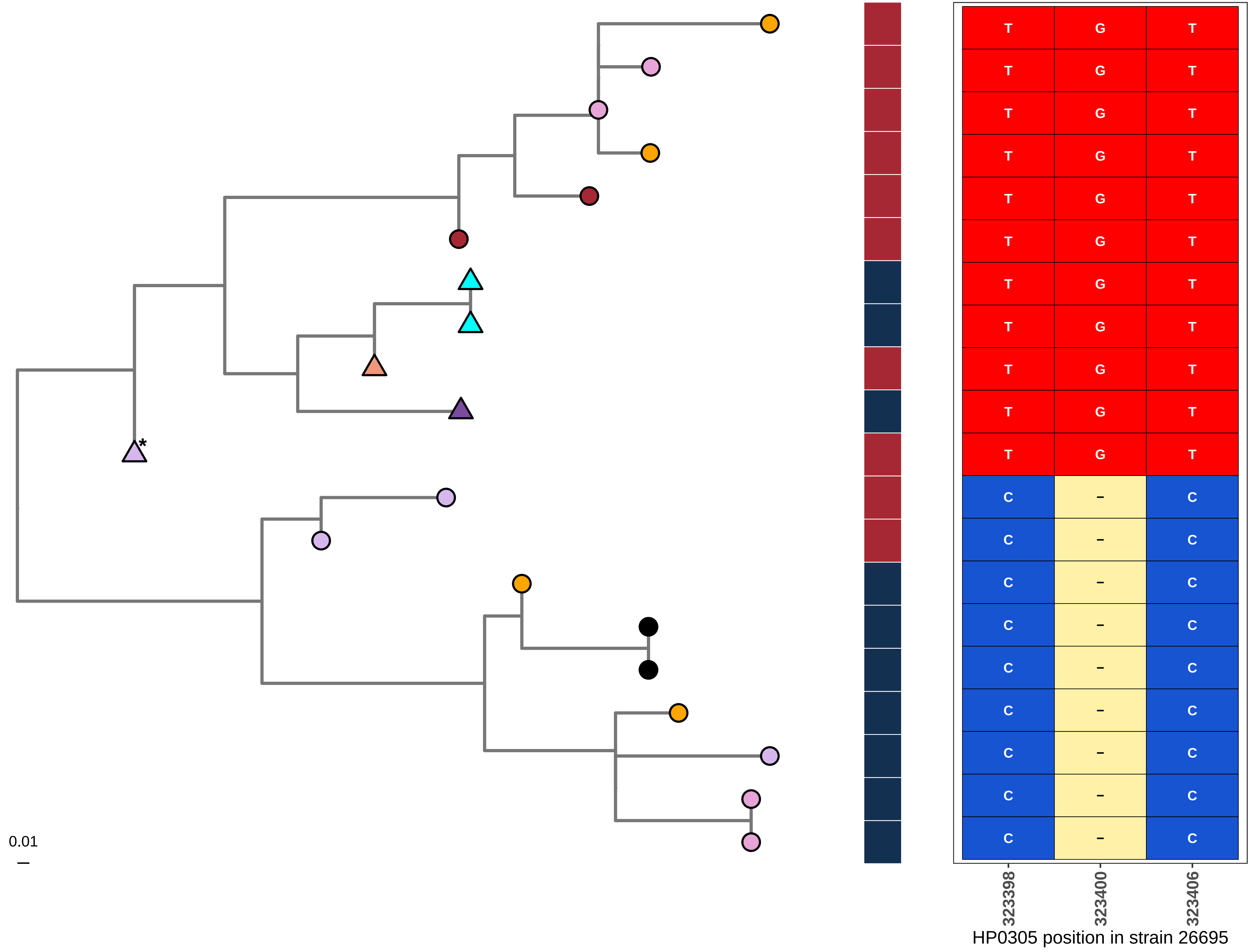
0.01
*
T G T
C – C
323398
323400
323406
HP0305 position in strain 26695

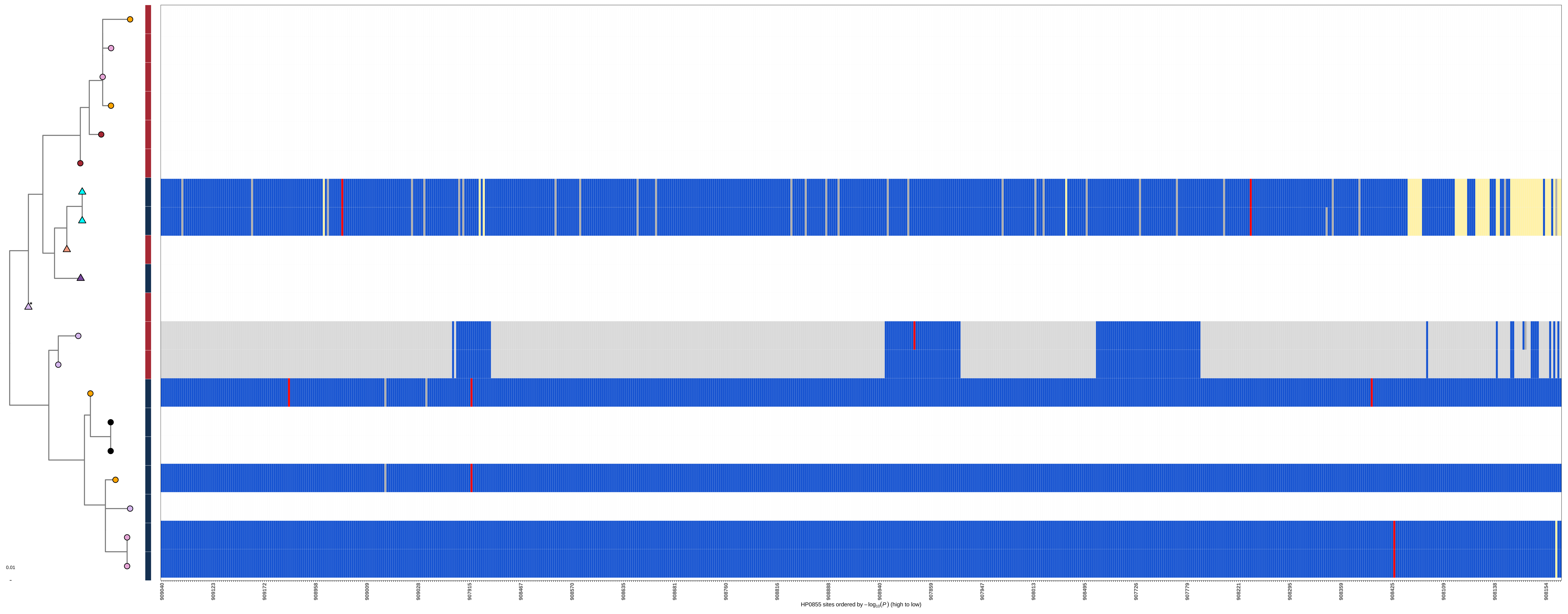
0.01
909040
909123
909172
908958
909009
909028
907915
908467
908570
908635
908681
908760
908816
908888
908940
907859
907947
908013
908495
907726
907779
908221
908295
908359
908425
908109
908138
908154
HP0855 sites ordered by $-\log_{10}(P)$ (high to low)
