## Extended_Data_Fig_3 for "History of coadaptation to the *cag* pathogenicity island in *Helicobacter pylori*"

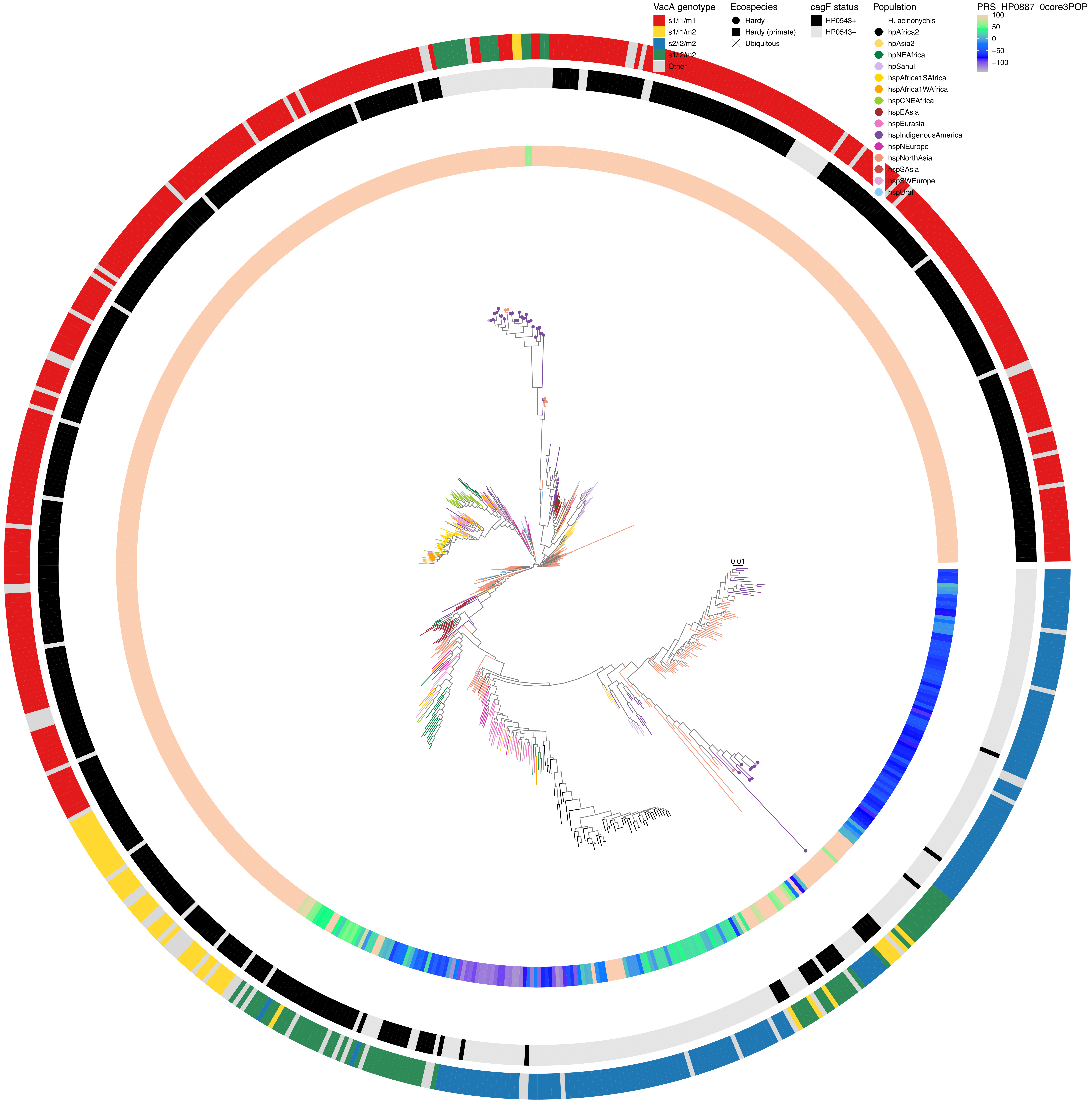

VacA genotype
s1/i1/m1
s1/i1/m2
s2/i2/m2
s1/i2/m2
Other
Ecospecies
Hardy
Hardy (primate)
Ubiquitous
cagF status
HP0543+
HP0543−
Population
H. acinonychis
hpAfrica2
hpAsia2
hpNEAfrica
hpSahul
hspAfrica1SAfrica
hspAfrica1WAfrica
hspCNEAfrica
hspEAsia
hspEurasia
hspIndigenousAmerica
hspNEurope
hspNorthAsia
hspSAsia
hspSWEurope
hspUral
PRS_HP0887_0core3POP
100
50
0
−50
−100
0.01

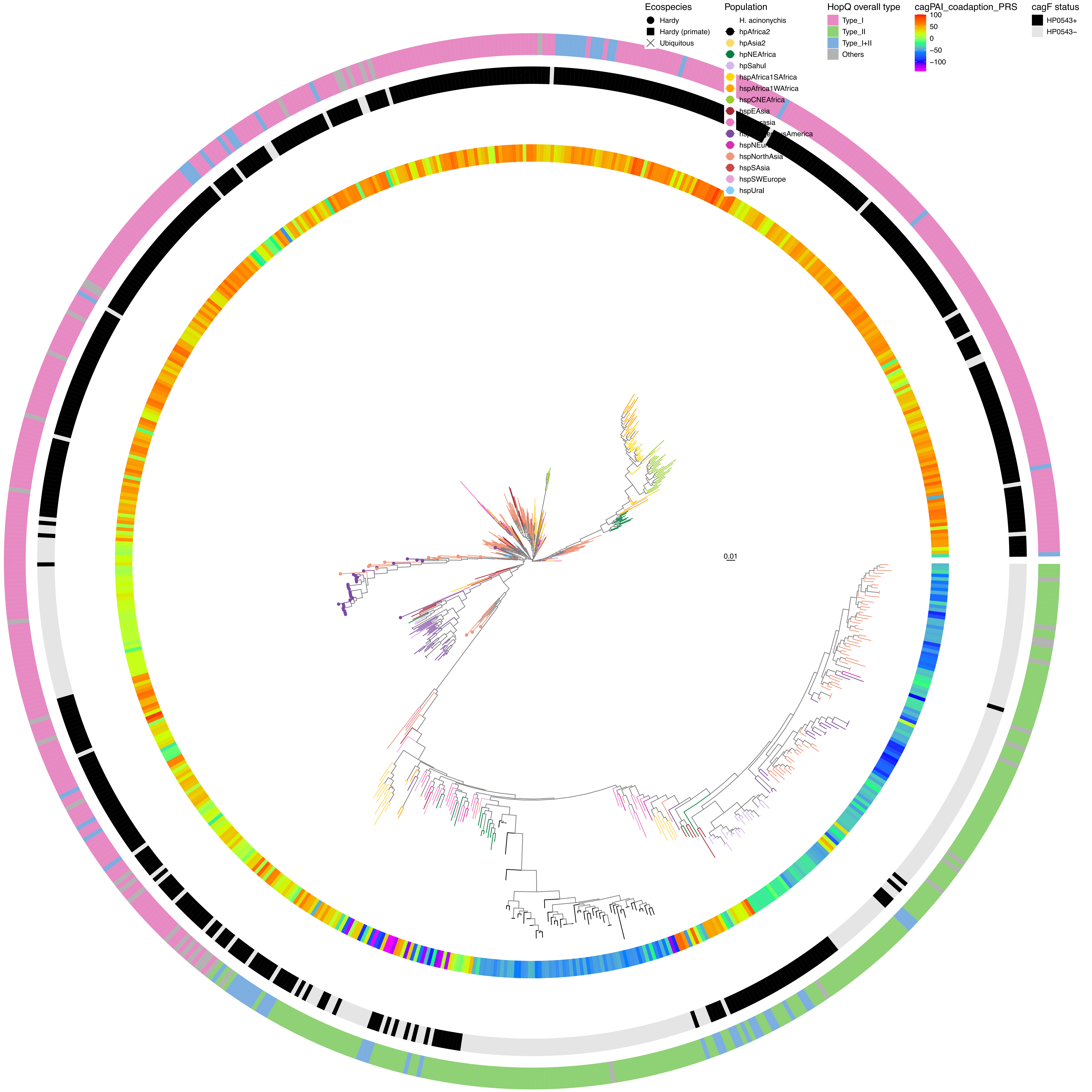
Ecospecies
Hardy
Hardy (primate)
Ubiquitous
Population
H. acinonychis
hpAfrica2
hpAsia2
hpNEAfrica
hpSahul
hspAfrica1SAfrica
hspAfrica1WAfrica
hspCNEAfrica
hspEAsia
hspNorthAsia
hspSAsia
hspSWEurope
hspUral
HopQ overall type
Type_I
Type_II
Type_I+II
Others
cagPAI_coadaption_PRS
100
50
0
−50
−100
cagF status
HP0543+
HP0543−
0.01

**HP0887**
**Value: 199**

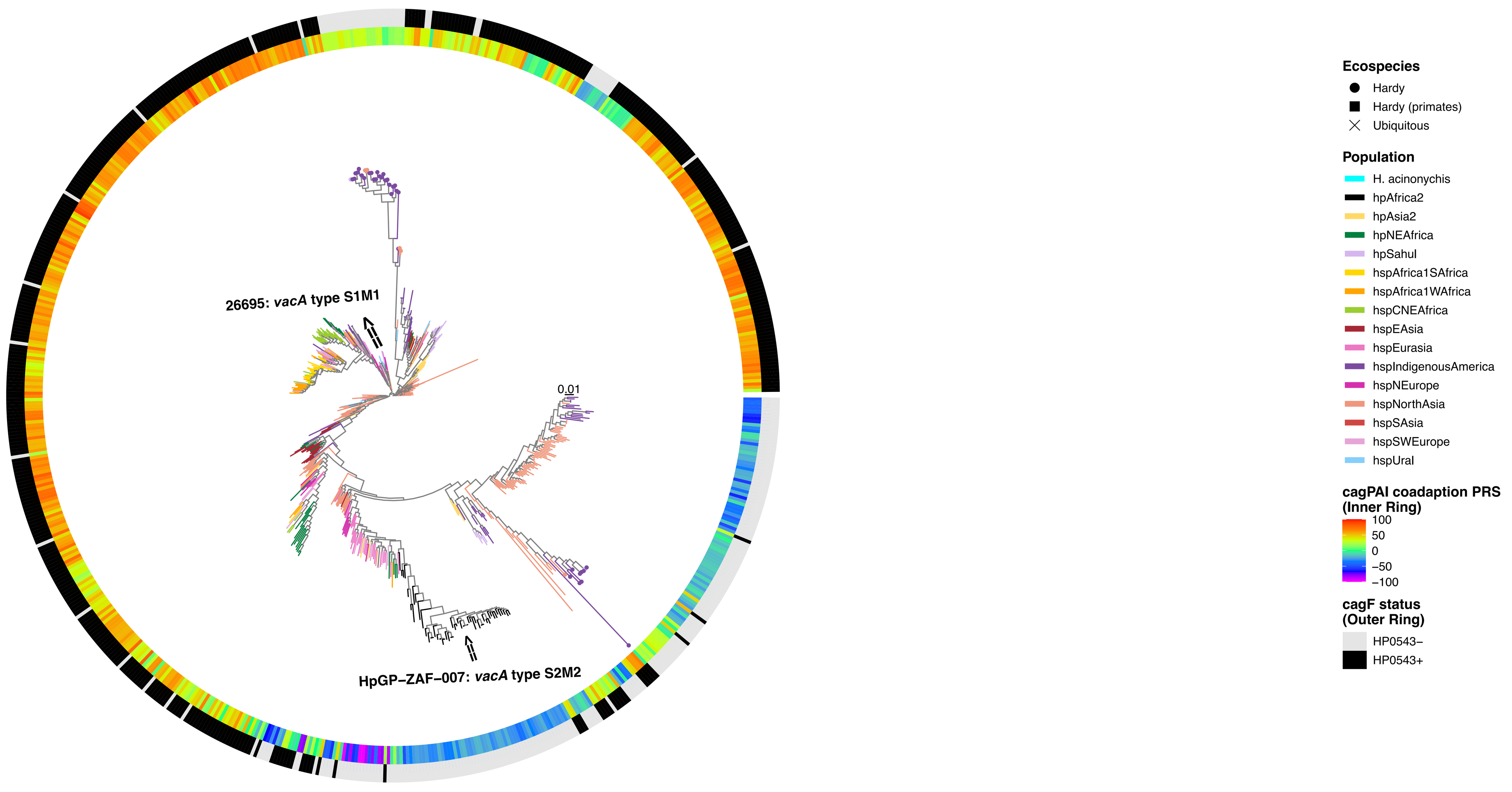


**HP1177**
**Value: 182**

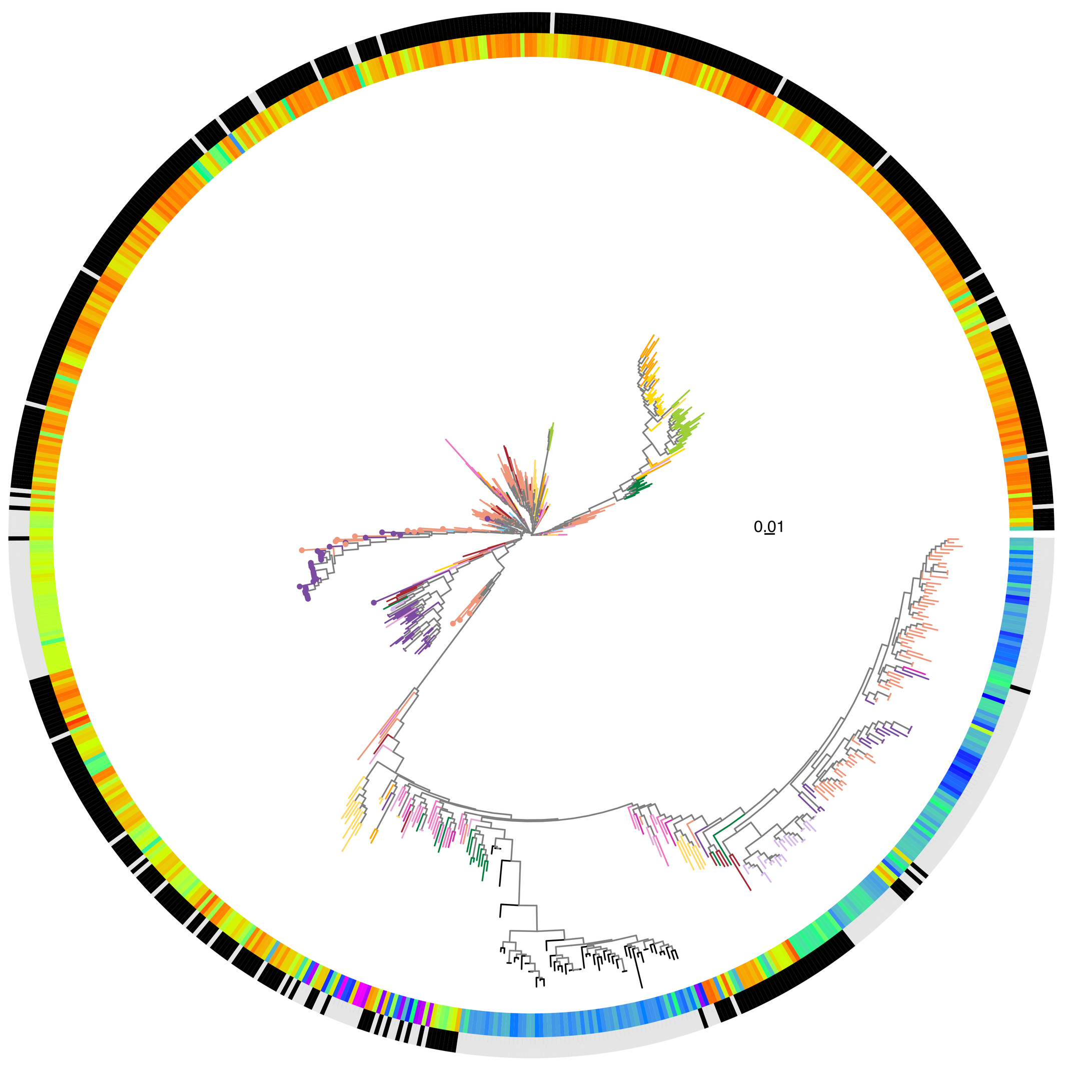


**HP0722**
**Value: 177**

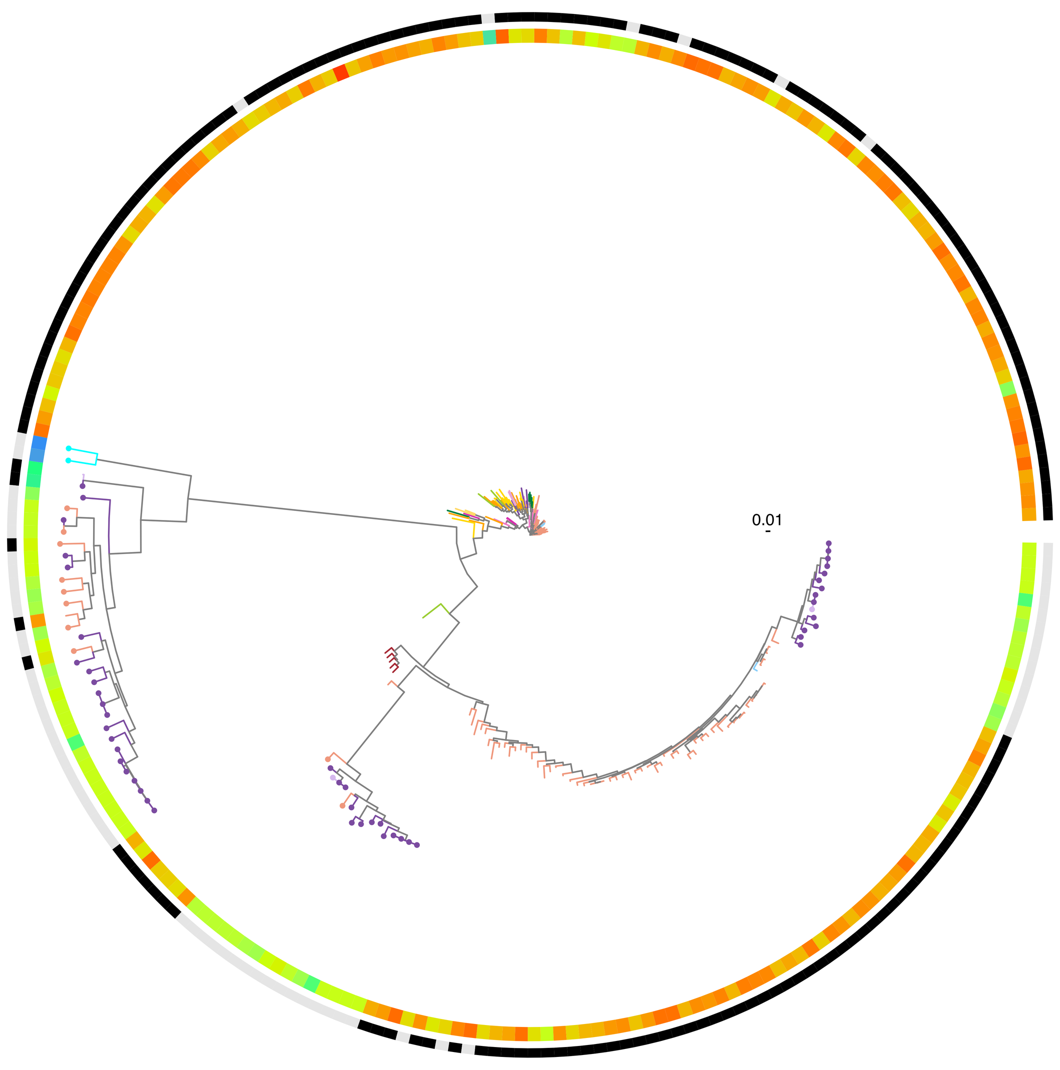


**HP0554**
**Value: 140**

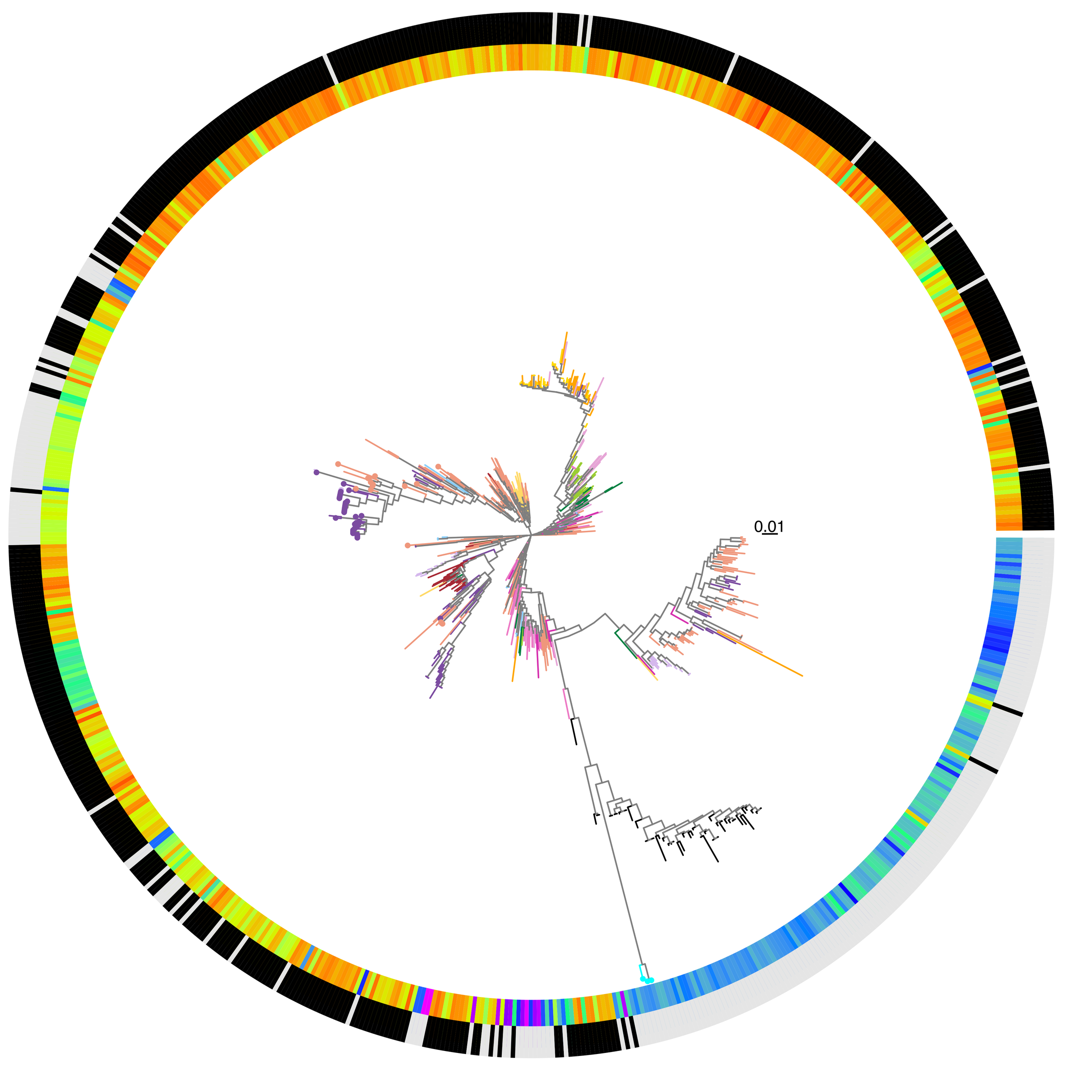


**HP1105**
**Value: 123**

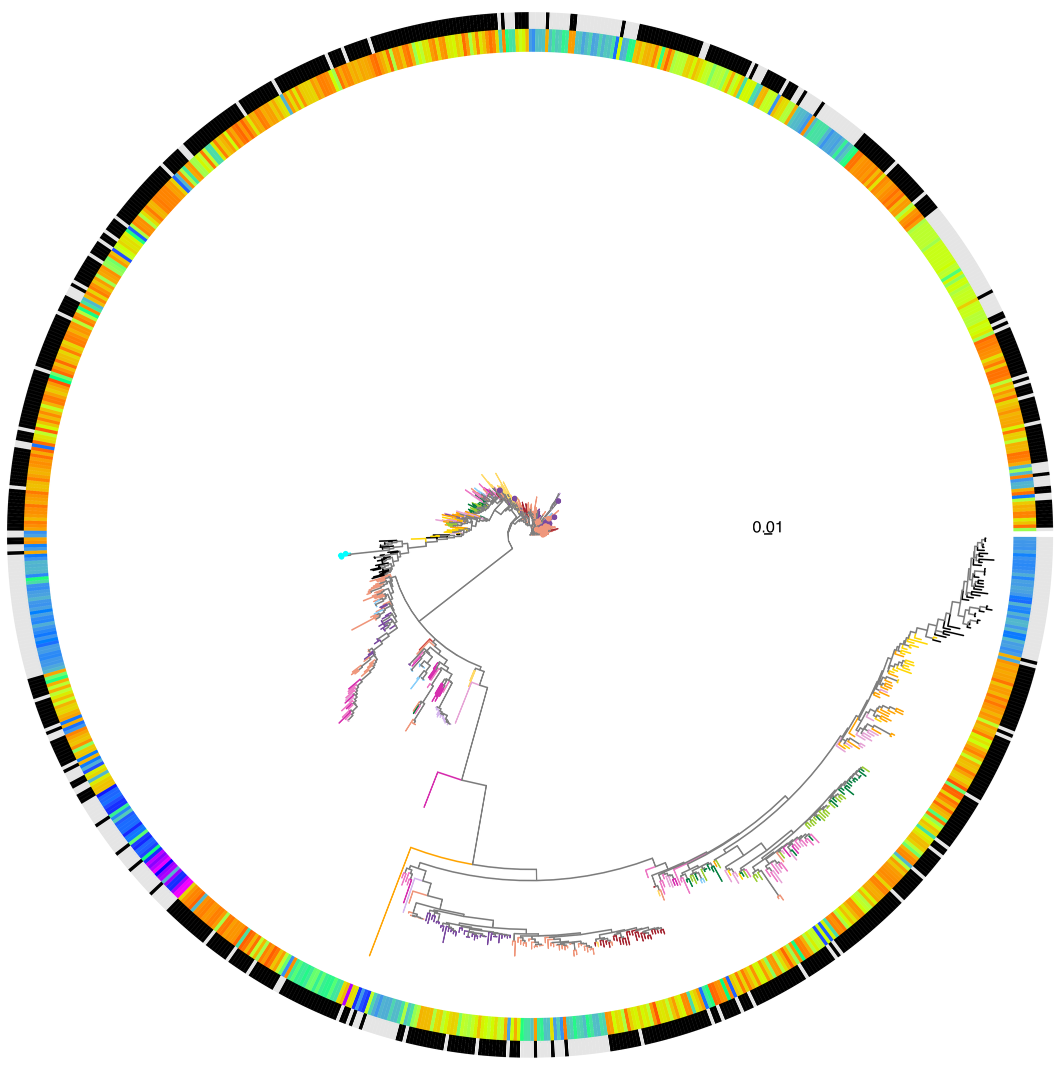


**HP0725**
**Value: 114**

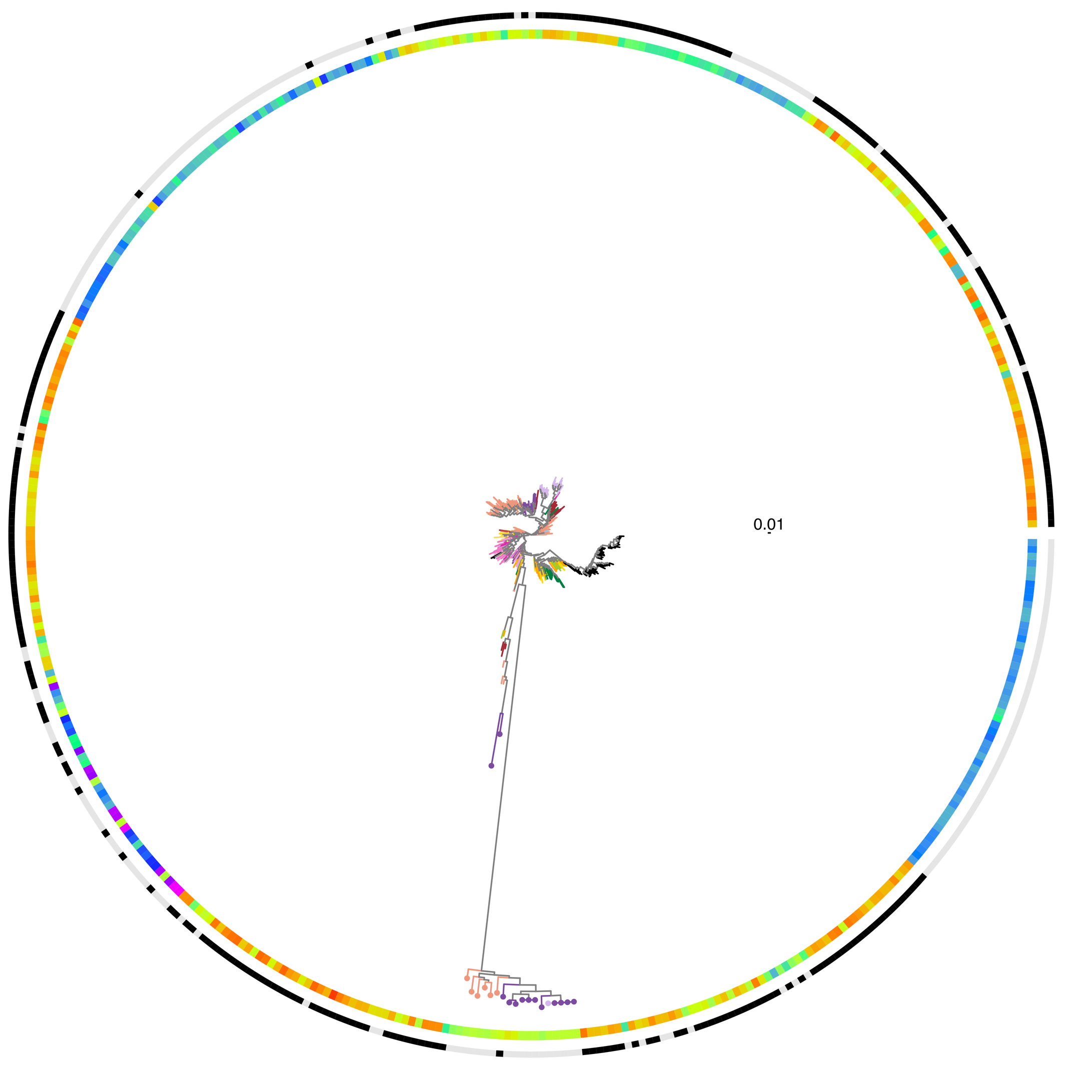


**HP1243**
**Value: 110**

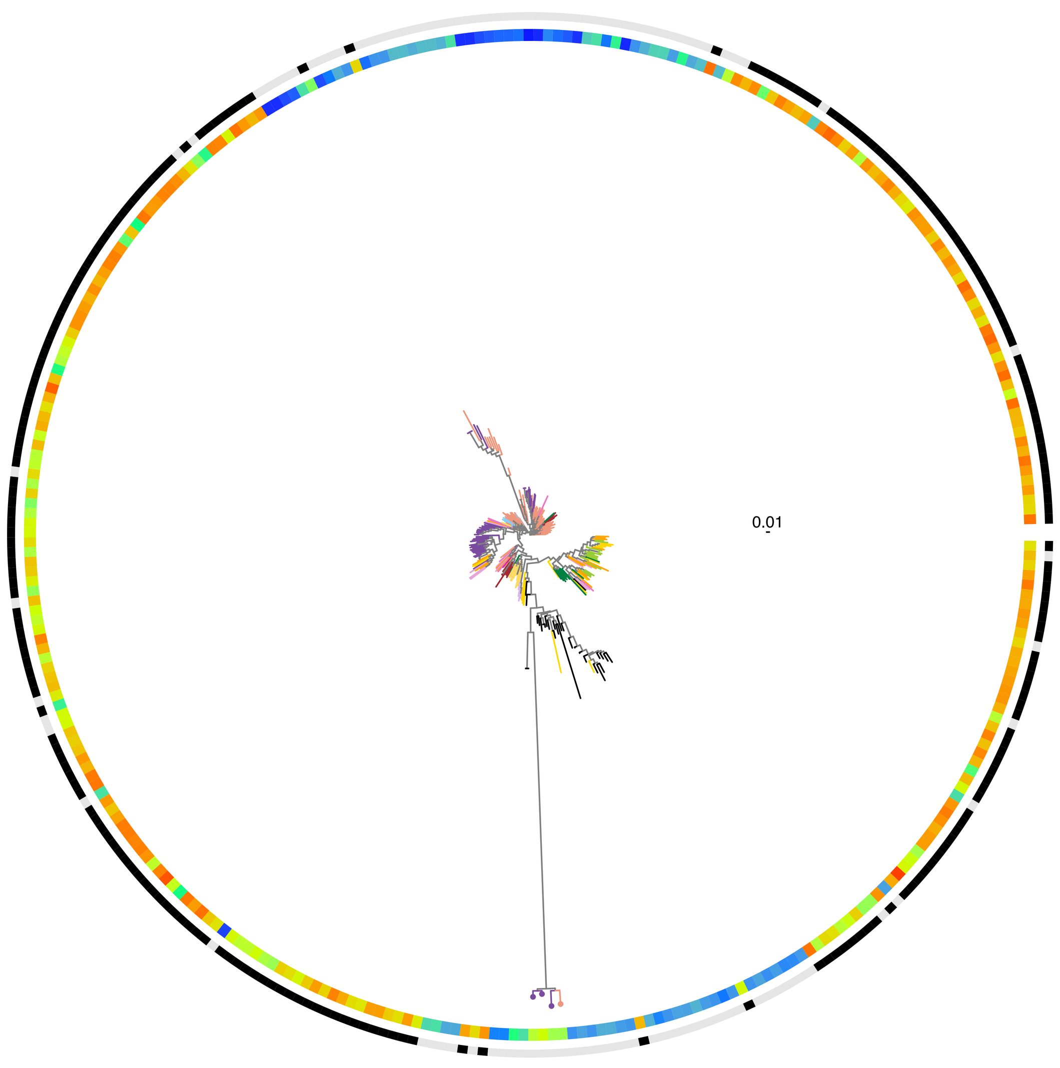

**HP0805_1**
**Value: 96**

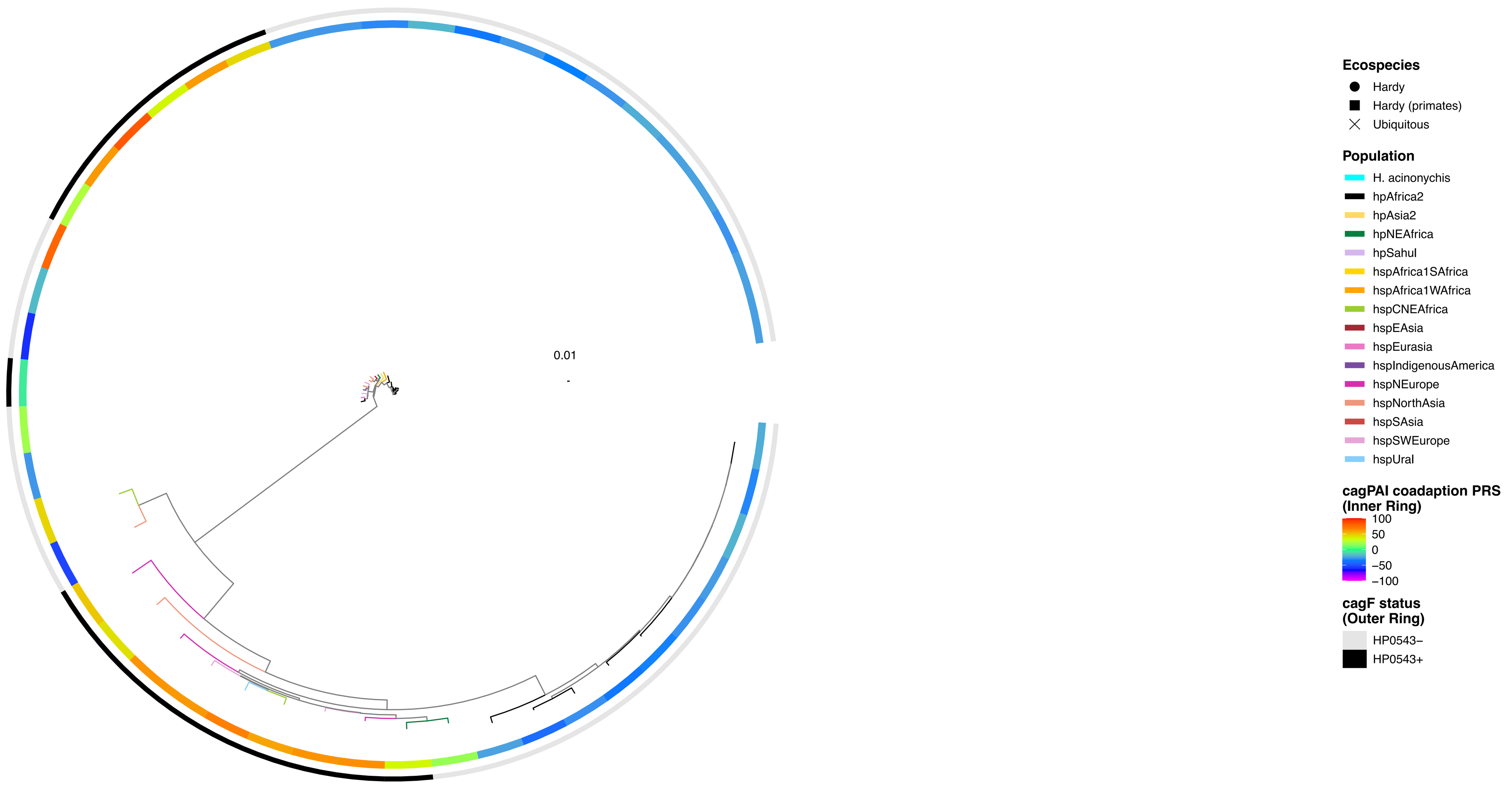


**HP0855**
**Value: 96**

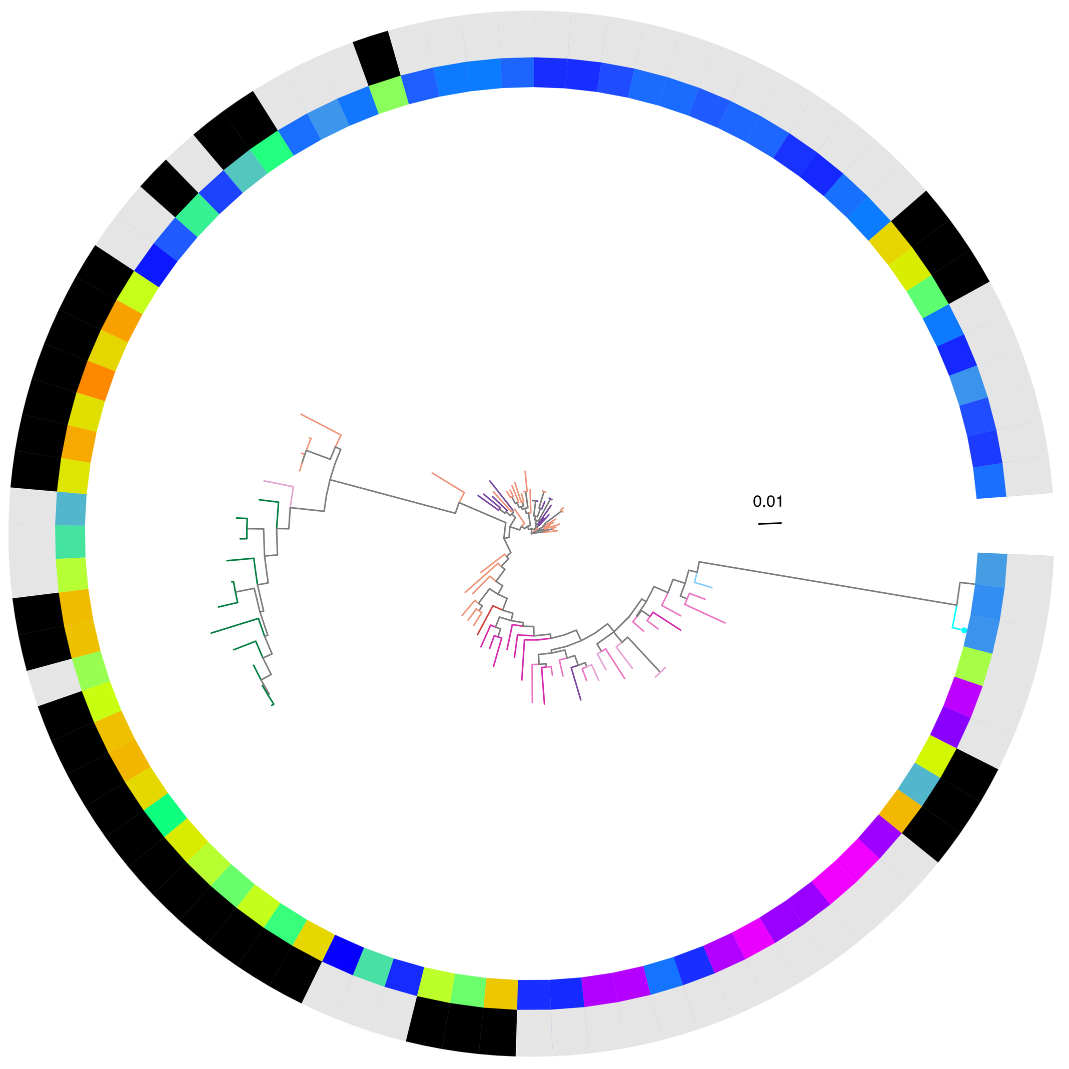


**HP0856**
**Value: 96**

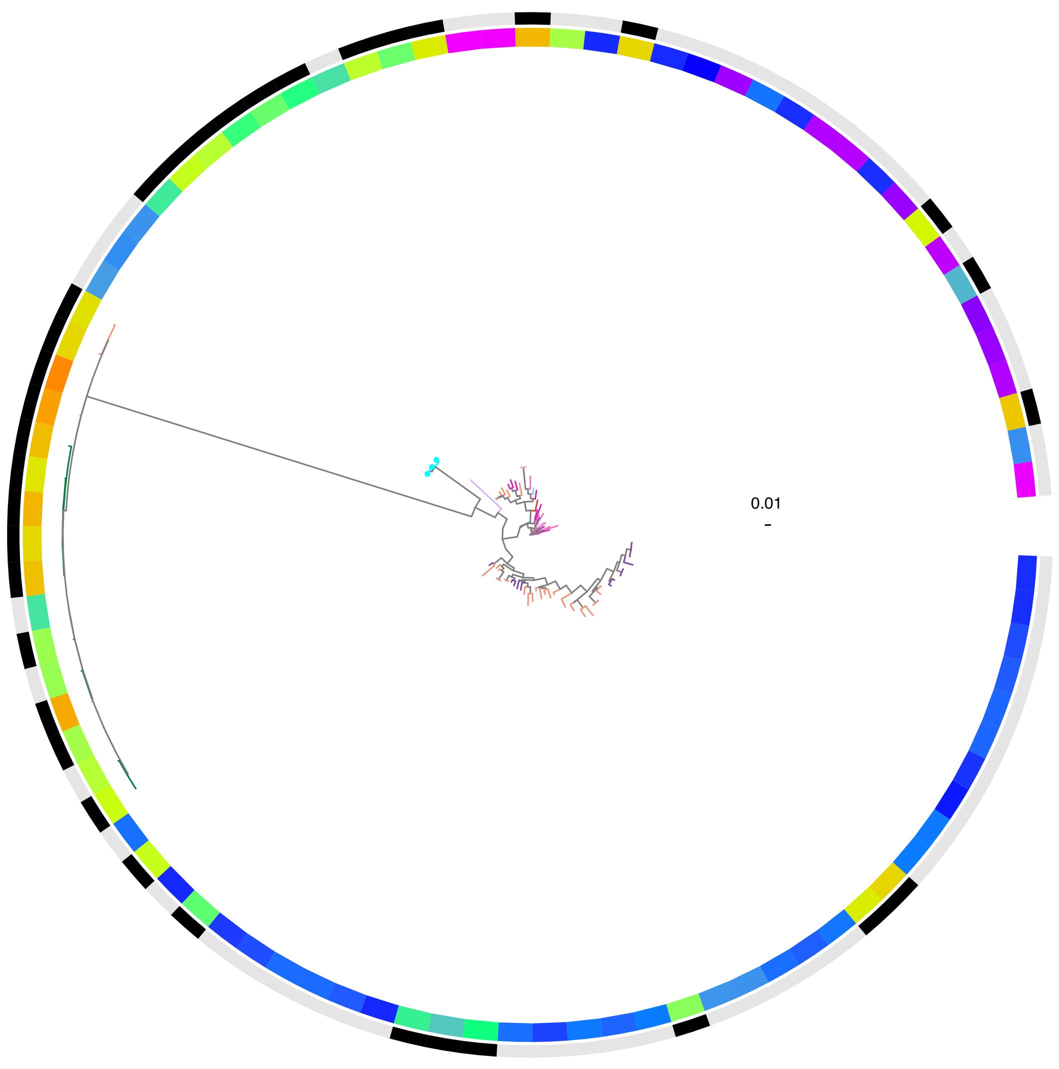


**HP0129_1**
**Value: 92**

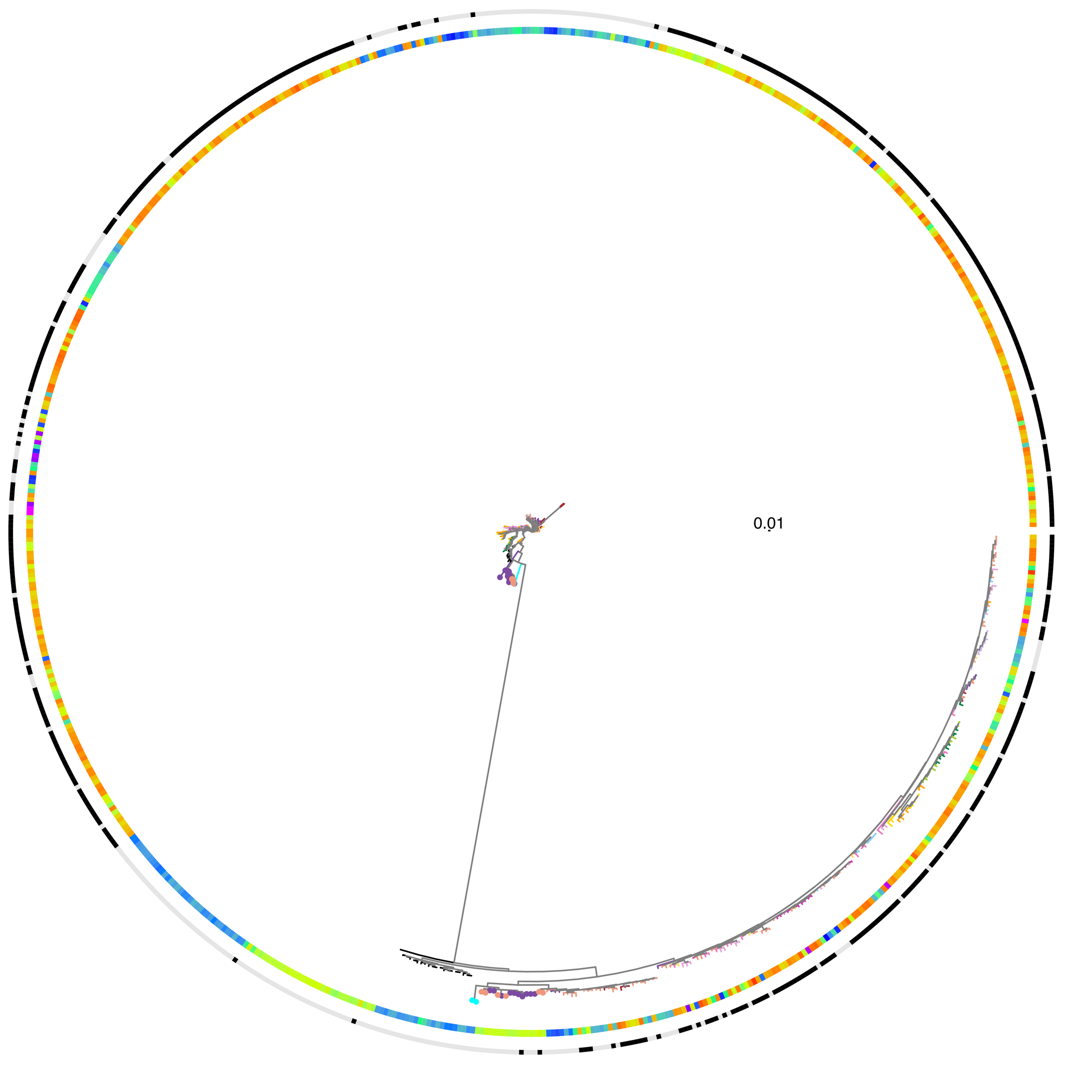


**HP0127**
**Value: 89**

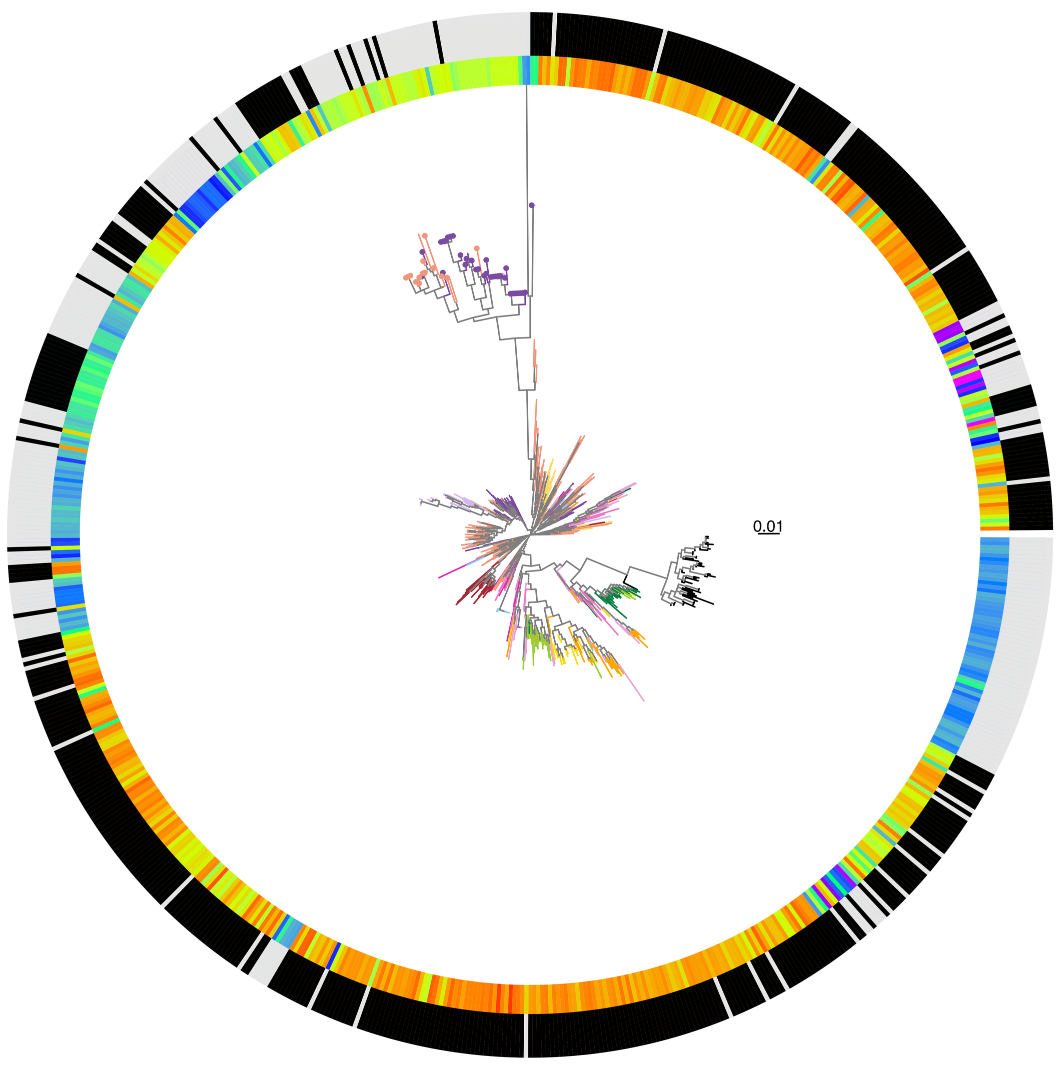


**HP0160**
**Value: 86**

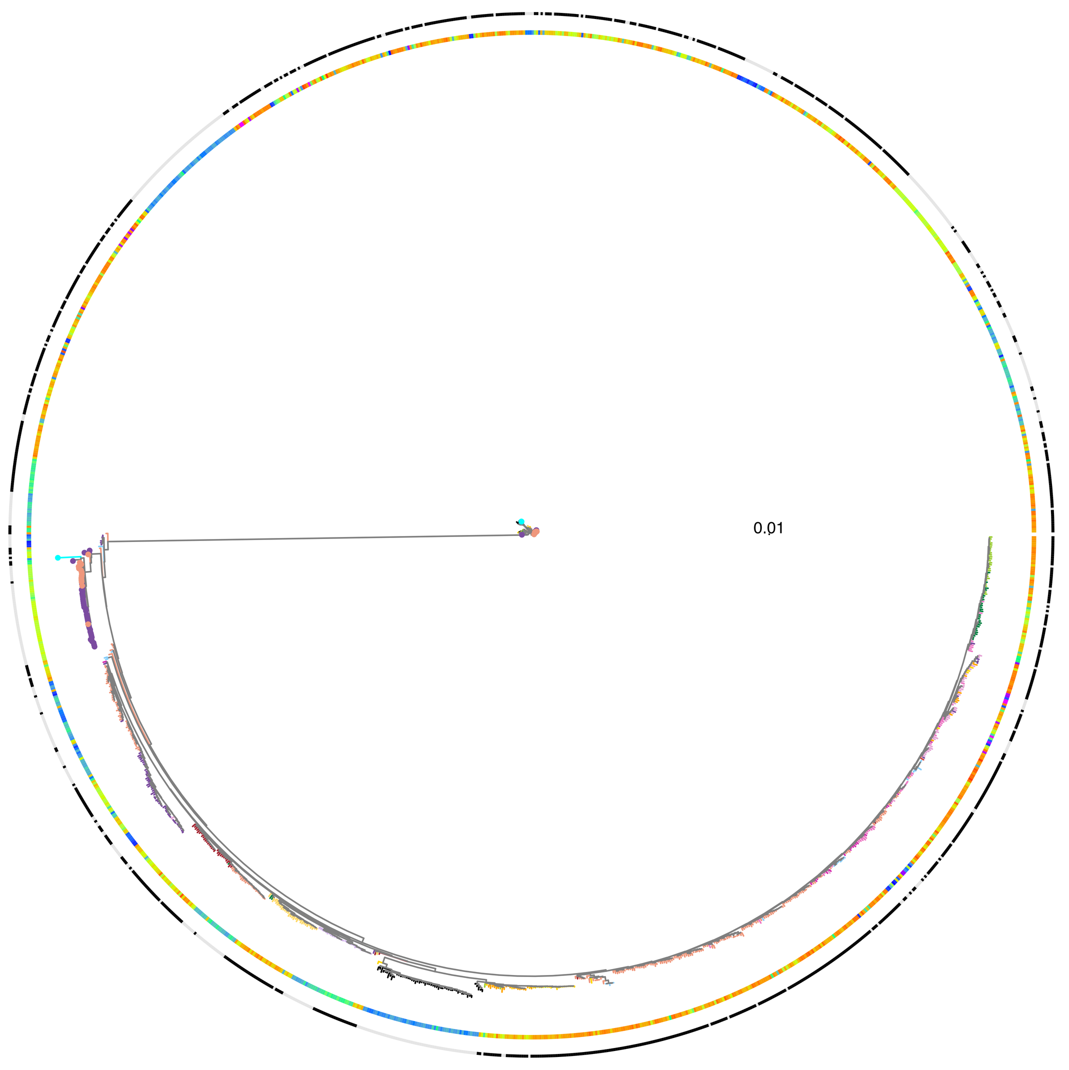


**HP0025**
**Value: 86**

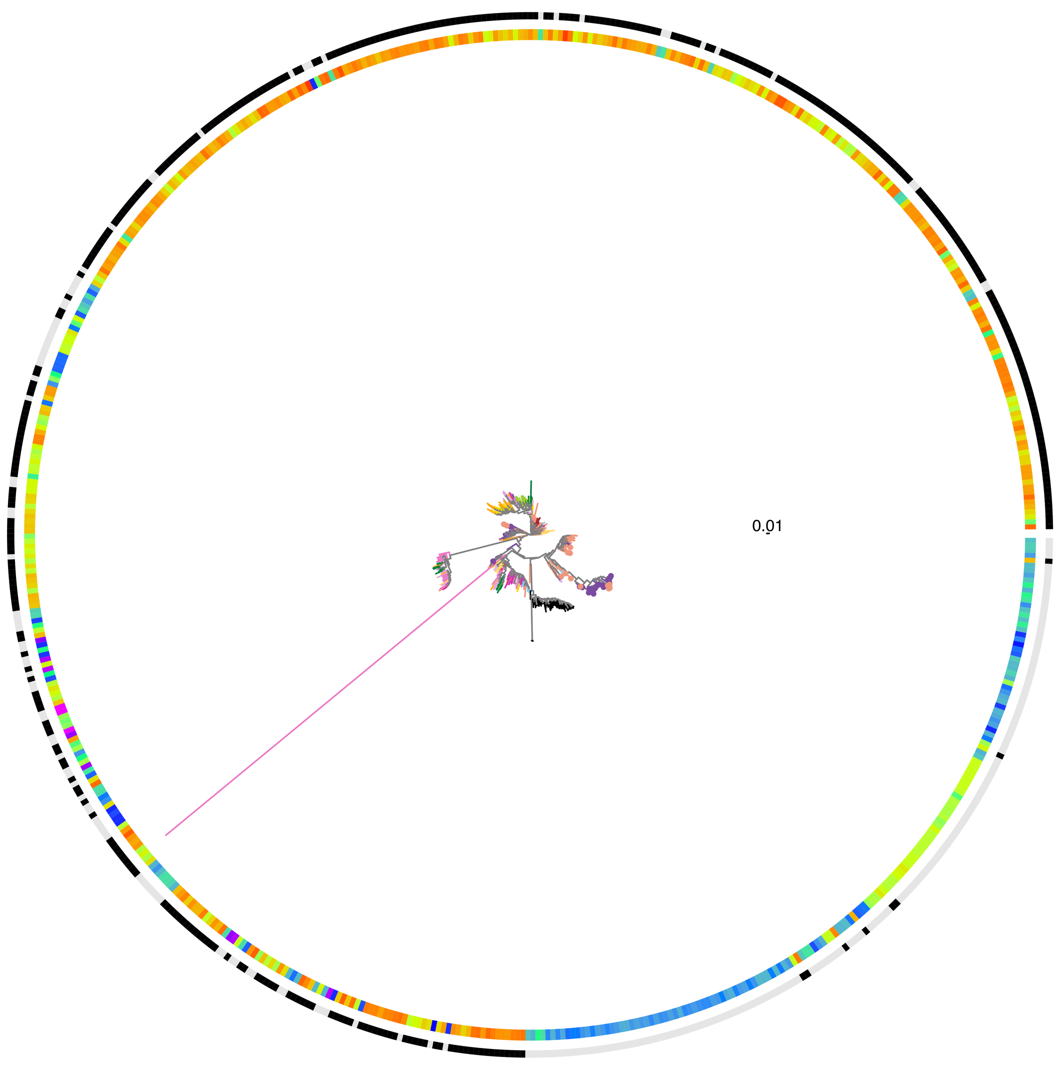

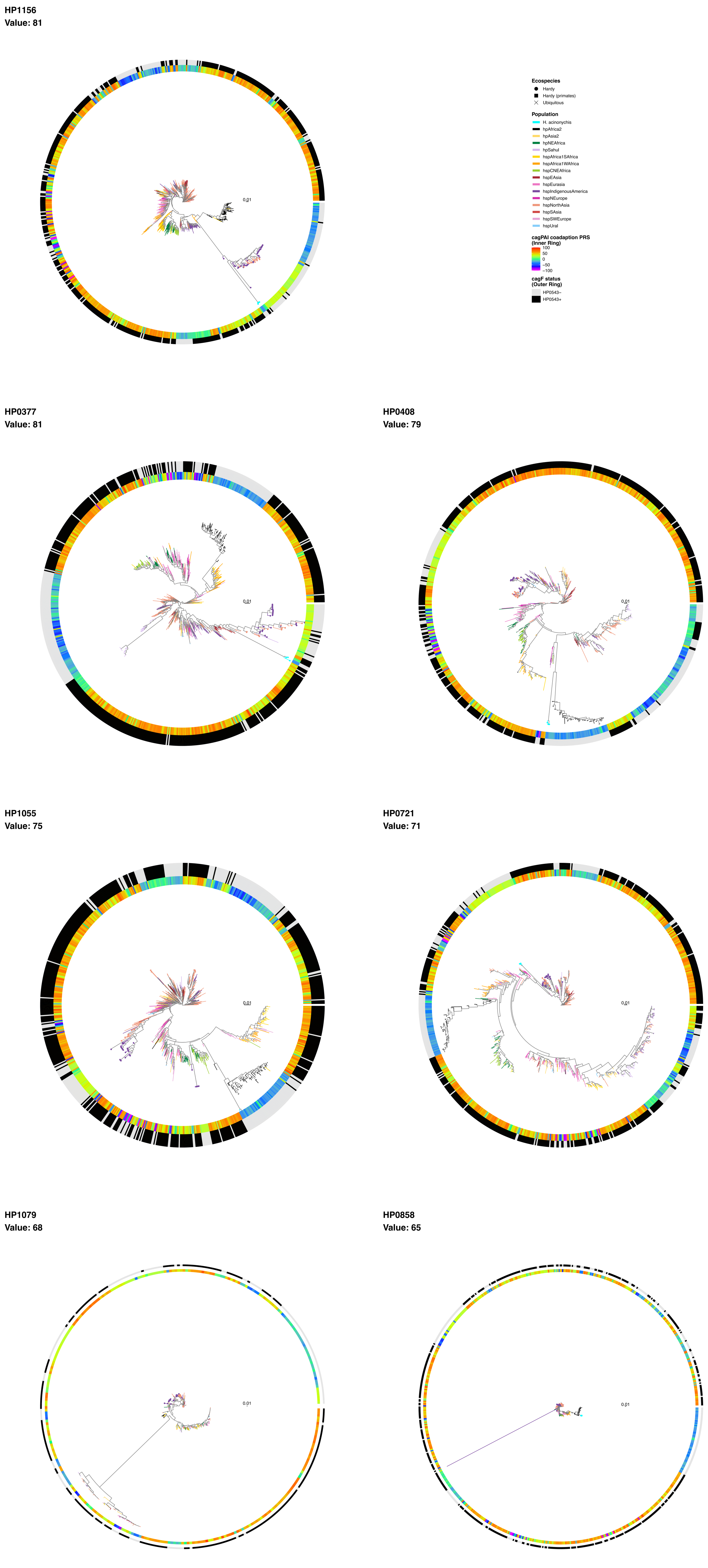
HP1156
Value: 81
0.01
Ecospecies
Hardy
Hardy (primates)
Ubiquitous
Population
H. acinonychis
hpAfrica2
hpAsia2
hpNEAfrica
hpSahul
hspAfrica1SAfrica
hspAfrica1WAfrica
hspCNEAfrica
hspEAsia
hspEurasia
hspIndigenousAmerica
hspNEurope
hspNorthAsia
hspSAsia
hspSWEurope
hspUral
cagPAI coadaption PRS
(Inner Ring)
100
50
0
−50
−100
cagF status
(Outer Ring)
HP0543−
HP0543+
HP0377
Value: 81
0.01
HP0408
Value: 79
0.01
HP1055
Value: 75
0.01
HP0721
Value: 71
0.01
HP1079
Value: 68
0.01
HP0858
Value: 65
0.01

**HP0710**
**Value: 65**

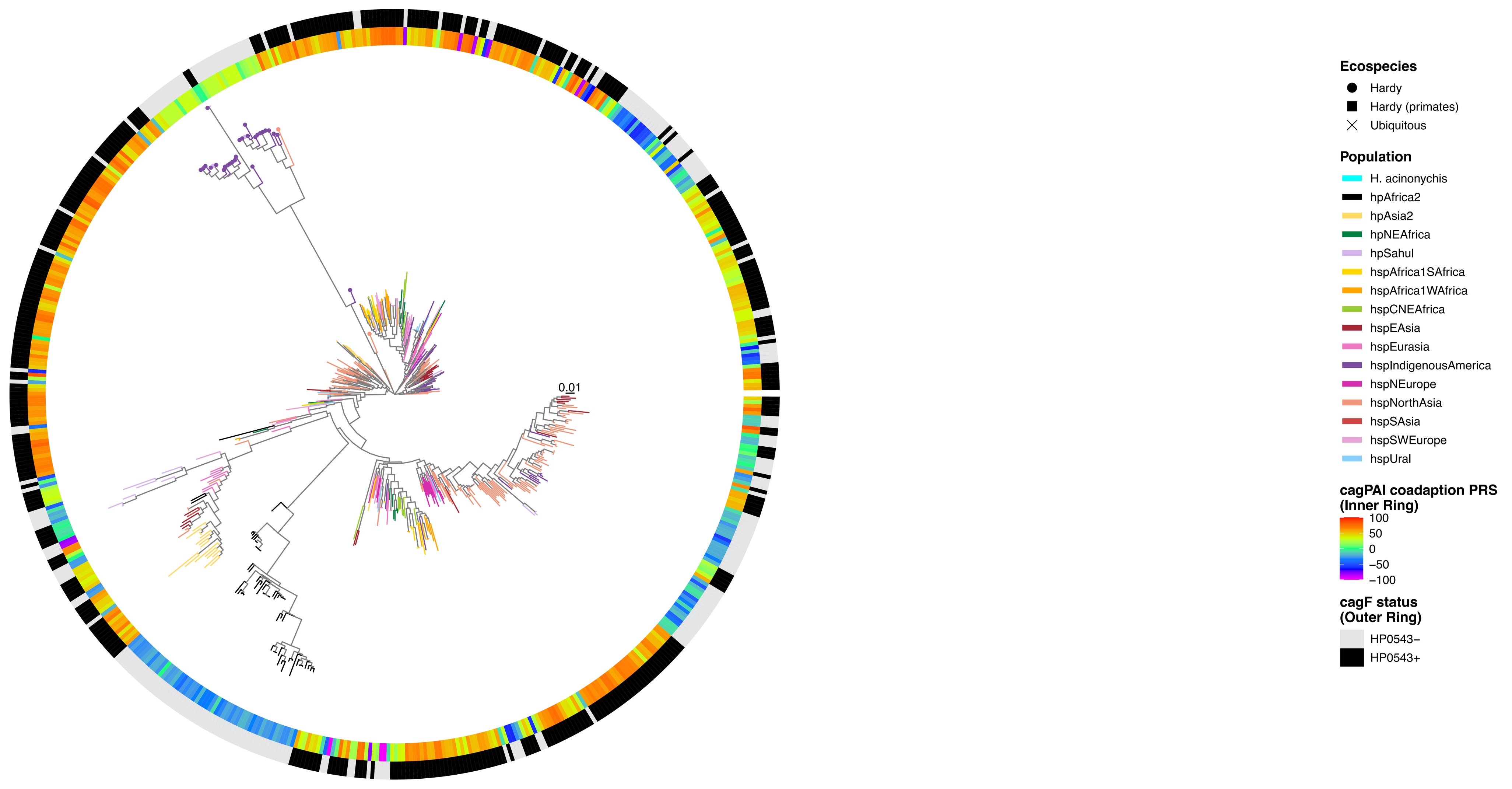


**HP0896**
**Value: 63**

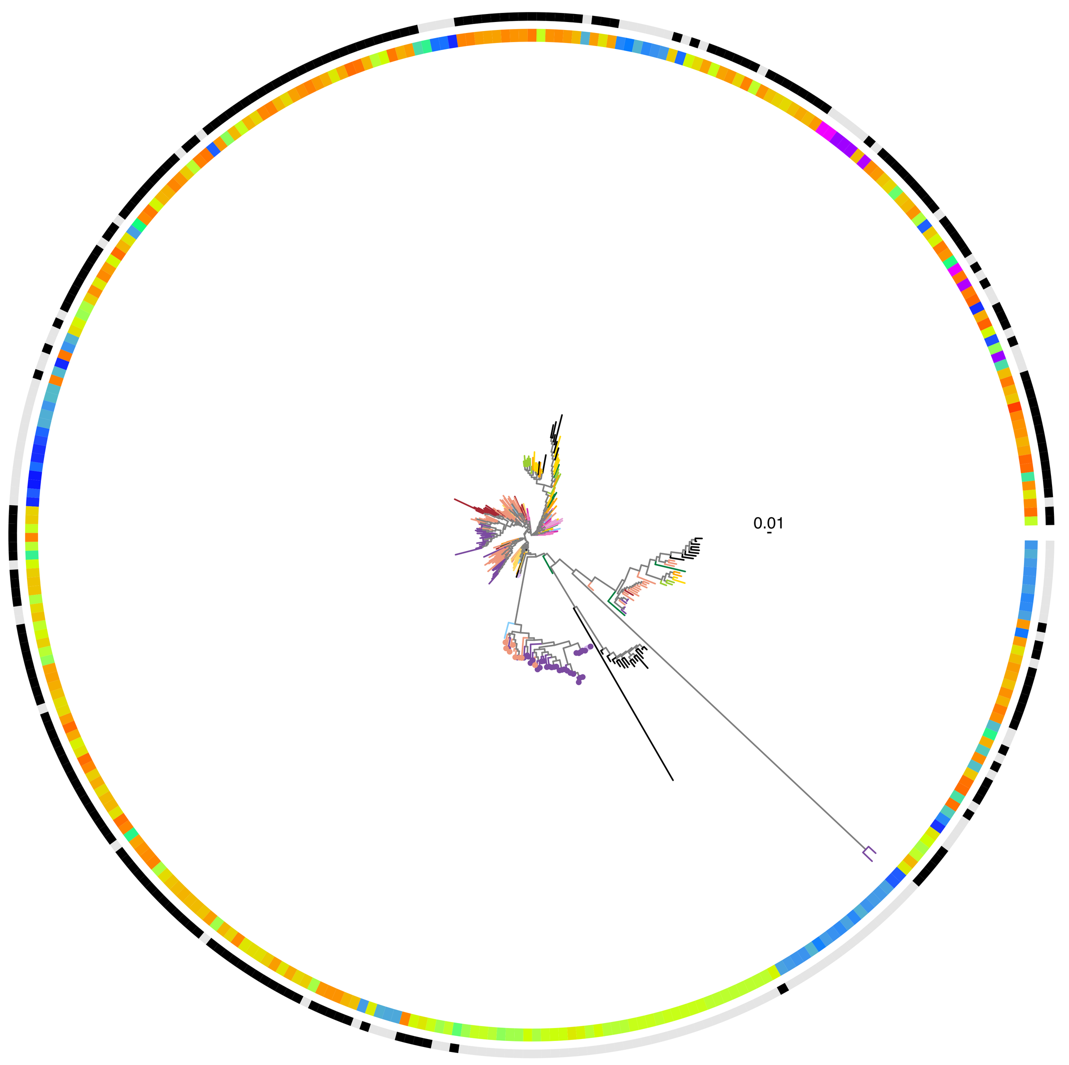


**HP0305**
**Value: 62**

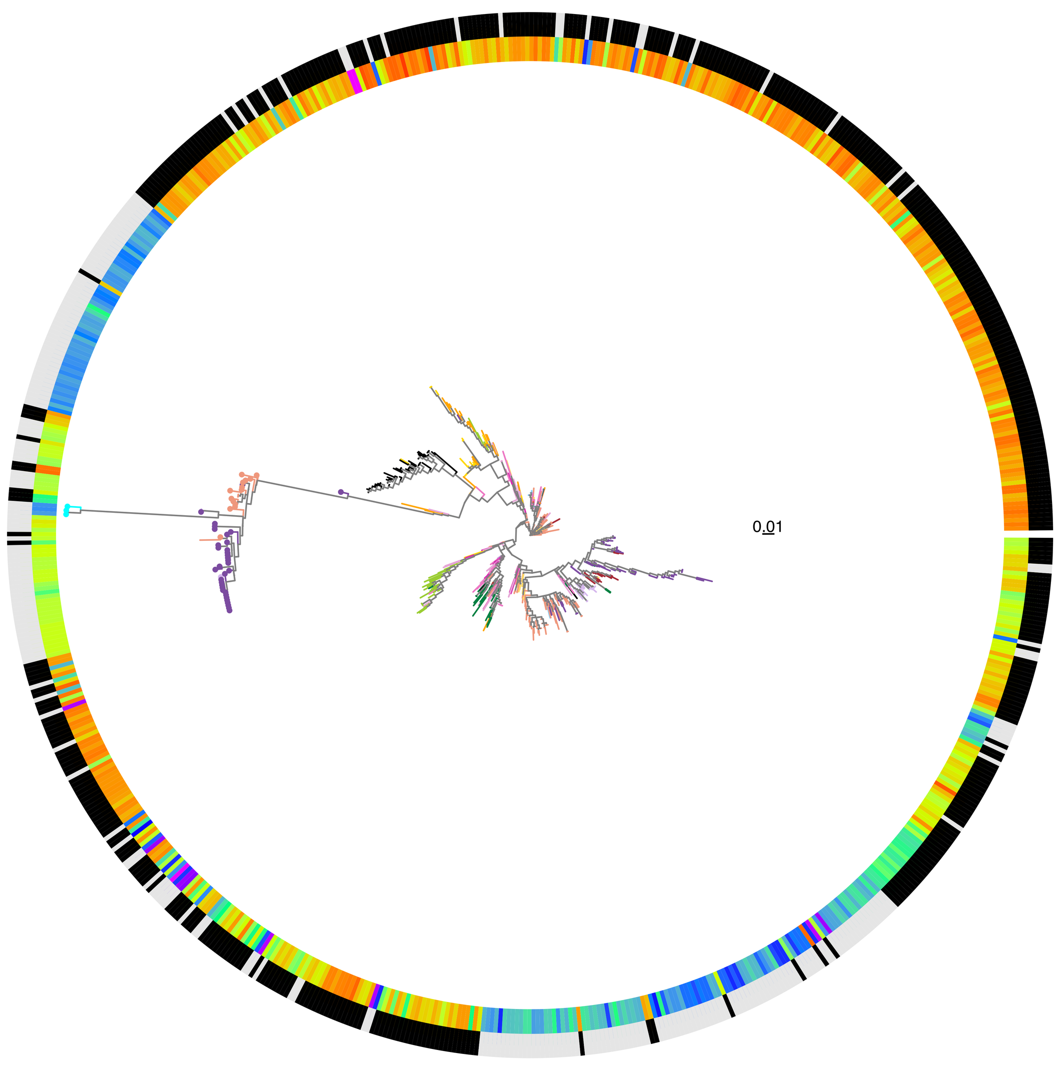


**HP0597**
**Value: 57**

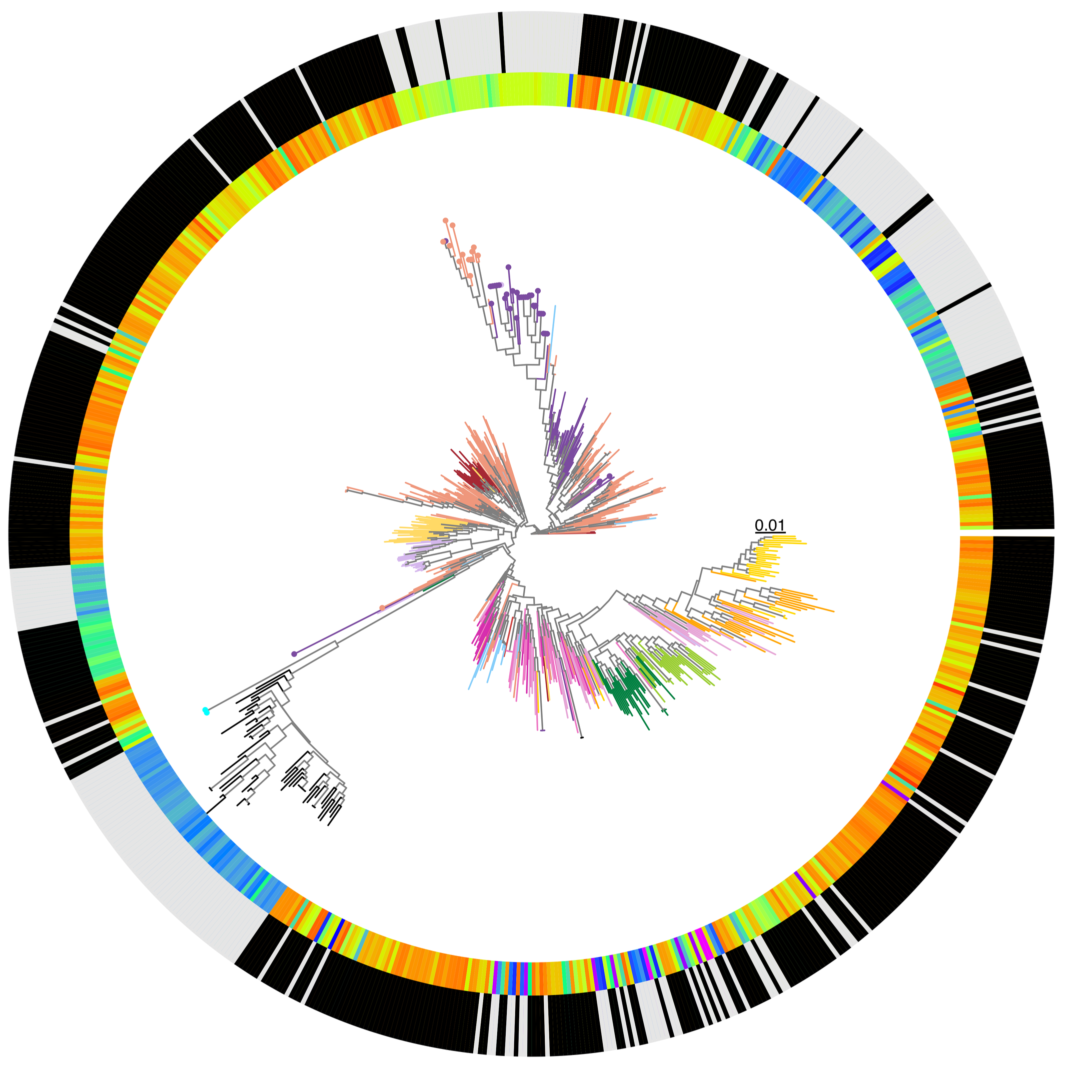


**HP0486**
**Value: 55**

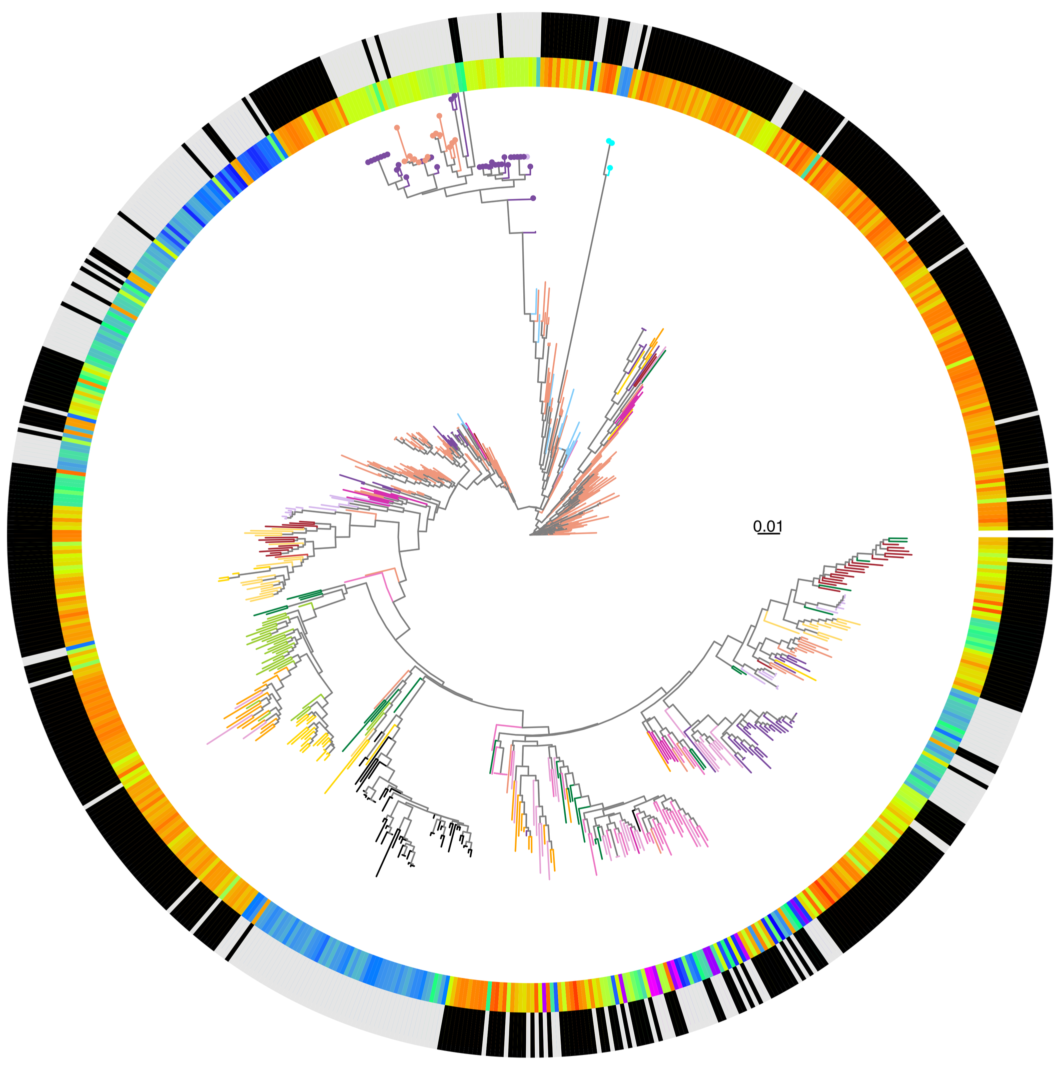


**HP0079**
**Value: 52**

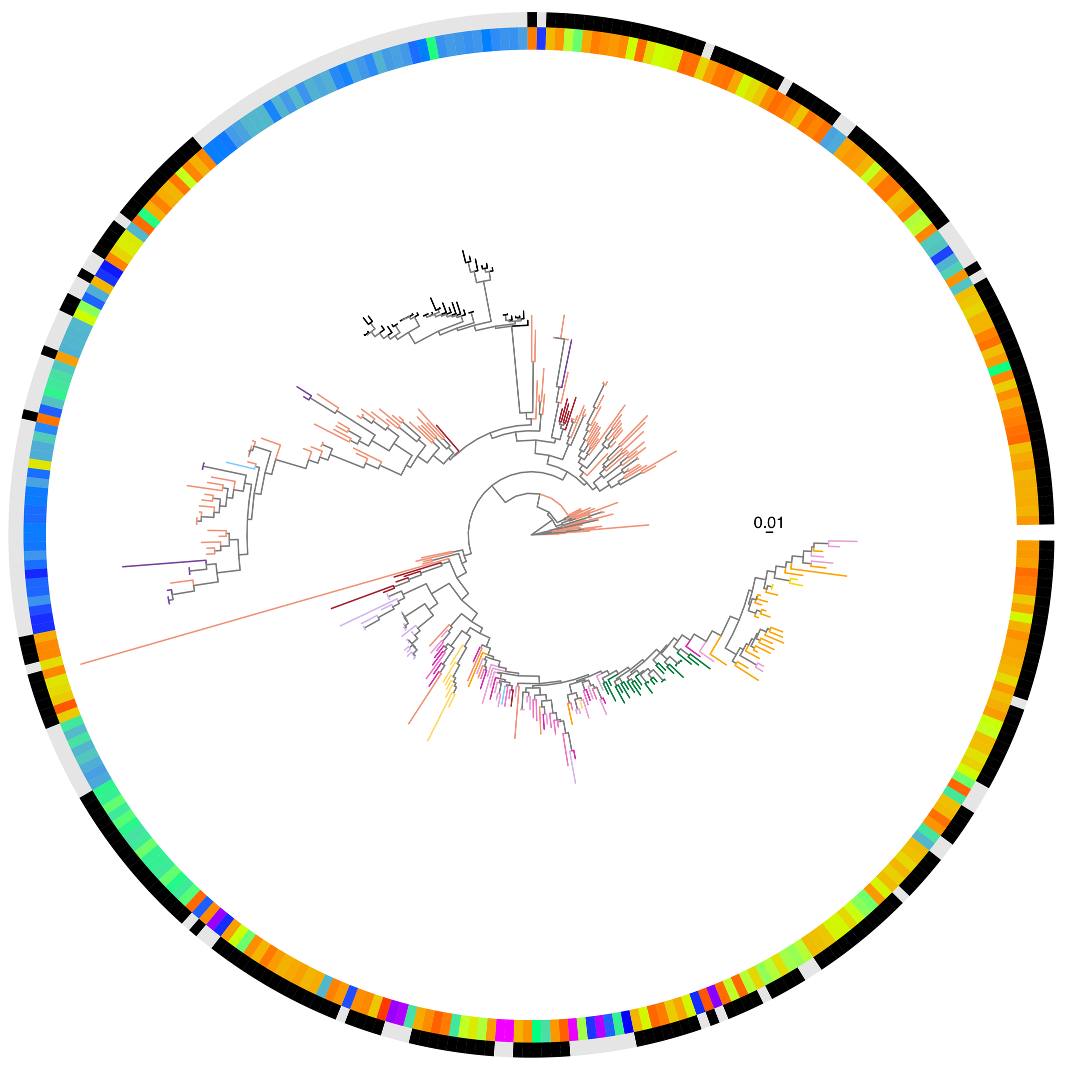


**HP0913**
**Value: 51**

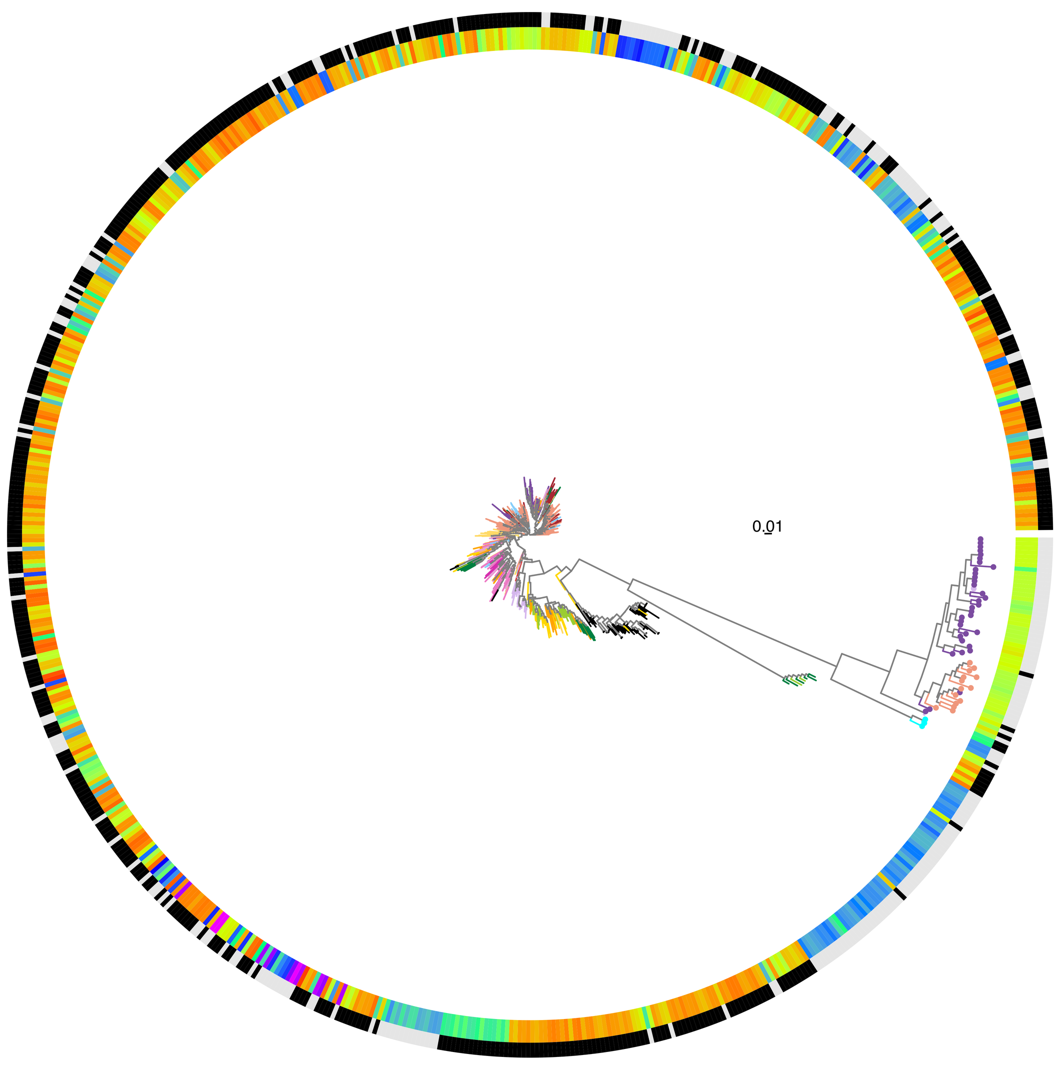

HP0723
Value: 50

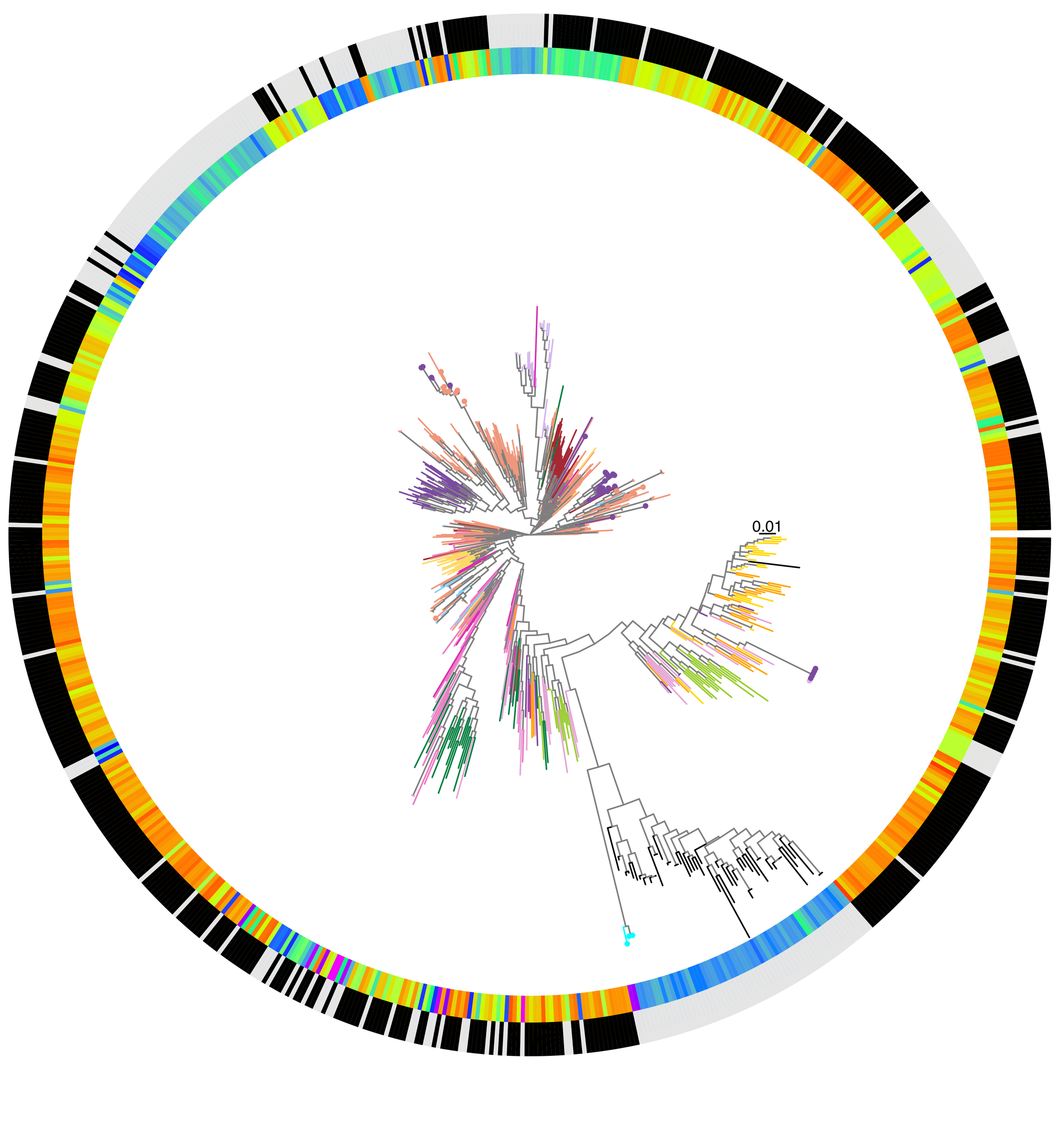


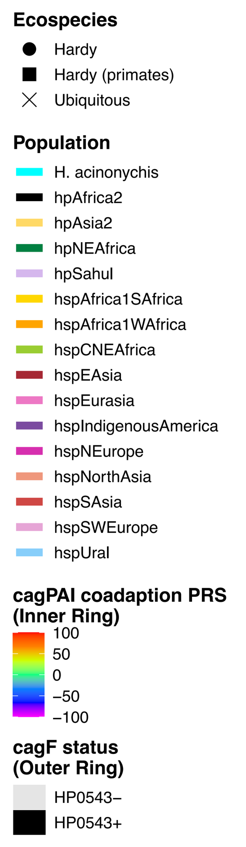


HP0252
Value: 49

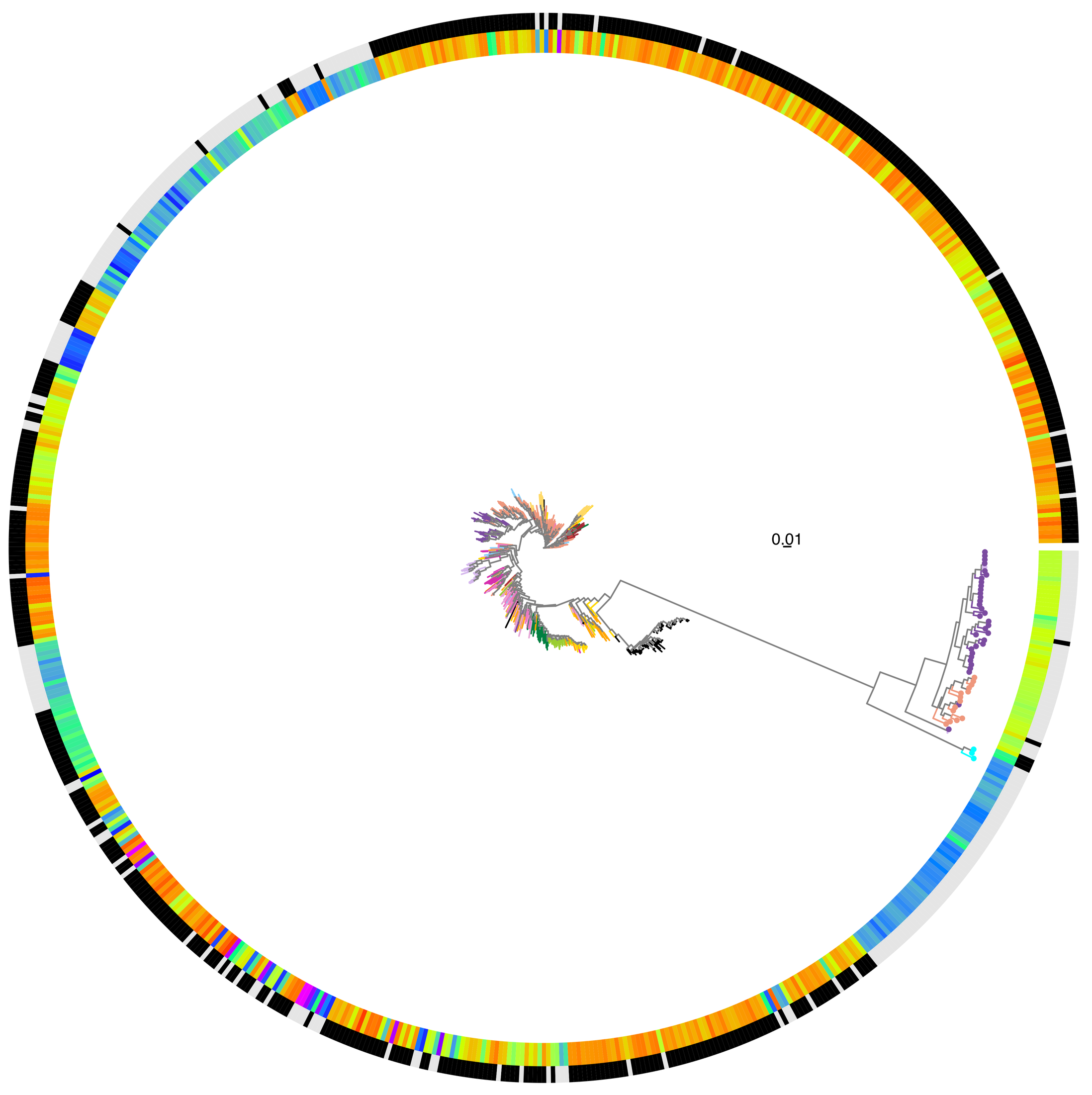


HP0561
Value: 45

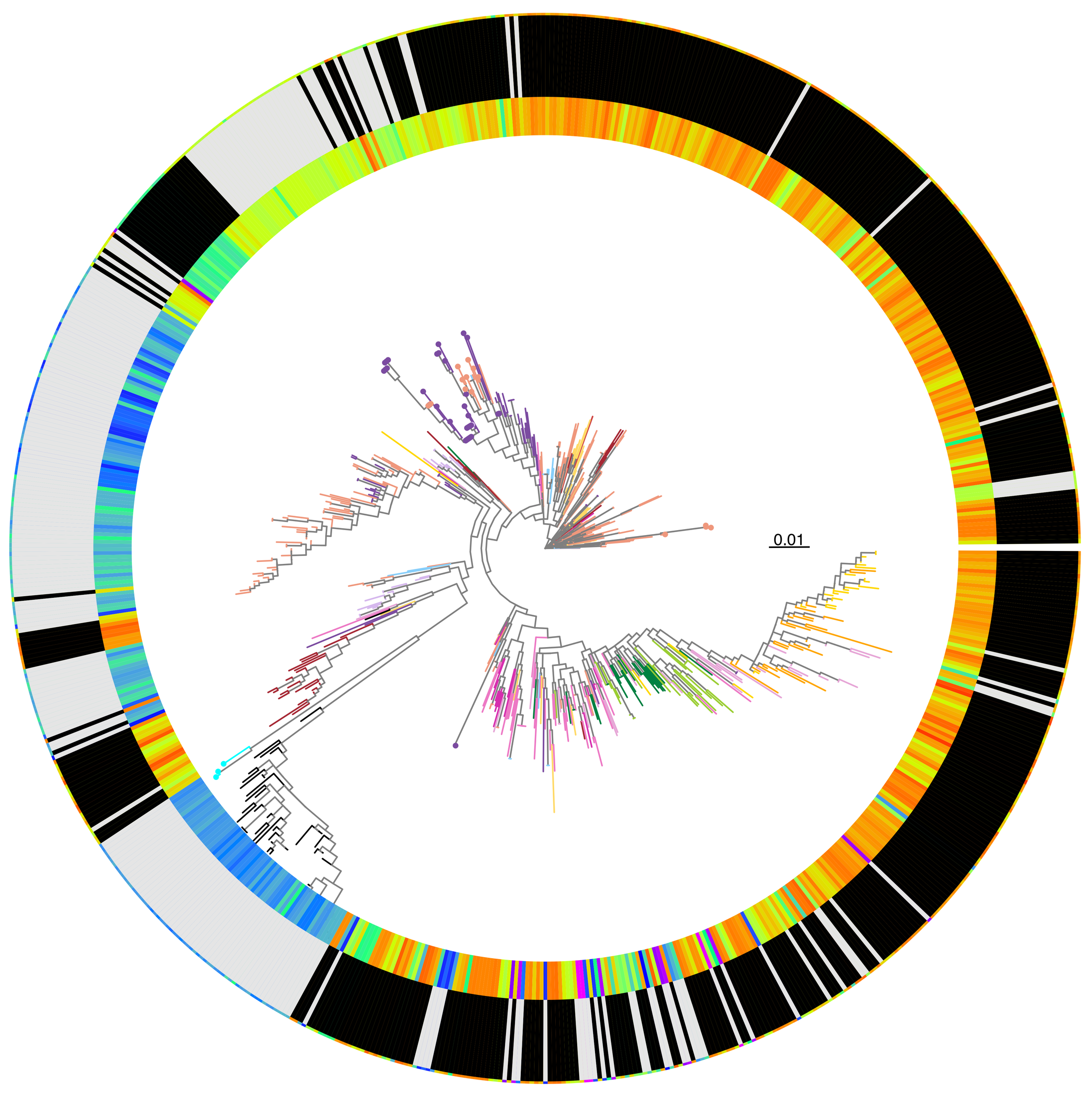


HP0608
Value: 42

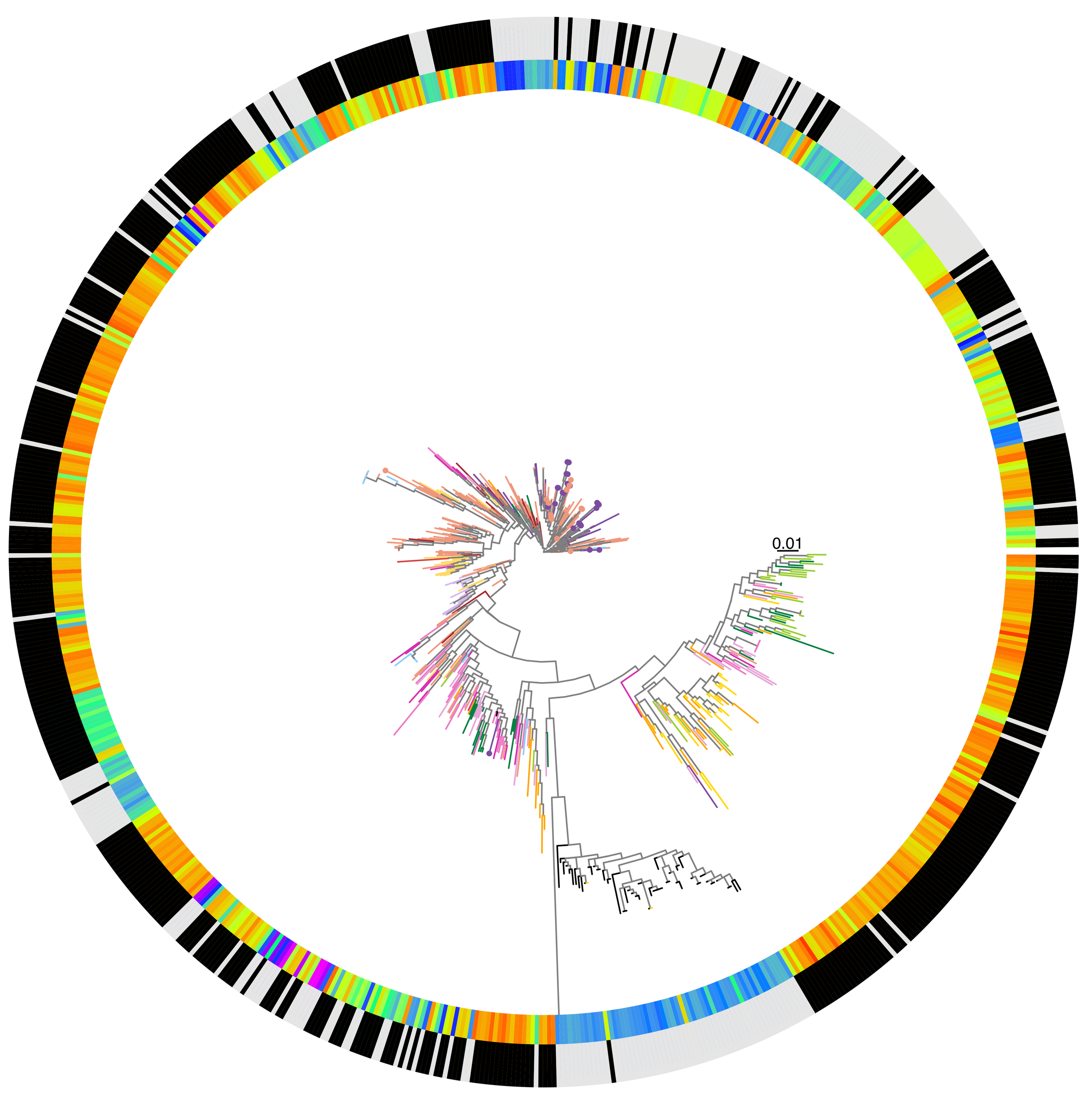
