## Extended_Data_Fig_4 for "History of coadaptation to the *cag* pathogenicity island in *Helicobacter pylori*"

**HP0721 | HP0722**
**P−val: 216.0227**

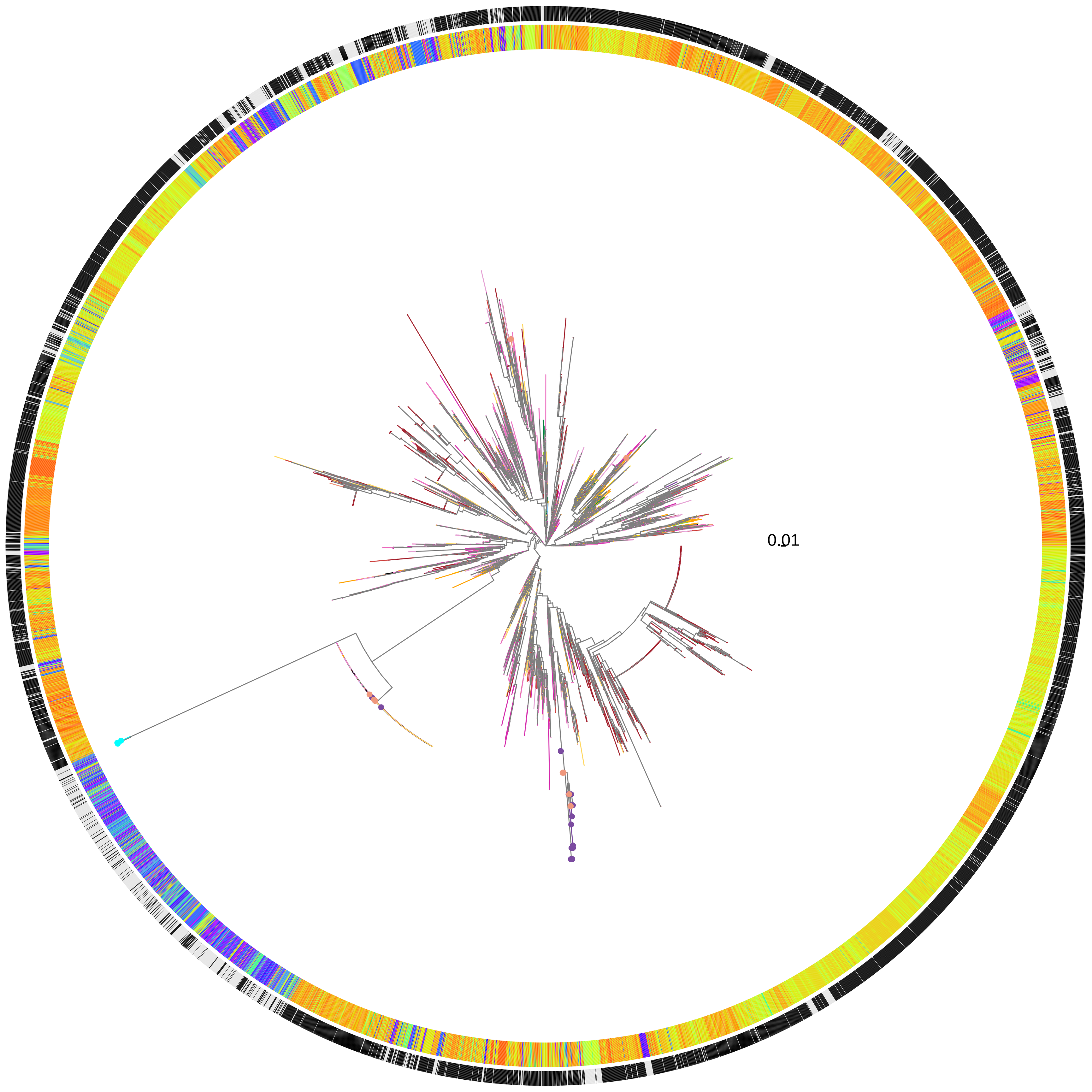


**HP0722 | HP0723**
**P−val: 215.0227**

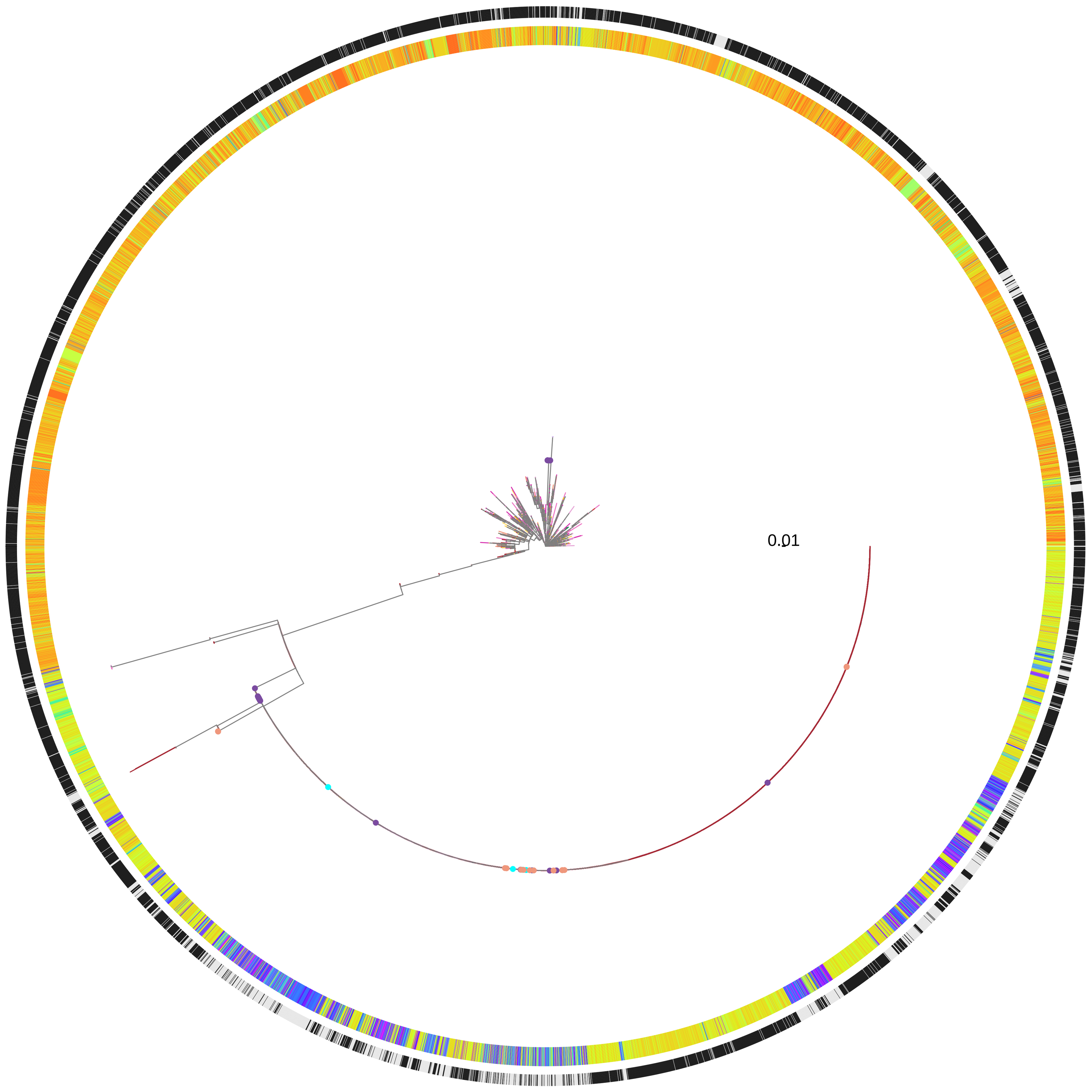


**Ecospecies**
Hardy
Hardy (primates)
Ubiquitous

**Population**
H. acinonychis
hpAfrica2
hpAsia2
hpNEAfrica
hpSahul
hspAfrica1SAfrica
hspAfrica1WAfrica
hspCNEAfrica
hspEAsia
hspEurasia
hspIndigenousAmerica
hspNEurope
hspNorthAsia
hspSAsia
hspSWEurope
hspUral

**Total gene adaption PRS (Inner Ring)**
100
50
0
−50
−100

**cagF status (Outer Ring)**
HP0543−
HP0543+

**HP0407 | HP0408**
**P−val: 106.0057**

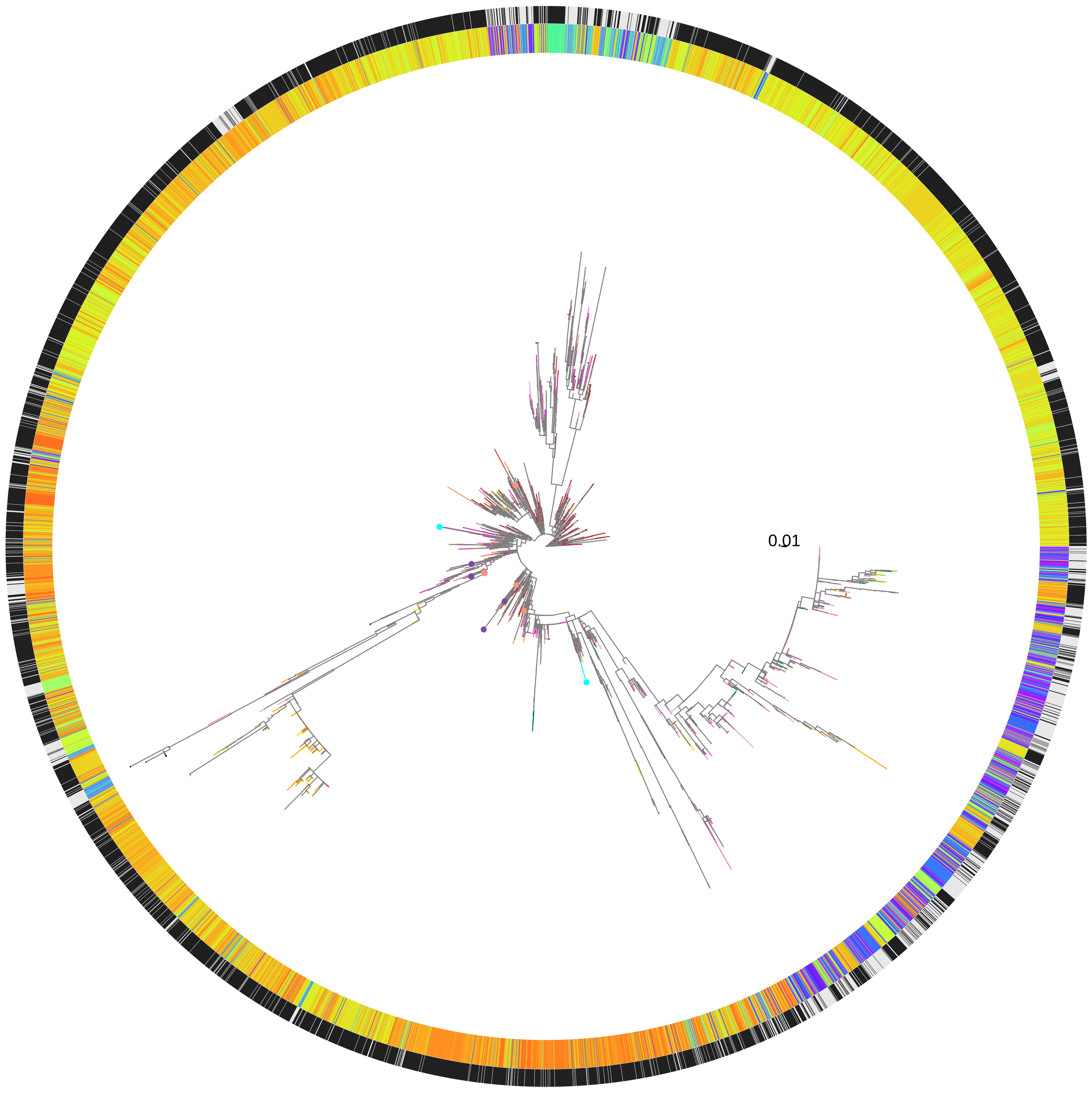


**HP0313 | HP0315**
**P−val: 98.1959**

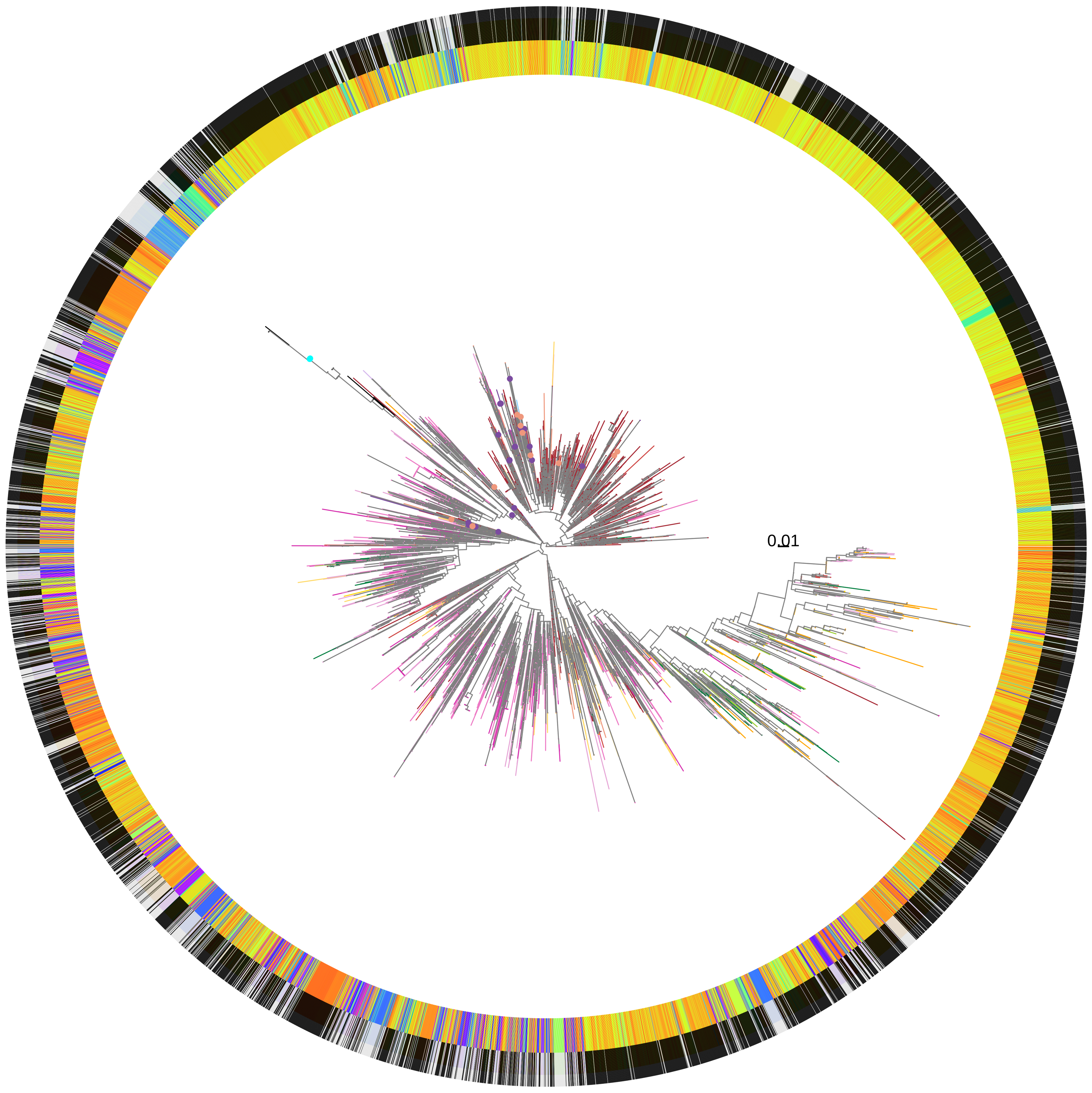


**Ecospecies**
- Hardy
- Hardy (primates)
- Ubiquitous

**Population**
- H. acinonychis
- hpAfrica2
- hpAsia2
- hpNEAfrica
- hpSahul
- hspAfrica1SAfrica
- hspAfrica1WAfrica
- hspCNEAfrica
- hspEAsia
- hspEurasia
- hspIndigenousAmerica
- hspNEurope
- hspNorthAsia
- hspSAsia
- hspSWEurope
- hspUral

**Total gene adaption PRS (Inner Ring)**
100
50
0
−50
−100

**cagF status (Outer Ring)**
- HP0543−
- HP0543+

**HP0855 | HP0856**

**P−val: 95.0022**

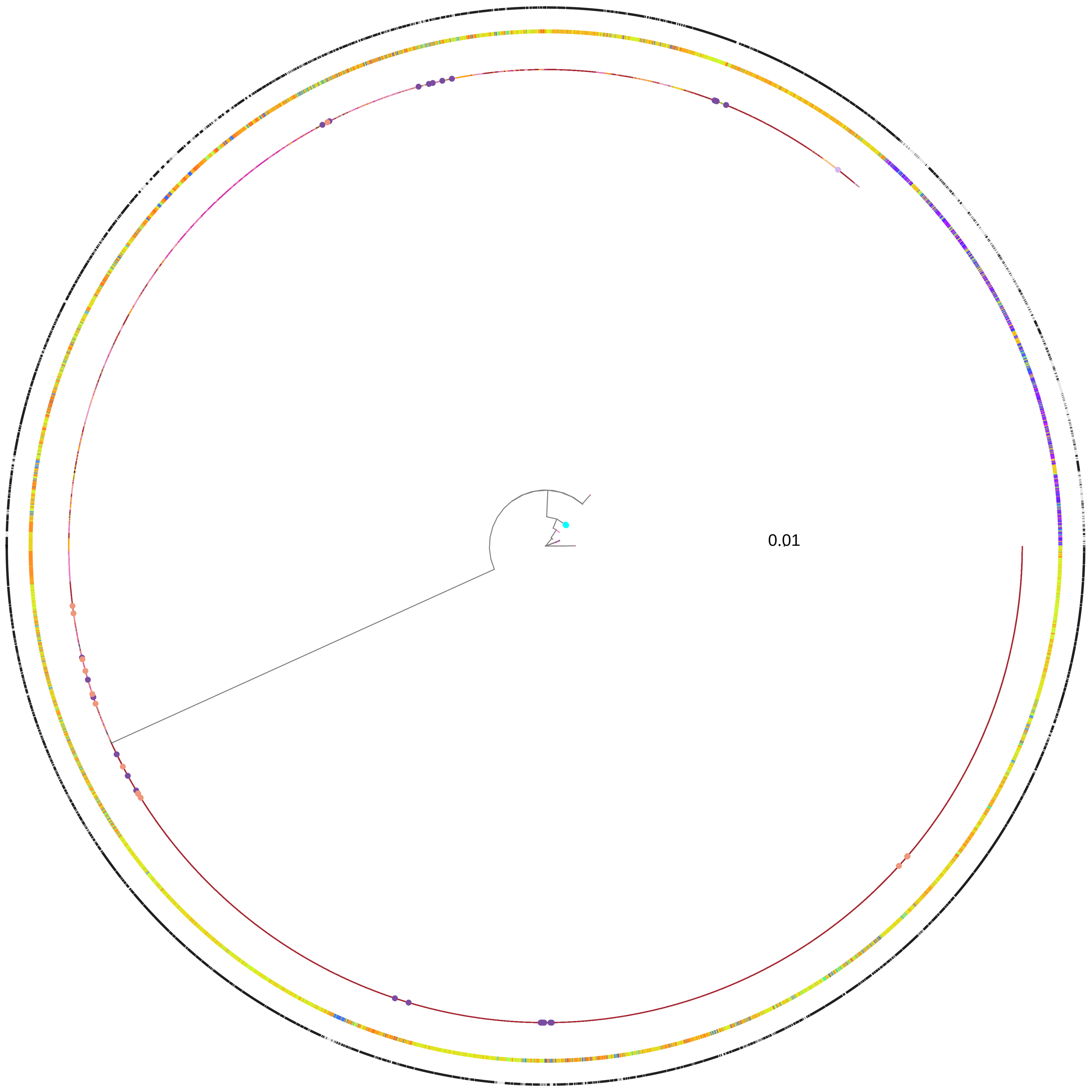


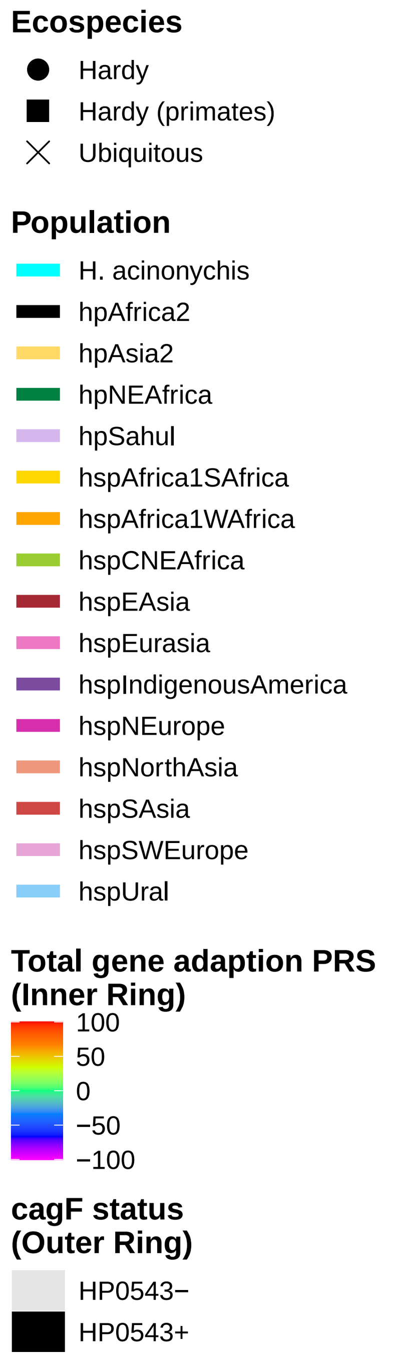


**HP0886 | HP0887**

**P−val: 93.4271**

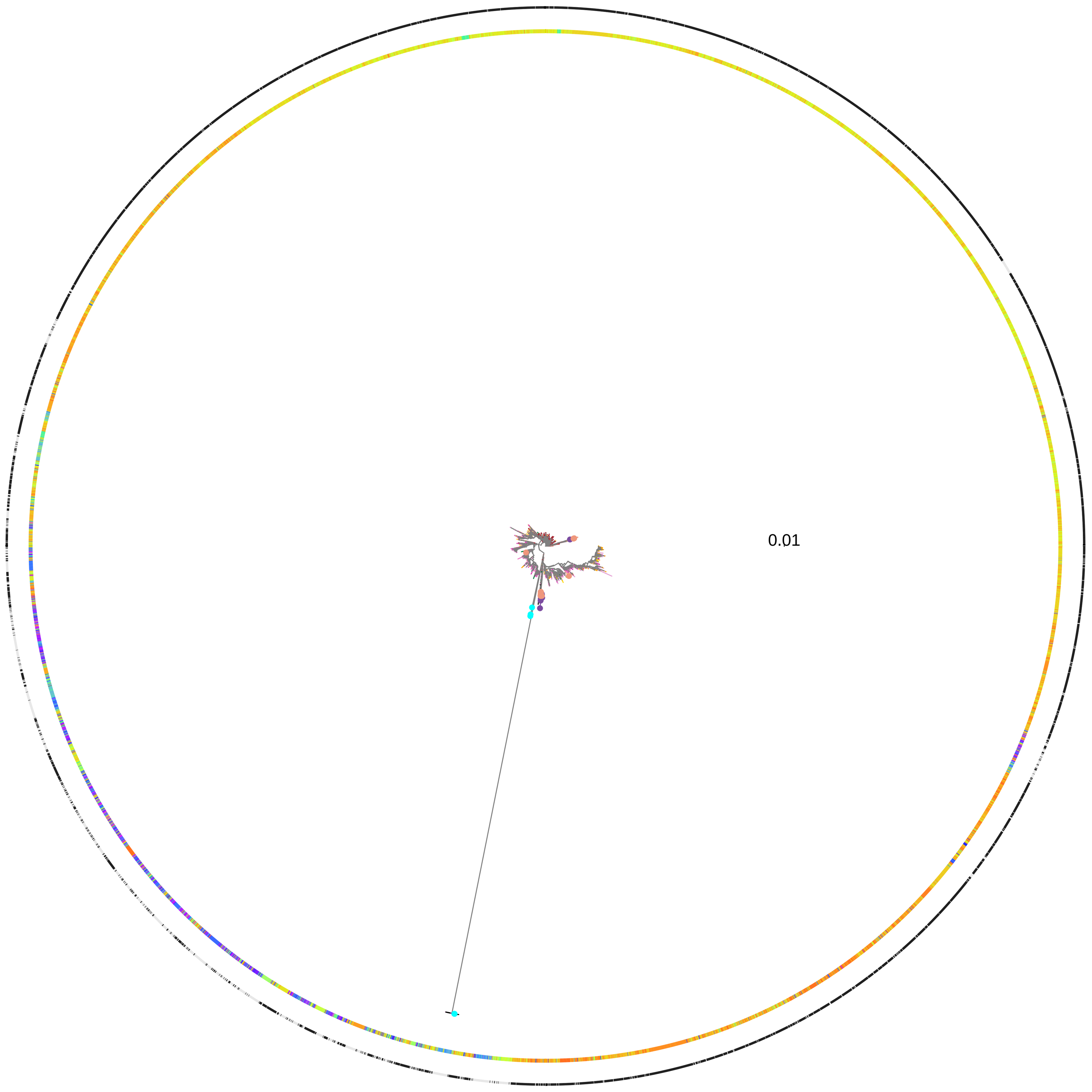

**HP0854 | HP0855**
**P−val: 83.2**

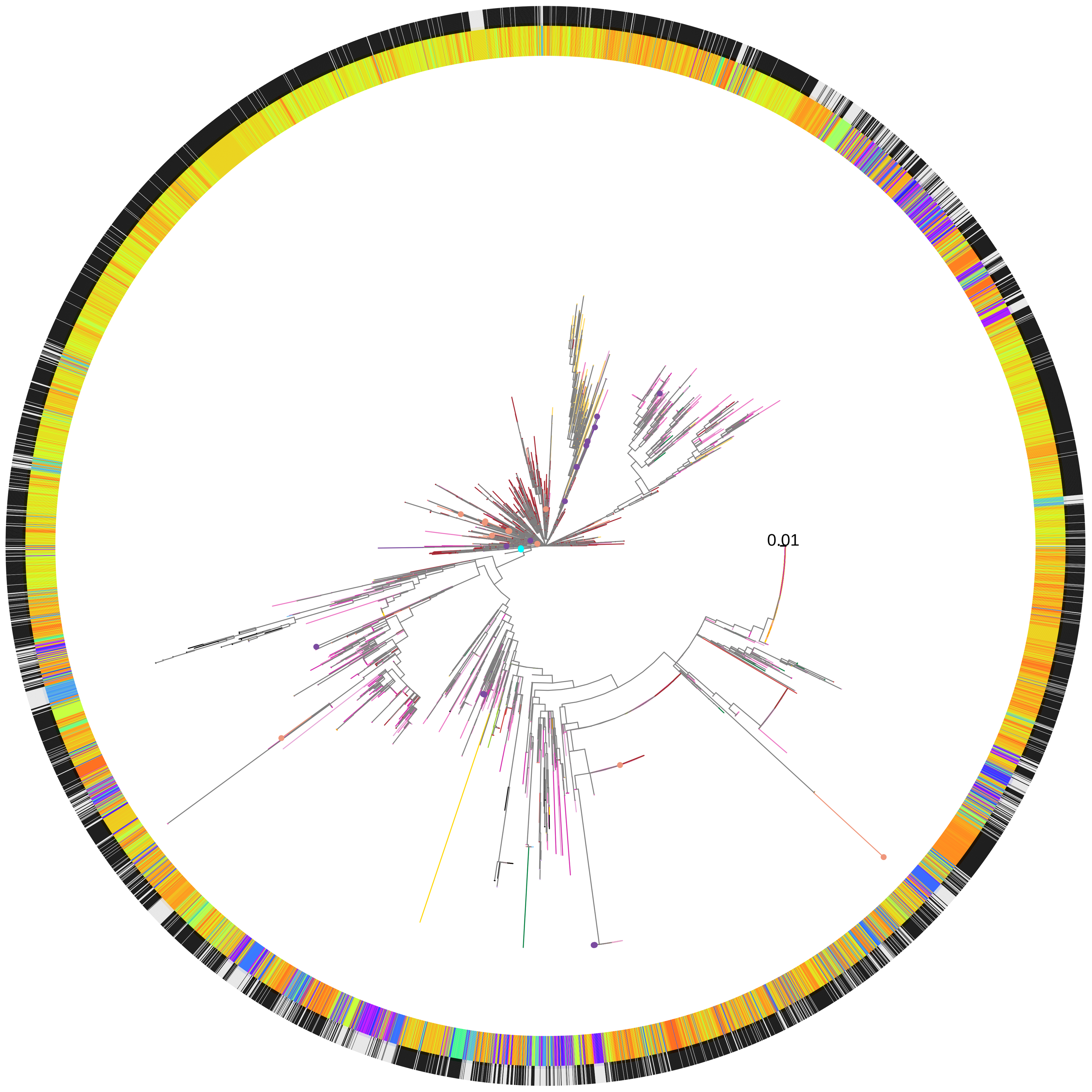


**HP0709 | HP0710**
**P−val: 76.289**

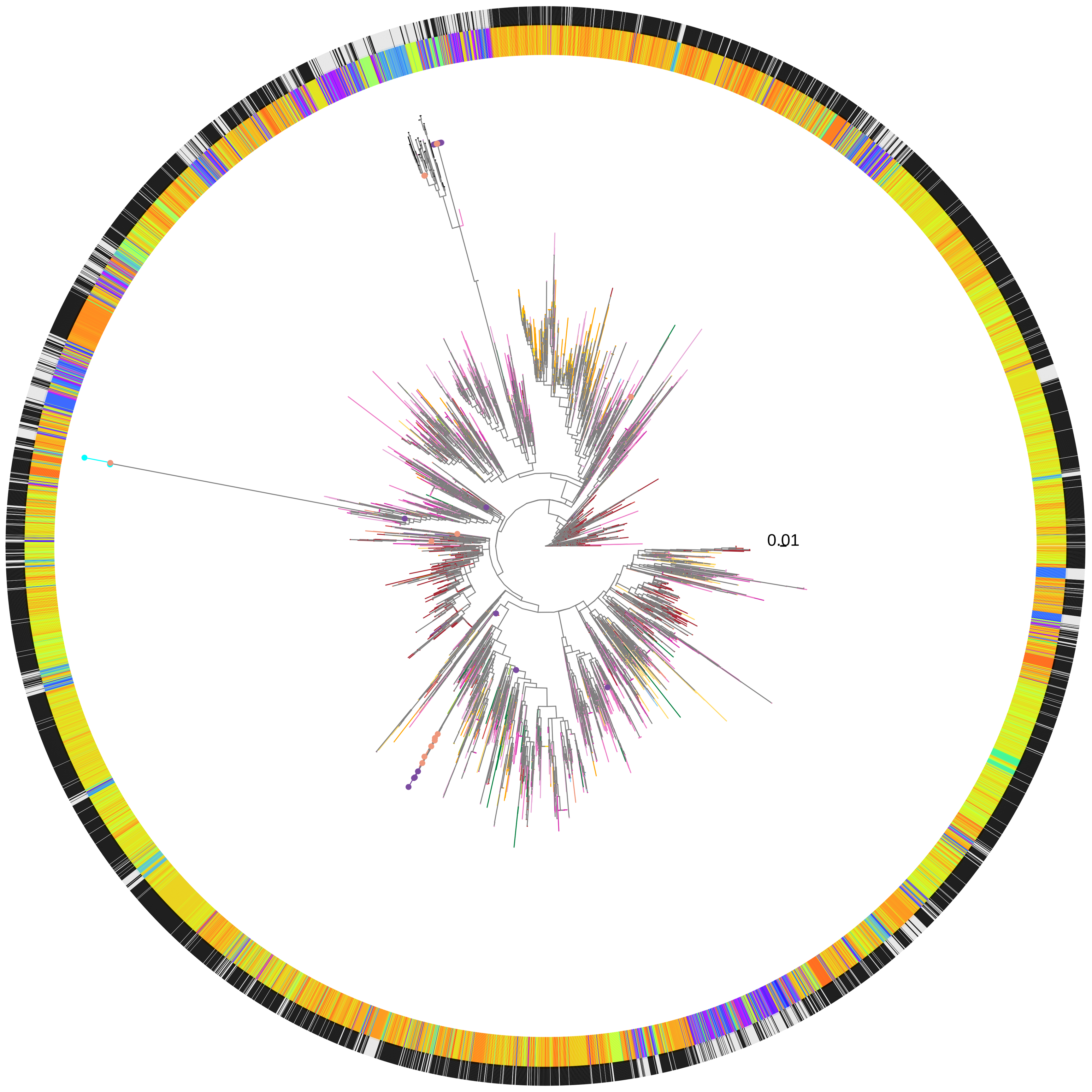

**HP0510 | HP0511**
**P−val: 70.0132**

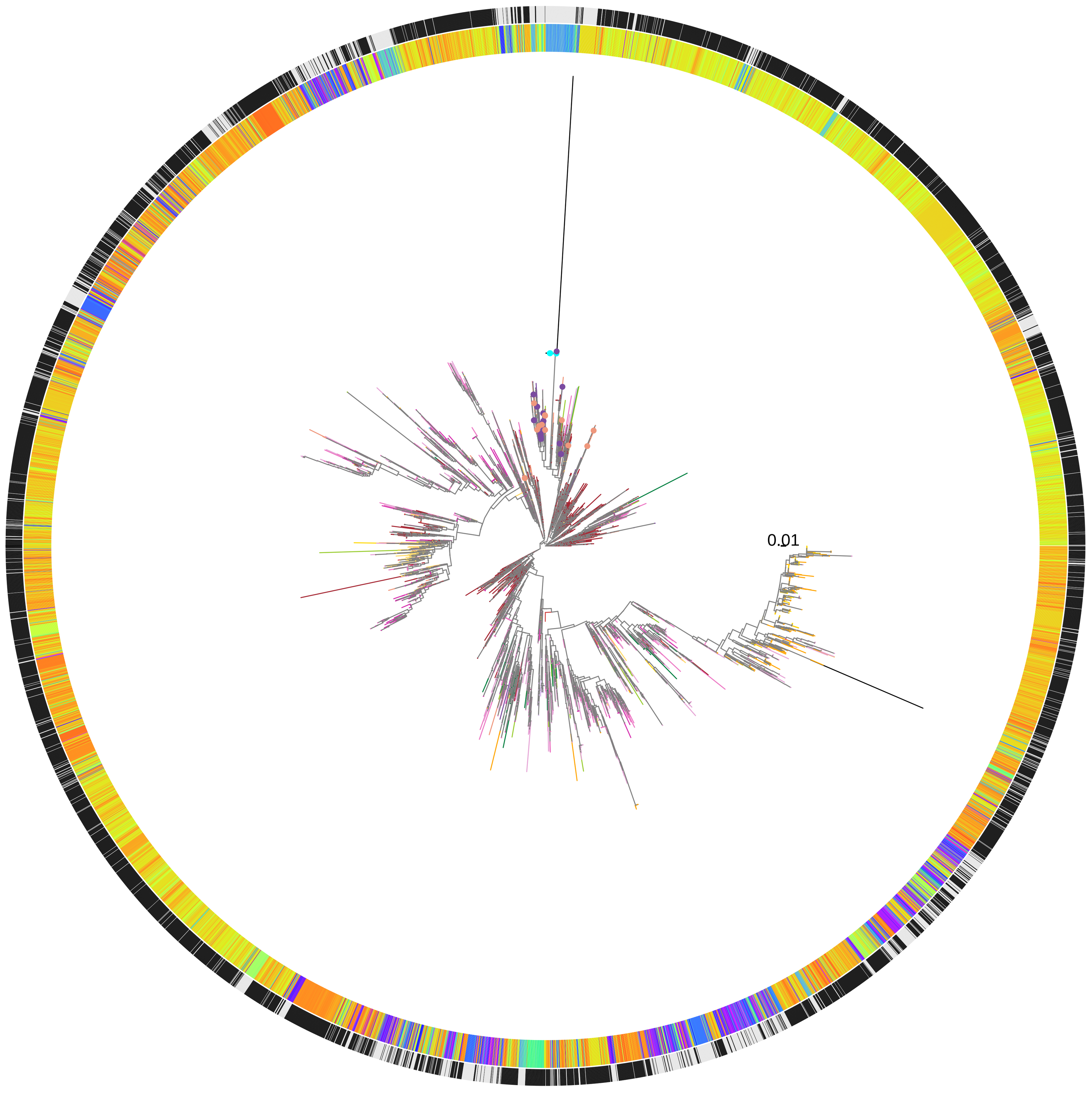


**HP1243 | HP1244**
**P−val: 68.767**

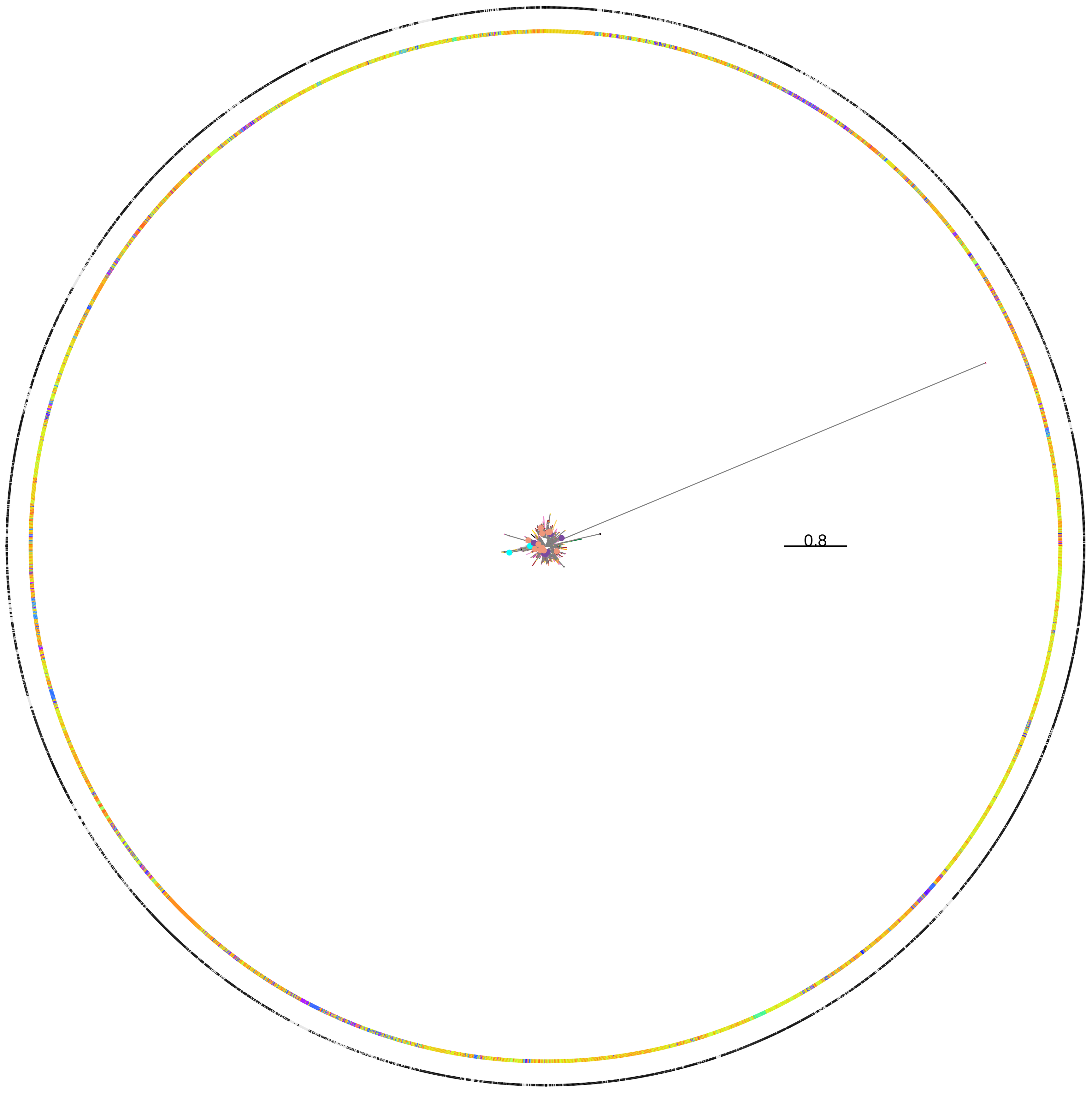


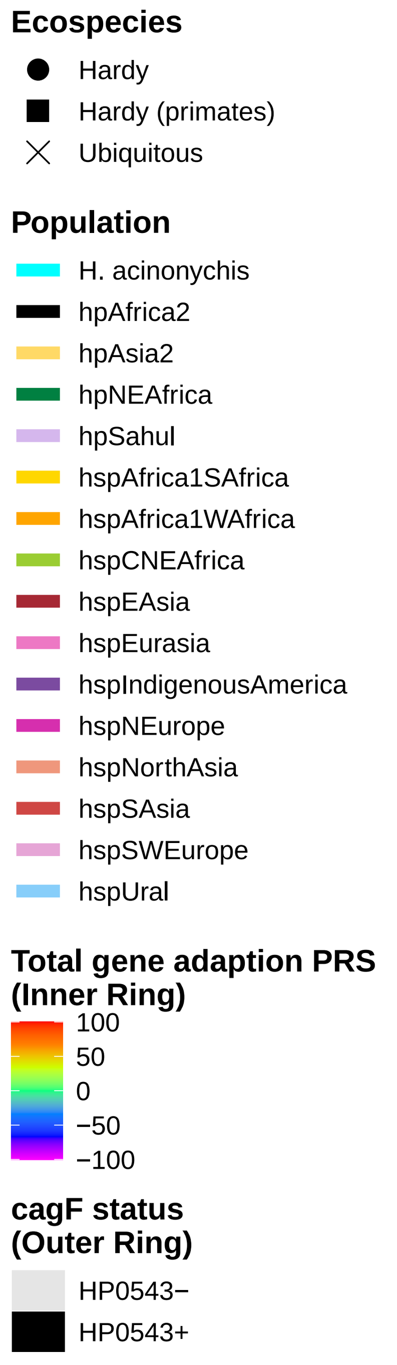

**HP1079 | HP1080**

**P−val: 63.5376**

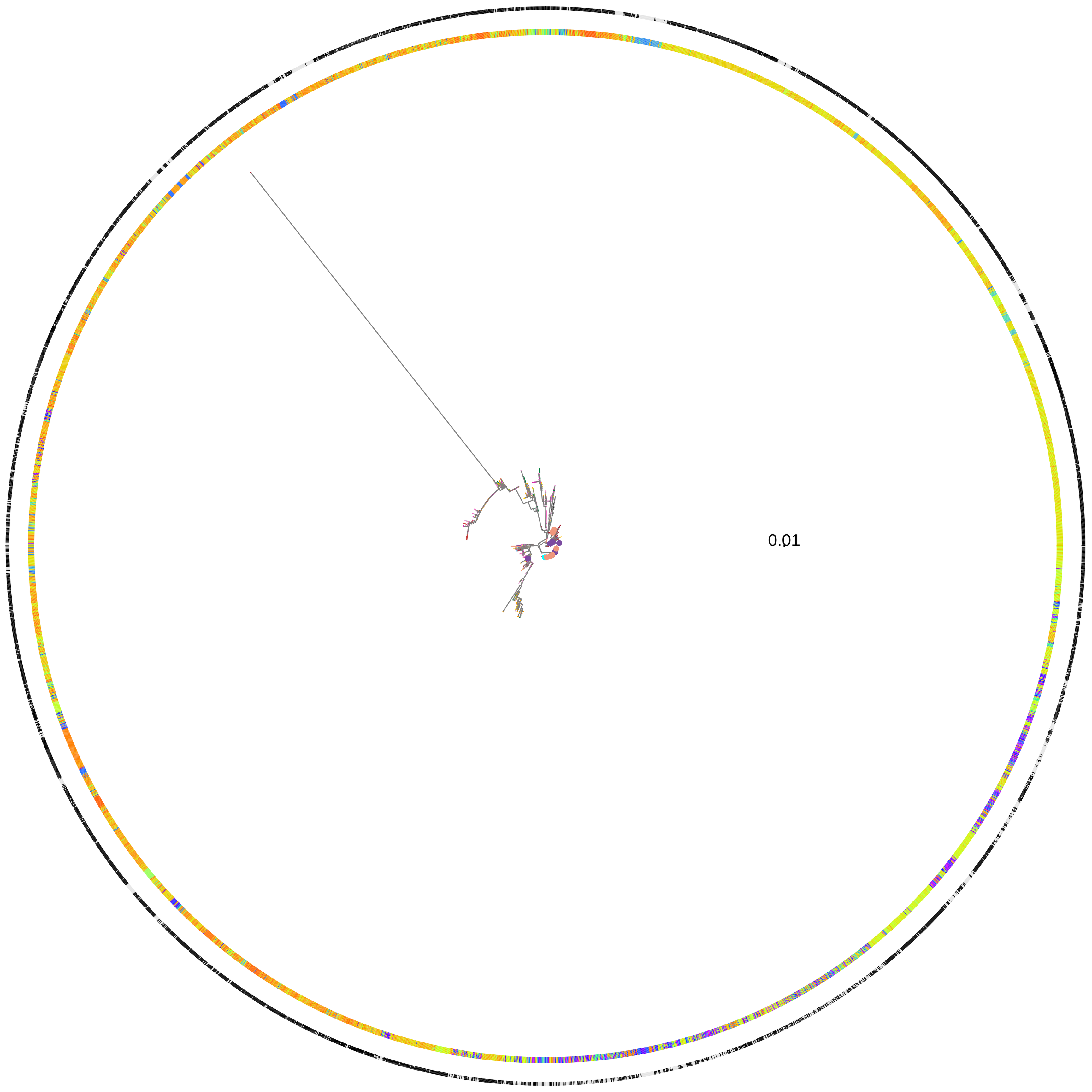


**HP0856 | HP0857**

**P−val: 58.4522**

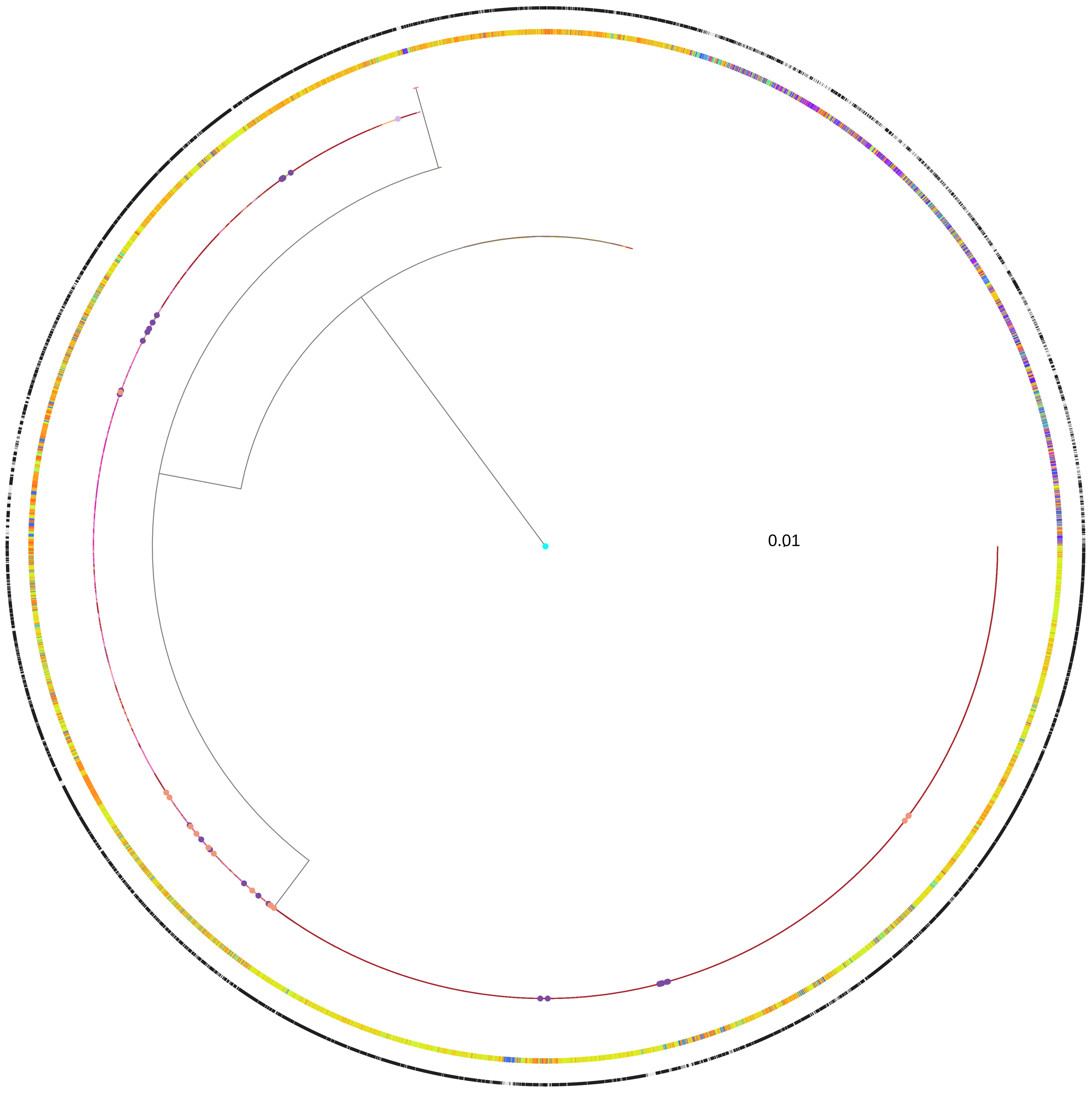

**HP1417m_2 | HP1418**
**P−val: 57.2306**

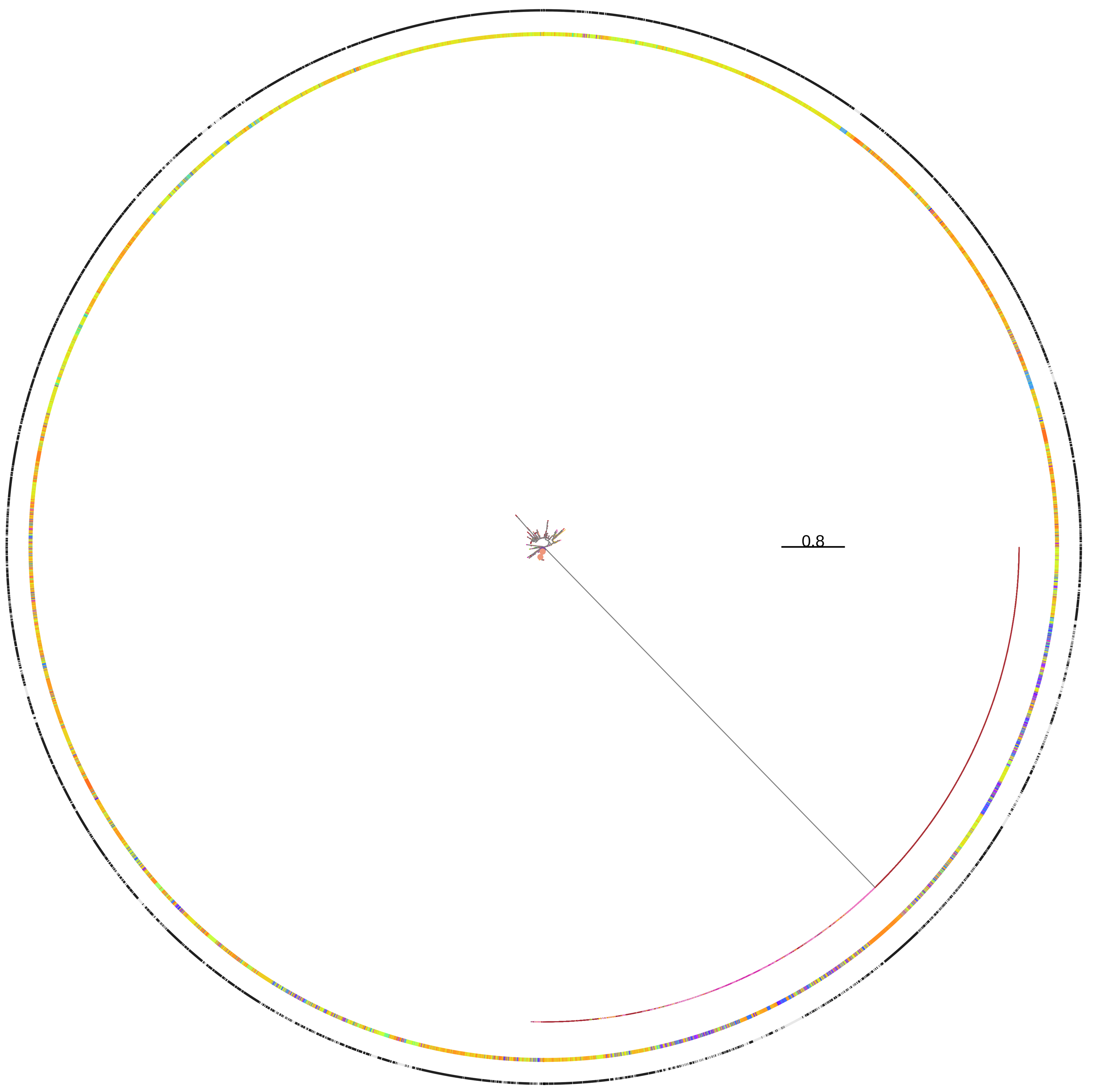


**HP0561 | HP0562**
**P−val: 52.5702**

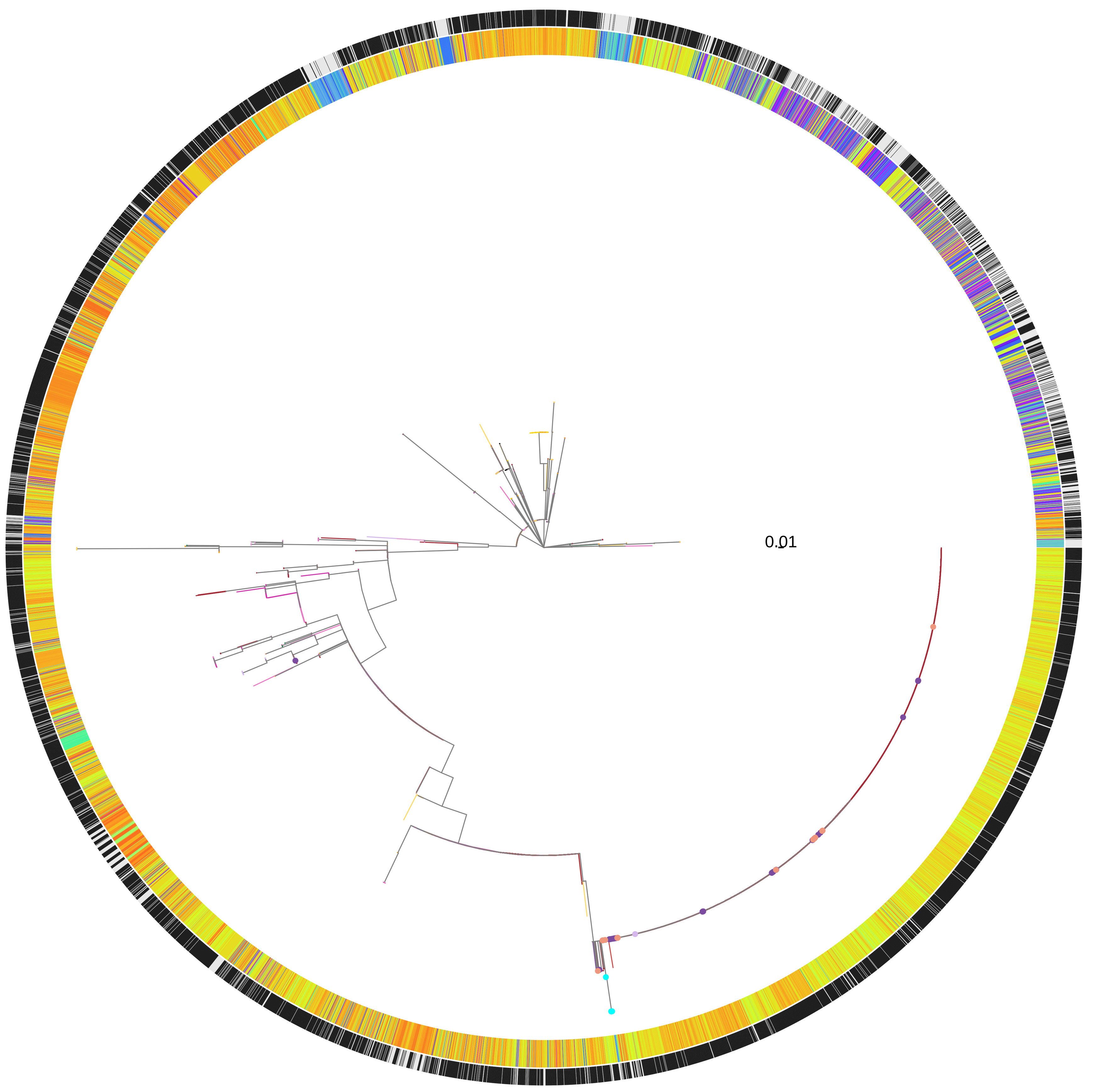


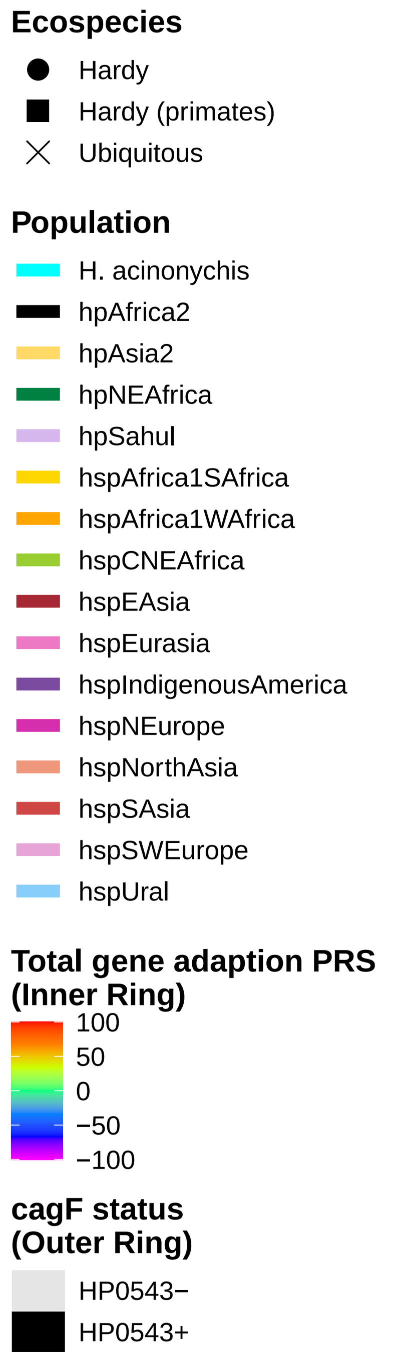

**HP0608 | HP0609**
**P−val: 48.8268**

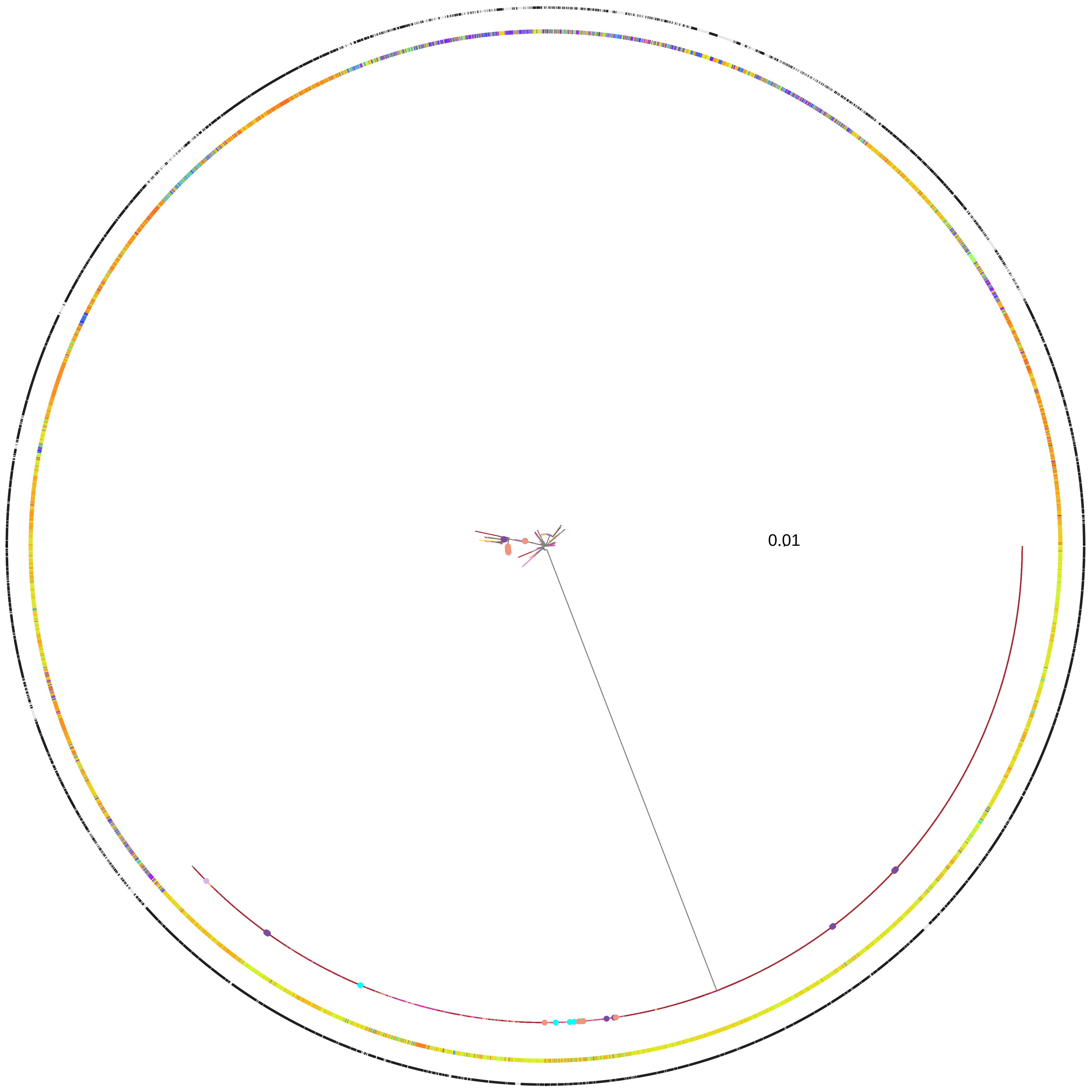


**HP0448 | HP0449**
**P−val: 42.52**

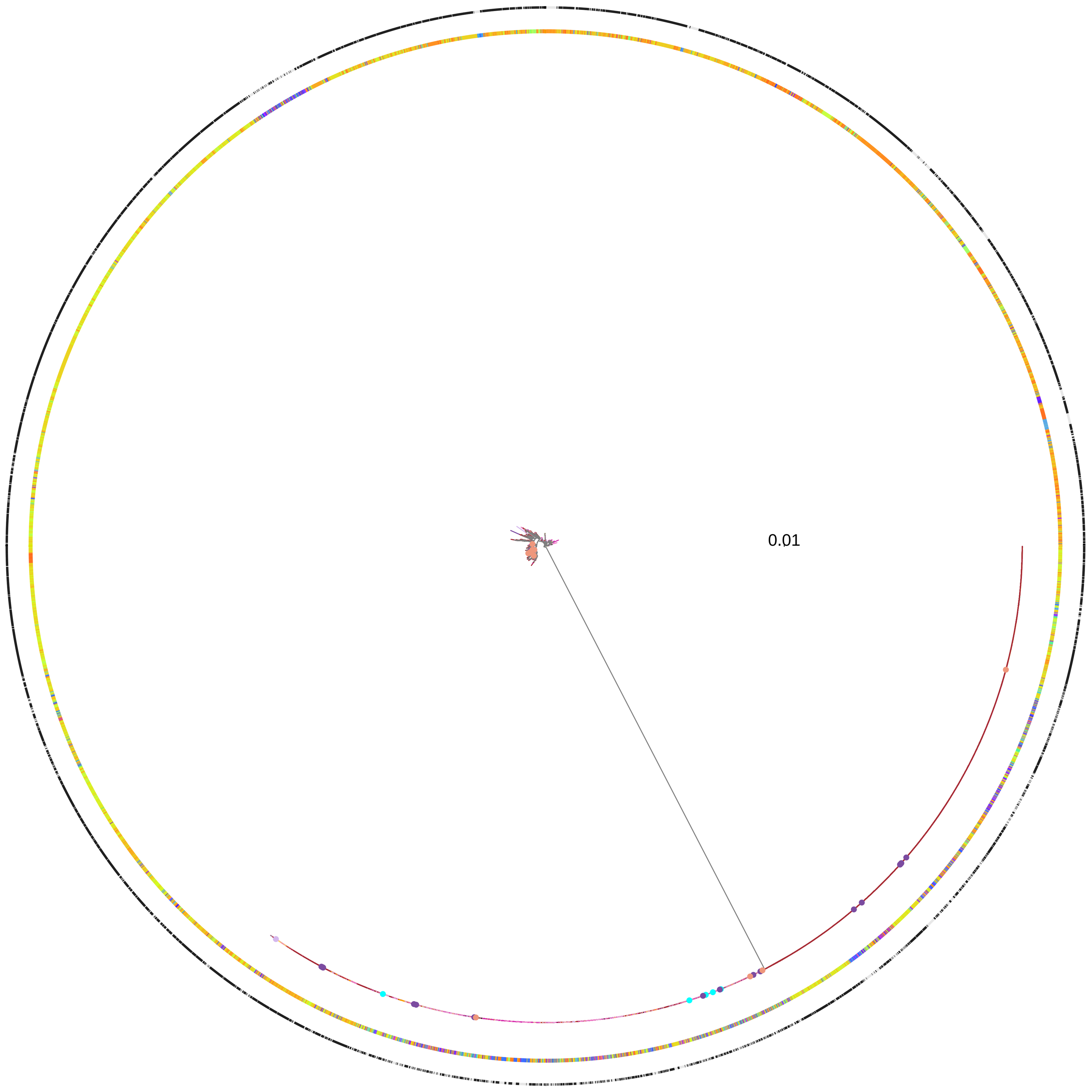


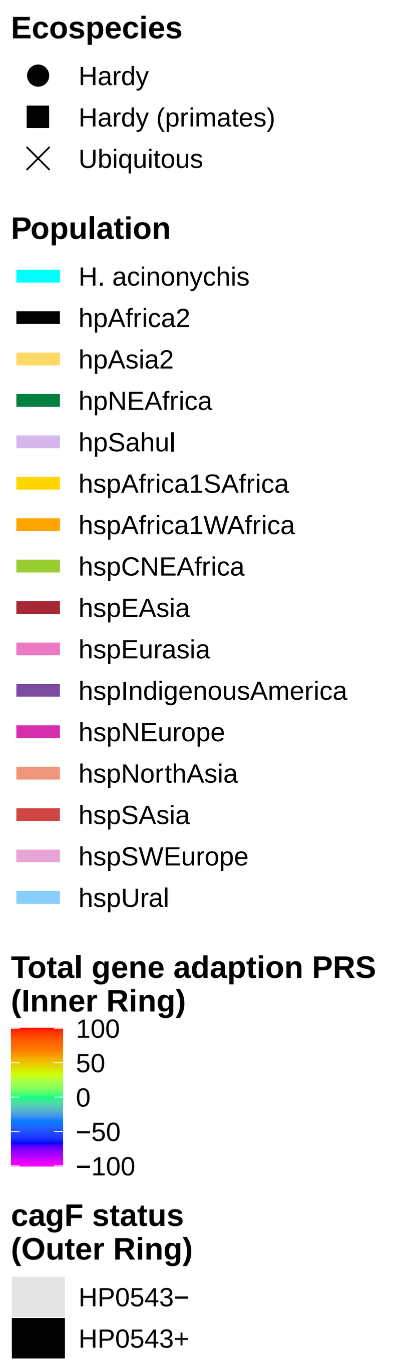
